# Identification of Ephrin type-B receptor 4 as a critical mediator of tissue fibrosis

**DOI:** 10.1101/2024.11.14.622787

**Authors:** Brian Wu, Starlee S. Lively, Shabana Vohra, Noah Fine, Chiara Pastrello, Anca Maglaviceanu, Osvaldo Espin-Garcia, Evan Pollock-Tahiri, Sayaka Nakamura, Paramvir Kaur, Keemo Delos Santos, Jason S. Rockel, Pratibha Potla, Himanshi Gupta, Poulami Datta, Laura Tang, Jacob Kwon, Akihiro Nakamura, Matthew B. Buechler, Rajiv Gandhi, Jiangping Wu, Boris Hinz, Igor Jurisica, Mohit Kapoor

## Abstract

Pulmonary fibrosis (PF) is a pathology associated with interstitial lung diseases (ILDs), including idiopathic pulmonary fibrosis (IPF). Fibrosis promotes continual secretion of extracellular matrix (ECM), producing non-functional scar tissue and causing organ failure. This study investigated the tyrosine kinase receptor Ephrin type-B receptor 4 (EphB4) as a mediator of PF. To this end, we generated mice with conditional *Col1a2-*driven deletion of *Ephb4* and used a preclinical mouse model of PF, total and single nuclei RNA (snRNA) sequencing, NanoString, previously published single cell data, computational analysis, and functional assays of mouse and human healthy control and IPF lung fibroblasts. *Col1a2*-Cre*^ERT^*-driven *Ephb4* deletion, or EphB4 inhibition via NVP-BHG712, markedly protected against bleomycin-induced PF. Total RNA-sequencing of fibroblasts isolated from *Ephb4*-deficient fibrotic mouse lungs exhibited reduced expression of ECM, ER Cargo, and protein trafficking-related genes. NVP-BHG712 reduced expression of these identified genes in mouse lung fibroblasts under fibrotic conditions *in vitro*. SnRNA sequencing of mouse lungs treated with NVP-BHG712 identified transcriptomic changes of ECM genes in specific fibroblast subpopulations. RNA sequencing, computational, and functional assays using mouse and human IPF fibroblasts identified elastin as a key mediator involved in EphB4 signaling. Combined, our data show that EphB4 is a crucial mediator of PF.

## INTRODUCTION

Pathological overactivation of tissue repair facilitates fibrosis, causing irreversible tissue damage which can lead to organ failure and patient mortality (1). Pulmonary fibrosis (PF) is characterized by excessive extracellular matrix (ECM) deposition that severely impairs oxygen exchange and deteriorates health. Idiopathic pulmonary fibrosis (IPF) and systemic sclerosis-related interstitial lung disease (SSc-ILD) are two diseases where PF is a primary pathology causing patient mortality (2). Advancements in PF research have led to the approval of growth factor receptor inhibitors nintedanib and pirfenidone for the treatment of IPF, along with the subsequent repurposing of nintedanib for the treatment of progressing fibrotic ILDs(3–5). Both nintedanib and pirfenidone inhibit multiple receptors in epidermal and connective tissue cells (‘fibroblasts’), which has accelerated the investigation of epithelial and connective tissue in fibrogenesis (3–5). Despite significant therapeutic advancements, current interventions neither halt nor reverse PF. Deciphering disease mechanisms underlying fibrosis will help uncover novel therapeutic avenues to improve patient outcomes.

Erythropoietin-producing hepatocellular receptor interacting protein (Ephrin) and Eph receptors are unique protein families that undergo bidirectional signaling, where ligand-receptor interactions initiate downstream signaling from both the receptor (forward signaling) and ligand (reverse signaling)(6). The Ephrin-B2 ligand and EphB4 receptor signaling axis mediates angiogenesis and vasculogenesis in prenatal development by regulating cell migration via actin cytoskeletal remodeling pathways (6). Further evidence suggests that Ephrin-B2 may play a key role in pulmonary, cardiac, skin, and hepatic fibrosis (7–9). Release of the soluble Ephrin-B2 active ectodomain into lung tissue, modulated by a disintegrin and metalloproteinase domain-containing protein 10 (ADAM10), was shown to promote fibrosis (7). Further, siRNA targeting of EphB4 abrogates soluble Ephrin-B2-mediated fibrotic responses in pulmonary fibroblasts *in vitro* (7). While EphB4 was proposed as a target of Ephrin-B2-mediated pro-fibrotic responses *in vitro*, to the best of our knowledge, the exact role of EphB4 signaling in PF is yet to be investigated.

To study the role of EphB4 receptor in PF, we used a preclinical PF mouse model, total and snRNA sequencing, NanoString, previously published single cell data, integrative computational biology analysis, and functional assays using primary mouse and human lung fibroblasts (**Figure 1**). We generated mice with conditional deletion of *Ephb4* in *Col1a2*-expressing cells, including fibroblasts, to show that *Ephb4* deletion reduced the severity of bleomycin-induced PF. This protective antifibrotic phenotype was replicated using the pharmacological EphB4 inhibitor NVP-BHG712. High-throughput RNA sequencing of lung fibroblasts derived from bleomycin-treated mice showed that *Ephb4* deletion induced downregulation of key genes associated with ECM regulation, protein trafficking and endoplasmic reticulum (ER) cargo concentration. NanoString analysis on mouse lung fibroblasts treated with EphB4 inhibitor also indicated a reduction in the expression of key ECM, protein trafficking and ER cargo genes. SnRNA sequencing identified 4 fibroblast subclusters in the lungs of bleomycin-treated mice. Elastin (*ELN*) expression was higher in identified fibroblast subclusters 1 and 2, and *ELN* expression was reduced with EphB4 inhibition under bleomycin condition. Total RNA sequencing identified *ELN* as the most differentially expressed ECM gene in primary human lung fibroblasts isolated from IPF patients, which also exhibited significant downregulation in response to EphB4 inhibition. Integrated computational analysis of mouse and human fibroblast total RNA sequencing data identified *ELN* as a crucial EphB4 signaling target. Transforming growth factor-β1 (TGFβ1)- induced ELN expression was reduced by inhibition or silencing of EphB4 in human IPF fibroblasts. Similarly, ELN was reduced in bleomycin-treated mouse lungs following pharmacological inhibition or genetic *Ephb4* deletion. This study suggests what we believe to be a novel mechanism by which EphB4 signaling contributes to PF.

**Figure 1.**
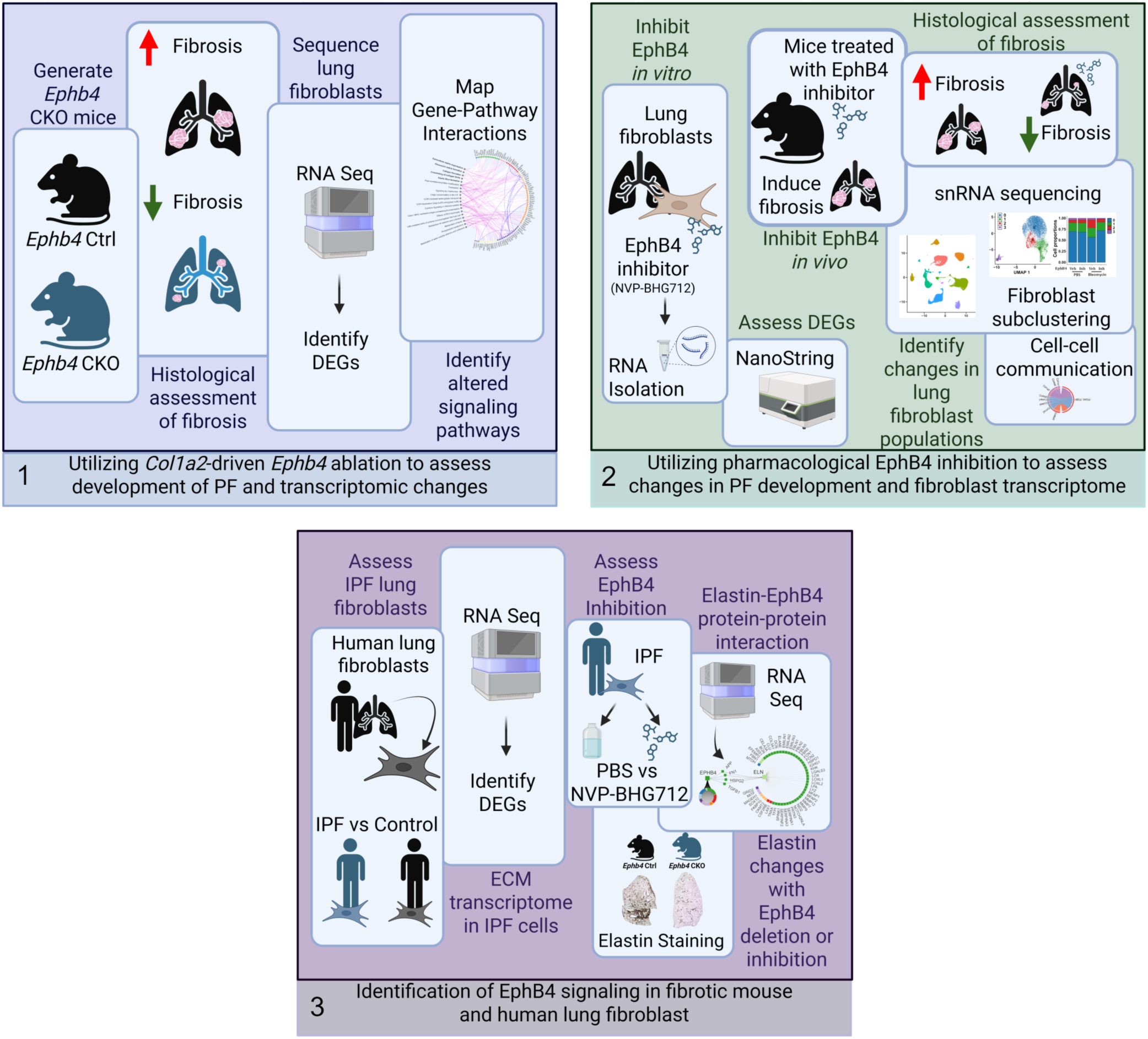
Methodological approaches used to assess the role of EphB4 in pulmonary fibrosis. A schematic representation of the methods and experimental approach used in this study.

## RESULTS

### Genetic deletion of *Ephb4* in *Col1a2*-expressing cells: generation of *Ephb4* conditional knockout mice

As part of the pathology of PF, fibroblasts are responsible for the deposition of excessive ECM components, like type I collagen. To examine whether *Ephb4* expression in *Col1a2-*expressing cells (such as fibroblasts) contributes to PF development, floxed *Ephb4* (*Ephb4^tm1.1Jwu^*) mice were crossed with B6.Cg-Tg(*Col1a2*-Cre/ERT,-ALPP)7Cpd mice to generate *Ephb4^f/f^*;*Col1a2-Cre^ERT^* mice (detailed pedigree for the generation of the mouse line is included in **Supplementary Figure 1**). *Col1a2-*driven expression of Cre in mice has been used in previous studies to ablate specific genes in fibroblasts(7, 10, 11). PCR-genotyping was performed to confirm the presence of *Col1a2*-*Cre^ERT^*in mice exhibiting homozygosity of the floxed *Ephb4* sites (500 bp) compared to the WT *Ephb4* gene (400 bp) (**Figure 2A**). At 4-5 weeks of age, *Ephb4^f/f^;Col1a2-Cre^ERT^*mice were treated with 4-hydroxytamoxifen (1 mg/day for 5 days) to induce *Ephb4* gene deletion in *Col1a2*-expressing cells, including fibroblasts, or with corn oil (vehicle). Sustained knockdown of *Ephb4* expression was confirmed at both mRNA and protein levels in primary lung fibroblasts isolated 7 days after the final 4-hydroxytamoxifen administration (**Figure 2B**). No significant reduction in the expression of EphB4 was observed in other cell types examined, such as macrophages and endothelial cells (**Supplementary Figure 2A/B**). Additionally, cell culturing of primary lung fibroblasts to passage 1 (P1) yielded high purity of fibroblast populations (**Supplementary Figure 2C/D**). Thus, *Ephb4* conditional knockout mice permit 4-hydroxytamoxifen-inducible deletion of *Ephb4* in *Col1a2-*expressing cells (herein referred to as *Ephb4* CKO mice), including fibroblasts, compared to control mice treated with corn oil (*Ephb4* control mice).

**Figure 2.**
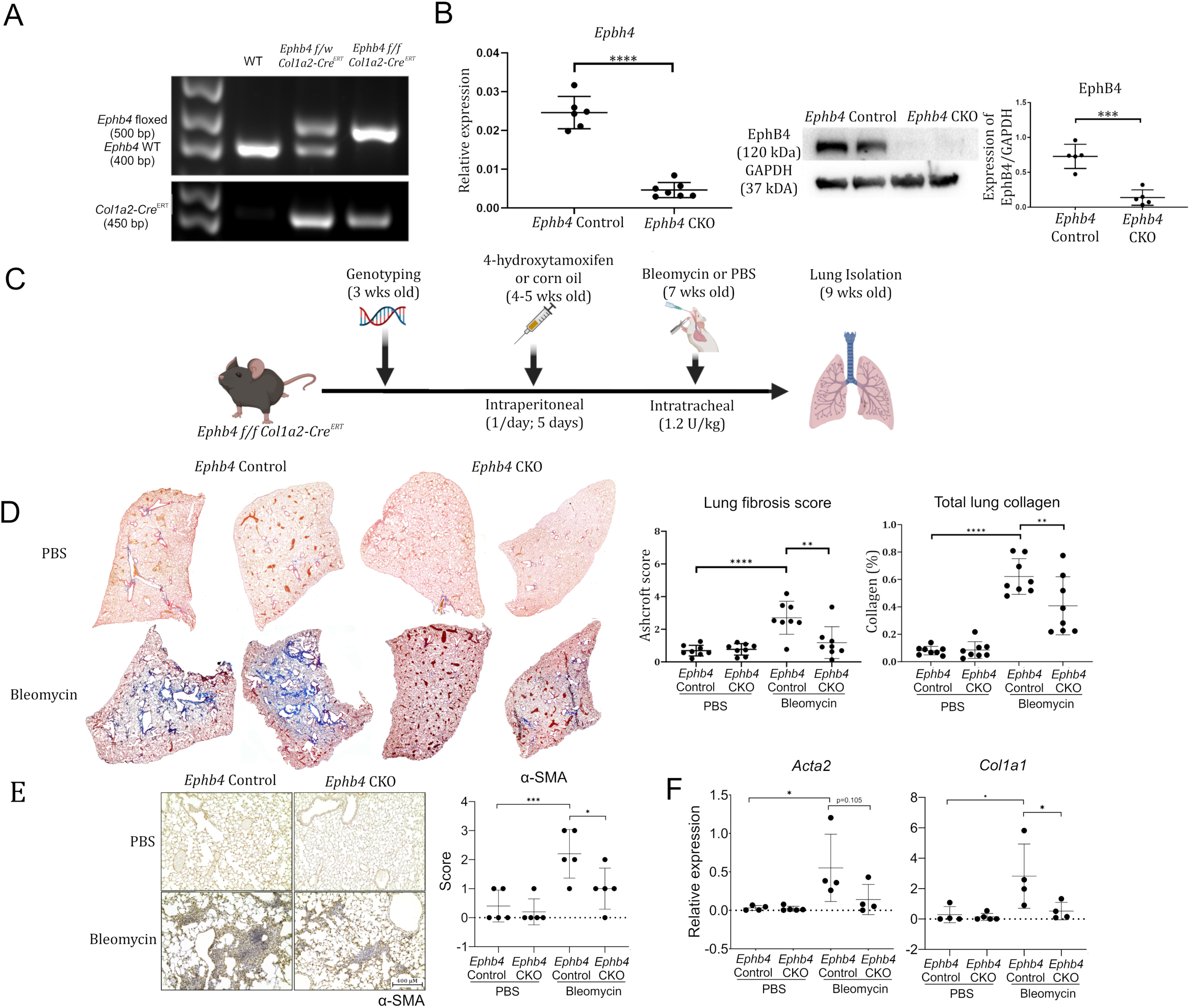
*Ephb4* conditional knockout mice exhibit a protective phenotype against the development of pulmonary fibrosis. *Ephb4^f/f^* mice were crossed with *Col1a2-Cre^ERT^* mice to generate a line with conditional *Col1a2*-driven deletion of *Ephb4* (*Ephb4* CKO). (A) Genotyping was performed to confirm the presence of homozygous floxed *Ephb4* alleles and *Cre*. (B) Mice treated with 4-hydroxytamoxifen to induce the CKO were sacrificed and primary lung fibroblasts were cultured to first passage. RNA was collected and *Ephb4* was quantified by qRT-PCR (n=6–7). Protein lysates were collected from cell cultures and EphB4 protein was quantified by Western blotting (n=5). (C) A mouse model for inducing fibrosis in *Ephb4* CKO mice was performed using intratracheal bleomycin challenge or PBS (control) following 4-hydroxytamoxifen-induced deletion. (D) Mouse lungs were collected at end point (9 weeks of age) and stained using Gomori’s One Step collagen stain (blue). Representative histology images from two separate animals/group are shown. Fibrosis was quantified either by modified Ashcroft scoring or with ImageJ total lung collagen content quantification (n=8 mice/group). (E) αSMA immunostaining was performed in lung tissue sections to assess fibrotic activity in *Ephb4* CKO and control mice with and without bleomycin challenge (n=5). Semi-quantitative scoring (0–3) of the staining intensity was assigned to each mouse lung. (F) Primary lung fibroblasts were isolated from *Ephb4 Col1a2-Cre^ERT^* mice treated with 4-hydroxytamoxifen or corn oil, then bleomycin. RNA was isolated from fibroblasts and qRT-PCR was performed to detect *Col1a1* and *Acta2* (n=4). For qRT-PCR (B, F), 2^-ΔCT^ values were used to represent relative expression of the gene of interest normalized to *Gapdh*. For all graphs, data are expressed as mean ± SD where differences (*, p < 0.05; **, p < 0.01; ***, p < 0.001; ****, p<0.0001) were identified using either Student’s t-test (B) or 2-way ANOVA with Tukey’s post-hoc tests (D-F).

### *Ephb4* CKO mice are protected against the development of PF

*Ephb4* CKO and *Ephb4* control mice were subjected to intratracheal instillation of bleomycin to induce PF, with PBS used as a corresponding bleomycin control (**Figure 2C**). Mouse lungs were collected 14 days later to assess collagen content using Gomori’s One Step Trichrome blue collagen stain. As shown in **Figure 2D**, collagen content was comparable in both *Ephb4* CKO and *Ephb4* control lungs treated with PBS. After bleomycin challenge, *Ephb4*-expressing control mice showed a striking increase in collagen staining, indicating development of PF. Interestingly, this effect was not readily observed in *Ephb4* CKO mice treated with bleomycin. Instead, deletion of *Ephb4* resulted in a marked decrease in PF severity and reduced the number of lobes exhibiting fibrosis, as determined by Ashcroft scoring and collagen staining quantification (**Figure 2D; Supplementary Table 1**) (12). Next, immunohistochemical staining of alpha smooth muscle actin (αSMA, gene name *Acta2*), a marker of activated fibroblasts associated with fibrosis (13), was examined. As expected, PBS control treatment yielded minimal αSMA staining in mouse lungs of *Ephb4* CKO and control mice. Bleomycin treatment induced strong αSMA staining in *Ephb4* control mice with substantially weaker staining in *Ephb4* CKO mice (**Figure 2E**). Further, periostin staining was also reduced in the lungs of *Ephb4* CKO compared to control mice, indicating a protective phenotype in mouse lungs with *Ephb4* ablation (**Supplementary Figure 3A**). Intriguingly, the antifibrotic phenomenon of *Ephb4* deletion in lung fibroblasts after bleomycin challenge was maintained *in vitro*. Primary lung fibroblasts derived from bleomycin-treated *EphB4* CKO mice also exhibited reduced αSMA *Acta2* and type I collagen (*Col1a1*) gene expression compared to bleomycin-treated *EphB4* control mice (**Figure 2F**). Taken together, it appears that *Ephb4* in lung fibroblasts plays an important role in PF development.

We next wanted to determine if *Ephb4* CKO had a similarly altered response of fibroblasts in another form of fibrosis. To this aim, we examined a skin fibrosis model. We first confirmed knockdown of *Ephb4* expression at both mRNA and protein levels in skin fibroblasts from *Ephb4* CKO and *Ephb4* control mice (**Supplementary Figure 4A**). After *Ephb4* knockdown, mice received daily subcutaneous bleomycin injections (0.5 U/mL) for 21 days (**Supplementary Figure 4B)**. Skin samples were then collected 22 days after *Ephb4* knockdown and subjected to histological Gomori’s trichrome staining. No differences were observed in fibrosis severity and dermal thickening (**Supplementary Figure 4C)**. Thus, the antifibrotic effect of *Ephb4* CKO observed in PF was not observed in the bleomycin-induced model of skin fibrosis, suggesting that the effects of *Ephb4* ablation in fibroblasts may depend on the type of tissue/organ and the tissue microenvironment.

### Total RNA sequencing identifies downregulation of ECM, endoplasmic reticulum (ER) cargo concentration, and protein trafficking-related transcriptome in lung fibroblasts from *Ephb4* CKO mice

To elucidate the effects of *Ephb4* ablation on the lung fibroblast transcriptome in the context of PF, total RNA sequencing was performed on lung fibroblasts isolated from *Ephb4* CKO (or *Ephb4* control mice) 2 weeks after bleomycin (or PBS) challenge (**Figure 3A**). First, the effects of bleomycin on lung fibroblast transcriptome were examined by comparing *Ephb4* control mice (bleomycin vs. PBS) two weeks after bleomycin treatment. A total of 257 differentially expressed genes (DEGs; false discovery rate (FDR) q<0.05) were identified (155 upregulated and 102 downregulated) in bleomycin-treated *Ephb4* control mice compared to PBS-treated *Ephb4* control mice. No DEGs were identified in lung fibroblasts from *Ephb4* control and *Ephb4* CKO mice after PBS treatment. We next examined the effect of *Ephb4* ablation on bleomycin-induced gene expression in lung fibroblasts. When Ephb4 CKO and *Ephb4* control mice challenged with bleomycin were compared, 317 DEGs were identified (103 upregulated and 214 downregulated) in lung fibroblasts from bleomycin-challenged *Ephb4* CKO mice (**Figures 3B/C**), suggesting that the effects of *Ephb4* ablation in lung fibroblasts are unique to fibrotic conditions. The list of top DEGs identified can be found in **Supplementary Table 2** and the full sequencing dataset can be accessed online on the GEO data repository (GSE205612).

**Figure 3.**
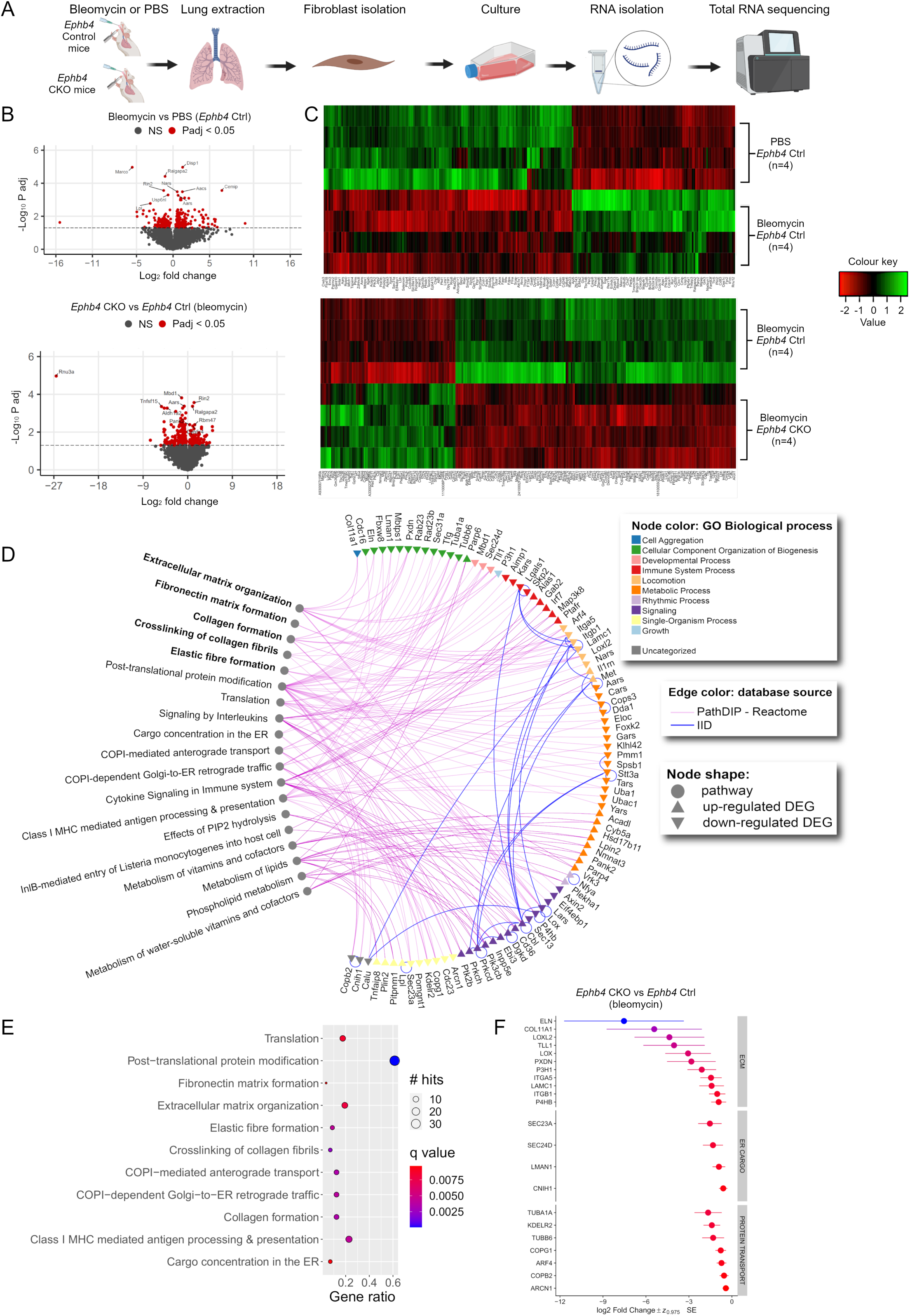
*Ephb4* CKO fibrotic mouse lung fibroblasts exhibit unique transcriptomic profiles. (A) *Ephb4^f/f^;Col1a2-Cre^ERT^* mice were treated with 4-hydroxytamoxifen (1 mg/day for 5 days) or corn oil control at 4 weeks of age, then intratracheally instilled with PBS or bleomycin (1.2U/kg) at 6 weeks of age. Mice were sacrificed 14 days after bleomycin or PBS challenge and lung fibroblasts were isolated and cultured (n=4 mice per group). Fibroblasts were enriched with a single passage followed by RNA extraction for total RNA sequencing. (B) Pairwise comparisons were made between bleomycin and PBS-treated *Ephb4* control mice, or between *Ephb4* CKO and control mice under bleomycin condition (n=4 mice/group). Differentially expressed genes (DEGs) were determined using an FDR cutoff of q<0.05 (red). In the bleomycin vs PBS comparison 257 DEGs were identified in fibroblasts, including 155 upregulated and 102 downregulated DEGs. In the *Ephb4* CKO vs control comparison, 317 DEGs were identified in fibroblasts, including 103 upregulated and 214 downregulated DEGs. (C) DEGs from both comparisons were visualized using heat maps. (D) Lists of both upregulated and downregulated DEGs in the RNA sequencing dataset of *Ephb4* CKO (n=4) vs *Ephb4* control mice (n=4) treated with bleomycin were analyzed separately using pathDIP to identify subsets of genes enriched for unique pathways in the Reactome database. Gene-pathway associations were visualized in NAViGaTOR v.3.0.16. (E) Unique gene-pathway enrichments from the downregulated gene list visualized by dotplot, with significance determined using the Benjamini and Hochberg method (with cutoff of q<0.01). (F) The expression of downregulated DEGs associated with ECM organization, protein transport and ER cargo concentration visualized by Wideplot.

Next, we used pathDIP (14) to identify pathways enriched in the up and downregulated DEG lists of *Ephb4* CKO lung fibroblasts relative to their *Ephb4* control counterparts under bleomycin challenge (**Supplementary Table 3)**. A subset of 31 upregulated DEGs were uniquely enriched for 8 pathways, and a subset of 57 downregulated DEGs were uniquely enriched for 11 pathways. Upregulated DEGs were uniquely enriched for pathways associated with phospholipid and lipid metabolism, cytokine signaling and interleukin signaling. Among the downregulated genes, 11 were associated with ECM organization, 7 with protein trafficking, 4 with ER cargo concentration, and 12 with class I mediated antigen processing and presentation. Gene set enrichment analysis (GSEA) using both Reactome and Gene Ontology Biological Process databases showed that downregulated genes were also mainly enriched for ECM and collagen organization pathways, while upregulated genes showed enrichment for innate immune response and cytokine signaling (**Supplementary Table 4**). The unique enriched gene-pathway associations identified using pathDIP (14) were visualized in an interaction map using NAViGaTOR (15) (**Figures 3D**).

The list of gene-pathway associations was used to identify cellular processes that *Ephb4* might regulate under fibrotic conditions. Unique DEGs downregulated in lung fibroblasts from *EphB4* CKO mice challenged with bleomycin were associated with ECM-relevant processes, including collagen formation and crosslinking, elastic fiber formation, fibronectin matrix formation or ECM organization as well as genes related to protein trafficking and ER cargo concentration related processes (**Figure 3E**). Interestingly, of the 11 downregulated DEGs associated with ECM organization identified, many have been documented as prominent profibrotic mediators, including: prolyl 4-hydroxylase beta polypeptide (*P4bh*), integrin b1 (*Itgb1*), integrin a5 (*Itga5*), peroxidasin (*Pxdn*), lysl oxidase like 2 (*Loxl2*), lysl oxidase (*Lox*), type XI collagen (*Col11a1*), tolloid like 1 (*Tll1*), and *Eln* (16–21). Among the ECM, ER cargo and protein trafficking genes, 20/22 exhibited a >1.5-fold change in gene expression in bleomycin-challenged *Ephb4* CKO animals compared to bleomycin-challenged *Ephb4* control mice, with *Eln* having the greatest mean fold change (**Figure 3F**).

Overall, total RNA sequencing showed that under fibrotic stimulation (bleomycin treatment), fibroblasts from *Ephb4* CKO mice compared to *Ephb4* control mice exhibit reduced expression of genes related to fibrosis, ECM organization, ECM remodeling, ER cargo concentration and protein trafficking.

### Pharmacological EphB4 inhibition reduces mouse lung fibroblast expression of genes related to ECM organization, ER cargo concentration and protein trafficking

We next examined whether pharmacological inhibition of EphB4 elicited comparable changes in gene expression (as observed in total RNA sequencing) in primary mouse lung fibroblasts. WT primary lung fibroblasts were isolated from C57BL/6J mice and stimulated with TGFβ1 (or PBS as control) in combination with the EphB4 small molecule inhibitor NVP-BHG712 (or vehicle) (**Figure 4A**). NVP-BHG712 has previously been shown to decrease EphB4 activity and phosphorylation *in vitro* and *in vivo,* including in mouse lung tissue(22). We first confirmed that NVP-BHG712 effectively inhibits phosphorylation of EphB4 in human lung fibroblasts at 250 nM (**Supplementary Figure 5**) and used this concentration for subsequent experiments. A NanoString CodeSet of 47 gene targets was designed using our previously identified downregulated DEGs involved in ECM organization, protein trafficking, ER cargo concentration, MHC Class I presentation and post-translational protein modification (remaining genes were excluded, including those related to translation). TGFβ1 significantly increased the expression of 25 of the 47 selected genes, with EphB4 inhibition significantly reducing the expression of 19 out of 25 of these genes (see **Supplementary Table 5** for the full dataset). Notably, EphB4 inhibition downregulated many genes associated with cellular processes involved in ECM organization (9/11), ER cargo concentration (3/4), and protein trafficking (6/7) (**Figure 4B**, see **Supplementary Figure 6** for graphical representations of the remaining genes). Thus, pharmacological inhibition of EphB4 in mouse lung fibroblasts recapitulated reduced expression of most genes identified in our *in vivo Ephb4* CKO mouse lung fibrosis model.

**Figure 4.**
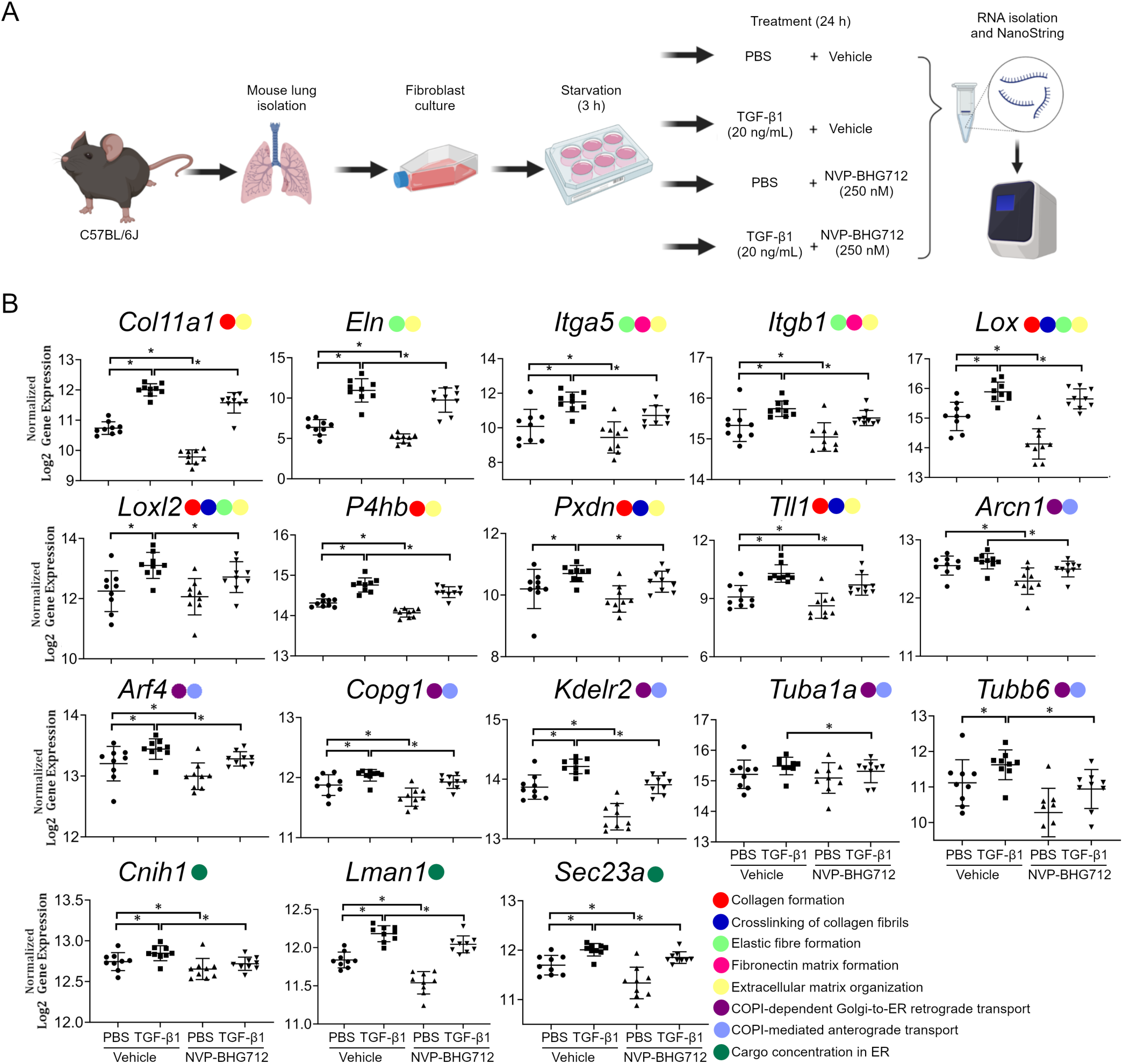
NVP-BHG712 downregulates TGFβ1-induced expression of genes related to ECM organization, protein trafficking, and ER cargo concentration regulation. (A) Lungs were isolated from WT mice and fibroblasts were cultured. Fibroblasts were then simultaneously treated with the pro-fibrotic cytokine TGF-β (20 ng/mL; or PBS as control) and the EphB4 inhibitor NVP-BHG712 (250 nM; or DMSO as inhibitor control) for 24 h (n=9). RNA was isolated and NanoString was performed on a 47 gene CodeSet to examine the expression of downregulated DEGs identified by RNA-seq in bleomycin-challenged *Ephb4* CKO lung fibroblasts. (B) Transcript levels of a subset of the CodeSet of genes that were significantly upregulated with TGFβ1 treatment and downregulated with NVP-BHG712 treatment and were associated with ECM organization, protein trafficking, and ER cargo concentration regulation (remaining genes can be found in **Supplementary Figure 6**). Gene expression was normalized to housekeeping genes *Rplp0* and *Acad9* and displayed as log2 transformed gene expression (mean ± SD). Wilcoxon signed rank test with false discovery rate (FDR) q≤0.05 was used to identify significant changes in gene expression. Legend colours indicate the gene-associated pathDIP-identified pathways. *, q-value < 0.05.

### NVP-BHG712 attenuates bleomycin-induced PF in mice

Since NVP-BHG712 attenuated the expression of key genes involved in ECM organization, ER cargo concentration and protein trafficking in cultured mouse lung fibroblasts, we next tested its anti-fibrotic potential *in vivo* using the bleomycin lung fibrosis mouse model. 7-week-old WT C57BL/6J mice were intratracheally challenged with bleomycin or PBS (control) and 5 days later received NVP-BHG712 (10 μg/kg daily, *p.o.*) or vehicle (control; N-methyl-2-pyrrolidone plus polyethylene glycol 300 at ratio of 1:9) for 5 days. Lungs were harvested for Gomori’s blue collagen stain histological assessment 14 days after the start of the bleomycin challenge (**Figure 5A**). As observed in *Ephb4* CKO mice, WT mice treated with NVP-BHG712 also exhibited reduced severity in the degree of bleomycin-induced lung fibrosis (**Figure 5B**). Ashcroft lung fibrosis score of affected areas showed reduced fibrosis severity in NVP-BHG712-treated mice, with a mean ± SD disease severity score of 2.48±0.55 compared to 4.05±0.61 in bleomycin challenged, vehicle-treated lungs (**Figure 5C, top**). Histomorphometric analysis of lung section images indicated that lung collagen content in NVP-BHG712 treated lungs decreased from 49.7±8.8% to 37.81±6.8% (mean ± SD) per NVP-BHG712-treated animal (**Figure 5C, bottom**). Notably, there was no significant change in Ashcroft scoring or lung collagen quantification in NVP-BHG712 treated mice compared to vehicle treated mice in the PBS-challenged groups. αSMA and periostin staining was also markedly reduced in NVP-BHG712-treated lungs compared to vehicle-treated controls after bleomycin challenge, with no difference found between PBS-treated controls (**Figure 5D; Supplementary Figure 3B**). Collectively, these results show that EphB4 pharmacological inhibition in mice confers protection in bleomycin-induced lung fibrosis, similar to the genetic deletion in *Ephb4* CKO mice.

**Figure 5.**
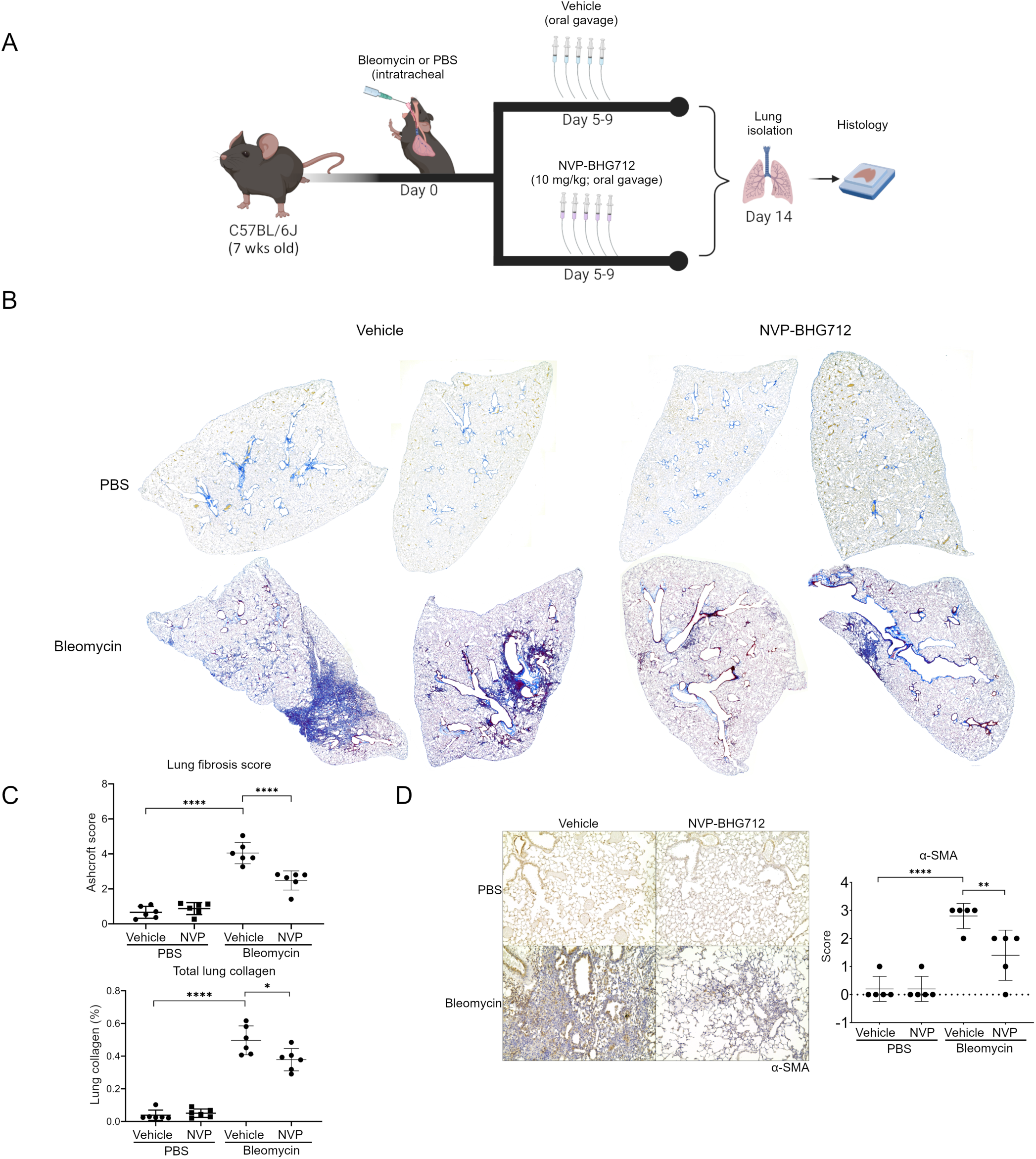
Pharmacological inhibition of EphB4 protects against development of PF in mice. (A) Bleomycin sulphate (1.2 U/kg) or PBS (control) was intratracheally administered to WT C57Bl/6J mice. Five days later, mice were given either NVP-BHG712 (10 μg/kg; p.o.) or vehicle (control) daily for 5 days (n=6/group). Lungs were harvested 14 days post-bleomycin challenge and histology was performed. (B) To assess fibrosis, collagen fibers in mouse lungs were stained via Gomori’s One Step Trichrome (blue). Representative histology images from 2 separate animals/group are shown. (C) Images of mouse lung sections (n=5 animals/group; 5 lobes/animal) were graded for severity of fibrosis using the Ashcroft scoring system (top graph). Images of lung sections underwent color deconvolution in Fiji V. 2.9.0 to isolate the blue channel as a representative of total lung collagen. Total lung collagen was normalized to total lung tissue and expressed as percent total lung collagen. Graphical representations indicate averages of 5 lobes per animal (bottom graph). (D) αSMA immunohistochemistry performed on mouse lung sections (n=5 animals/group). Semi-quantitative scores were assigned scores from 0–3 based on degree of staining. For all graphs, data expressed as mean ± SD and analyzed by 2-way ANOVA followed by Tukey’s post-hoc tests for multiple comparisons. *, p<0.05; **, p<0.01; ****, p<0.0001.

### Transcriptomic differences in fibroblast subclusters using snRNA sequencing of mouse lungs treated with NVP-BHG712 in the bleomycin model

Since NVP-BHG712 treatment resulted in reduced PF severity *in vivo* and reduced expression of ECM genes in mouse lung fibroblasts *in vitro*, we next conducted snRNA sequencing and bioinformatic analyses on bleomycin or PBS instilled mouse lungs treated with NVP-BHG712 or vehicle to identify lung fibroblast subsets and interrogate the differences in the ECM transcriptome of these fibroblast subsets in response to EphB4 inhibition. A total of 12 mouse lungs (n=3 PBS + vehicle, n=3 PBS+NVP-BHG712, n=3 bleomycin + vehicle and n=3 bleomycin+NVP-BHG712) were subjected to snRNA sequencing. 49,163 nuclei were sequenced and, after filtering, 39,611 nuclei were analyzed. Multiple cell types, including fibroblasts, macrophages, endothelial cells, type I alveolar cells, and type II alveolar cells, were identified based on canonical cell markers (**Figure 6A/B**). We next subclustered fibroblasts across all 12 samples, identifying 4 fibroblast subclusters (**Figure 6C**), which were present in all treatment conditions (**Supplementary Figure 7A**). A list of genes (q <0.05 and log2FC>0.5) associated with each fibroblast subcluster can be found in **Supplementary Table 6**. Interestingly, the cellular proportion of fibroblast subcluster 2 increased with the treatment of bleomycin whereas the proportion of subcluster 0 fibroblasts was decreased with bleomycin treatment. These changes in fibroblast subcluster proportions were ameliorated with NVP-BHG712 treatment. To better understand the cellular annotations of the four identified fibroblast subclusters in our dataset, we mapped our data to the markers identified in the previously published single cell studies(23, 24). Using the markers identified in these studies, we mapped the key transcriptomic profiles to our four fibroblast subclusters. Our analysis suggests that fibroblast subcluster 0 overlaps with the transcriptomic profiles associated with alveolar fibroblasts, subcluster 1 with adventitial fibroblasts, subcluster 2 with fibrotic fibroblasts, and subcluster 3 with peribronchial fibroblasts (**Figure 6D**). Furthermore, several genes previously reported to be associated with activated fibroblasts (25) mapped to fibroblast subcluster 2 in our dataset (**Supplementary Figure 7B**).

**Figure 6.**
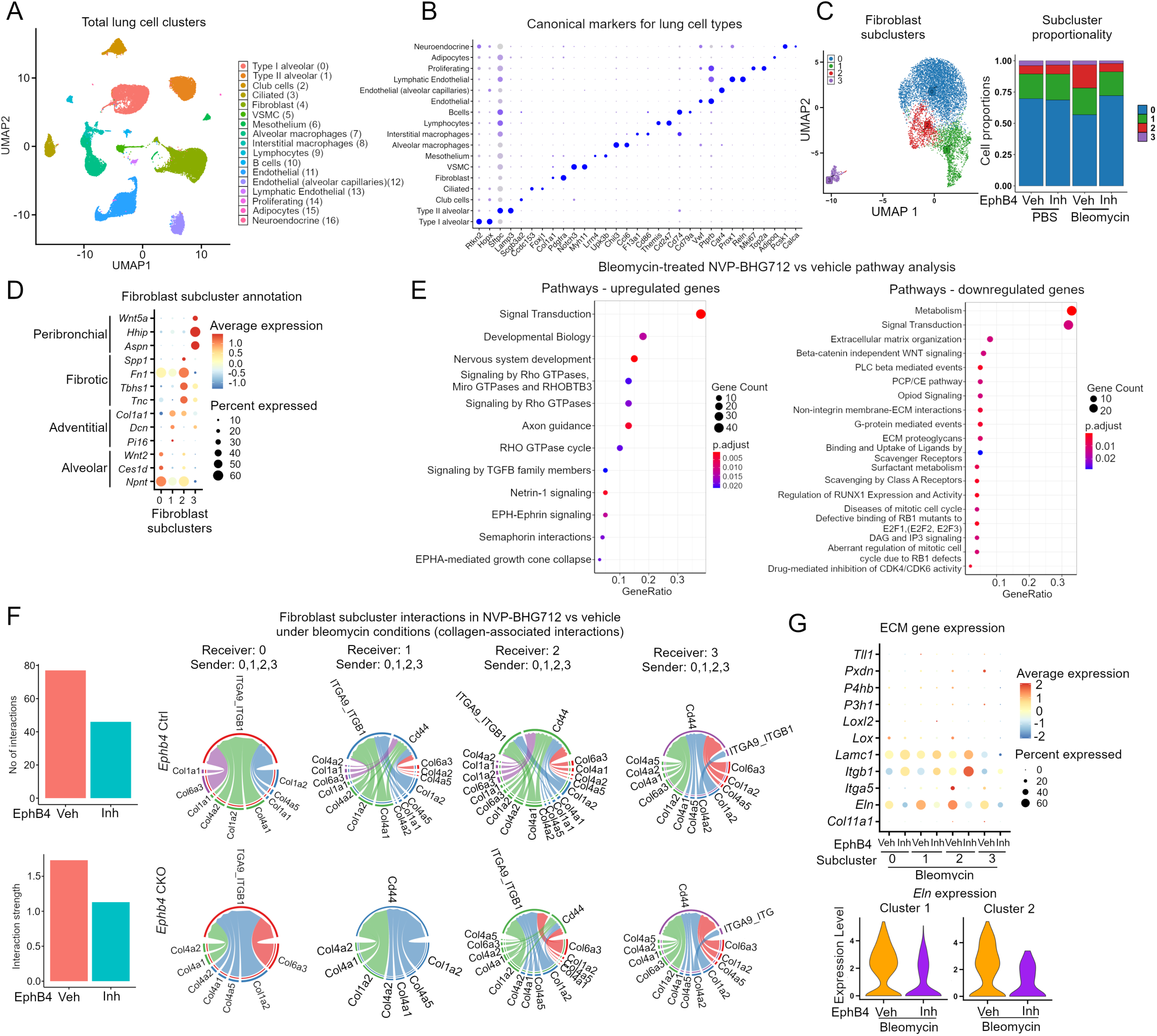
snRNA sequencing profiling of mouse lungs. (A) UMAP projection of distinct cell populations identified in mouse lung tissue (B) Dotplot showing expression of canonical markers used to annotate the identified cell clusters. The identity of each cell cluster was assigned by matching the cluster expression profile with established cell-specific marker genes. Dot color represents average expression and dot size represents the fraction of expressing cells. (C) UMAP projection of fibroblast subclusters from all treatment groups and the proportion of cells that contributed to each cluster by treatment group (veh, vehicle; Inh, NVP-BHG712). (D) Dotplot showing the expression of marker genes used to annotate the fibroblast sub- clusters(23–25) (E) Pathway analysis of upregulated and downregulated genes in bleomycin mice treated with NVP-BHG712 compared to vehicle. (F) Bar plots show total number of interactions and strength of interactions in fibroblasts in collagen signaling pathways in bleomycin mice treated with NVP-BHG712 (Inh) compared to vehicle; left panel). Circle plots showing cell-cell communication between fibroblast subclusters associated with collagen signaling in bleomycin mice treated with vehicle or NVP-BHG712. The plot shows interaction between ligands from sender and receptors from receiver. Each plot has fibroblast subcluster 0, 1, 2 and 3 as receiver and ligands from all other subclusters as senders. (G) Differential expression of 11 ECM genes in bleomycin-treated groups in all four fibroblast subclusters in NVP-BHG712 compared to vehicle (left panel). Violin plots showing the expression of *Eln* in fibroblast subclusters 1 and 2 (right panel).

Next, we performed pathway enrichment analysis using pathDIP (14) on up-and down-regulated genes in response to EphB4 inhibition under bleomycin condition. Results showed that both up- and down-regulated genes were enriched with multiple pathways (**Figure 6E**). Notably, down-regulated genes were associated with ECM-associated pathways, consistent to our previous findings in this study.

CellChat(26) was next used to investigate the signaling strength and interactions between fibroblast subclusters in NVP-BHG712 compared to vehicle-treated groups under bleomycin condition. CellChat analysis identified multiple ligand-receptor interactions across distinct fibroblast subclusters (**Supplementary Fig 8A**). Interestingly, collagen-based interactions exhibited one of the highest interaction strengths (**Supplementary Figure 8A/B**). While investigating collagen-based interactions specifically in NVP-BHG712 versus vehicle-treated groups under bleomycin condition, the total strength and number of interactions were reduced in response to EphB4 inhibition (**Figure 6F**)

We then evaluated the expression of all identified downregulated ECM-related genes in each fibroblast sub-cluster in response to NVP-BHG712 treatment under bleomycin condition. Interestingly, *ELN* expression was identified in all fibroblast subclusters, with subcluster 1 and 2 showing the highest expression. In response to NVP-BHG712 treatment, the expression of *ELN* was reduced (**Figure 6G)**.

Overall, our data have identified ECM transcriptomic changes associated with distinct fibroblast subsets in response to NVP-BHG712 treatment during fibrosis.

### Transcriptomic differences in control donor vs IPF human lung fibroblasts treated with NVP-BHG712: identifying ELN as an EphB4 signaling partner

To investigate the relevance of EphB4 inhibition in human fibroblasts, we treated primary human lung fibroblasts from control donors (HLF) and IPF subjects (IPF) (**Supplementary Table 7)** with NVP-BHG712 or vehicle for 24h and subjected each sample to total RNA sequencing (total 16 samples; n=4/group; **Figure 7A**). When IPF and HLF fibroblasts were compared under vehicle conditions, 3,108 DEGs (1,569 downregulated and 1,539 upregulated) were identified in the IPF group (**Figure 7B**). It is notable that the expression of *COL1A1* was significantly increased; and *ELN,* was determined as the most upregulated ECM gene.

**Figure 7.**
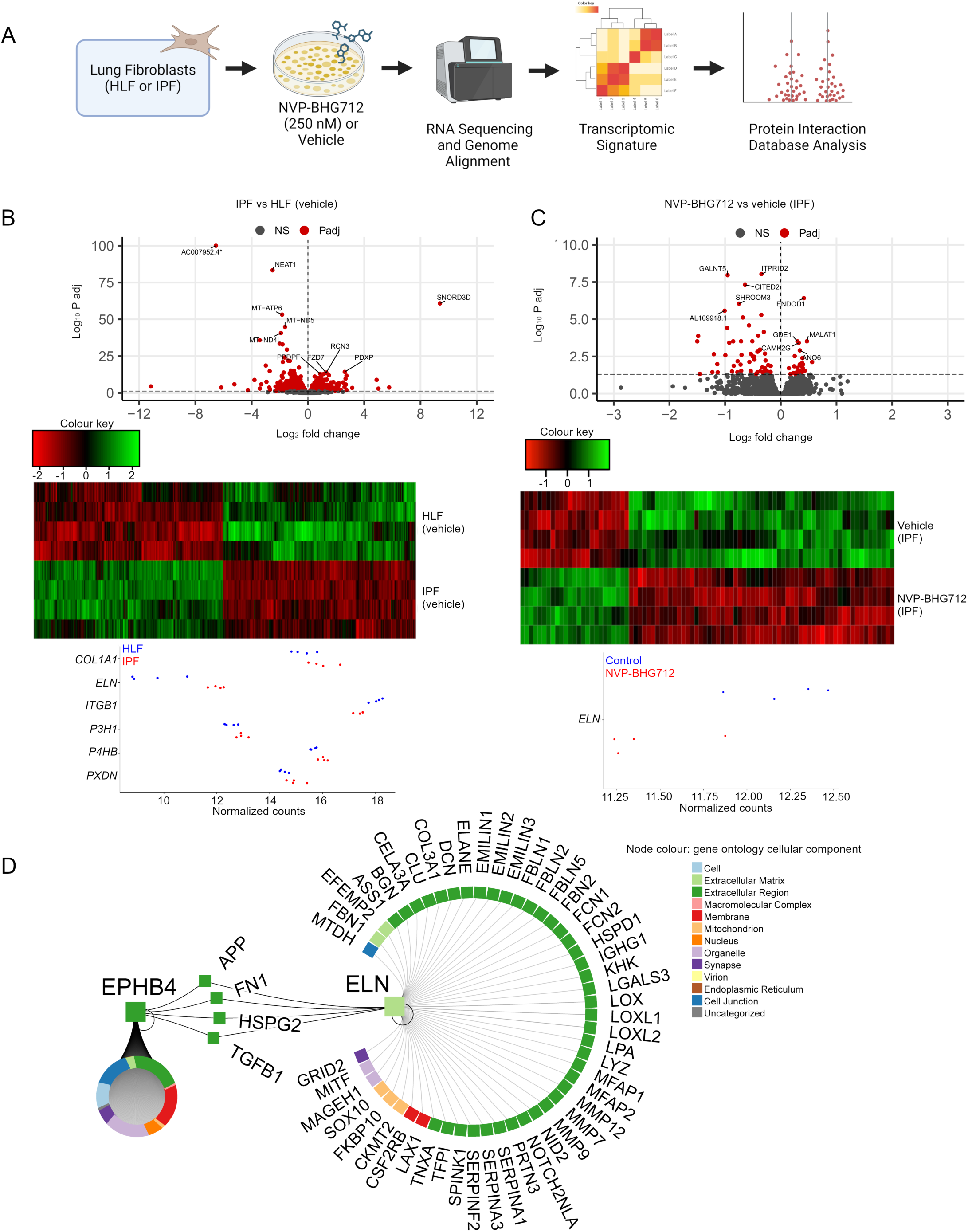
ELN is a key signaling partner of EphB4 in human IPF. (A) Schematic detailing total RNA sequencing pipeline of human lung fibroblasts (HLF) from control donors or IPF patients subjected to NVP-BHG712 (250 nM) or vehicle (n=4/group). (B) Differentially expressed genes (DEGs) comparing IPF to HLF fibroblasts using an FDR q<0.05 (red). Heat map depicting 3,108 DEGs (1,569 downregulated and 1,539 upregulated) identified in IPF vs HLF fibroblasts. Dotplot showing expression of top ECM genes in IPF vs HLF fibroblasts per donor. (C) DEGs comparing IPF fibroblasts treated with NVP-BHG712 or vehicle (red). Heat map depicting 85 DEGs (61 downregulated and 24 upregulated) identified in response to EphB4 inhibition of human IPF fibroblasts. Dotplot showing expression of *ELN* comparing NVP-BHG712 vs vehicle treatment in IPF fibroblasts per donor. (D) EphB4 was queried with ELN, as identified across human and mouse RNA sequencing, using the Integrated Interaction Database to find direct human PPIs and visualized using NAViGaTOR.

Next, the transcriptomic differences elicited by EphB4 inhibition were determined by comparing IPF fibroblasts treated with NVP-BHG712 compared to corresponding vehicle controls. In total, 85 DEGs (24 upregulated and 61 downregulated) were identified (**Figure 7C**; a list of the top 50 DEGs is provided in **Supplementary Table 8**). Notably, expression of *ELN* was significantly downregulated by NVP-BHG712 inhibition in IPF fibroblasts.

To identify common signaling partners involved in EphB4 signaling in mouse and human fibroblasts during lung fibrosis, the DEG list from human IPF RNA sequencing analysis was compared to the DEG list from *Ephb4* CKO mouse RNA sequencing. Four common gene targets were identified that included 2 upregulated genes [low density lipoprotein receptor (*LDLR*) and transmembrane protein 38b (*TMEM38B*)], and 2 downregulated genes [*ELN* and glycyl-TRNA synthetase 1 (*GARS1*)] **(Supplementary Table 9**). In our analysis using pathDIP database, *LDLR* and *TMEM38B* did not associate with any pathways, *ELN* associated with ECM organization and elastin synthesis pathways, and *GARS1* associated with protein translation. Among these 4 genes, *ELN* stood out as a prime target for investigating EphB4 signaling since *ELN* was consistently downregulated with *Ephb4* CKO (total RNA sequencing), and pharmacological EphB4 targeting in both mouse (snRNA sequencing and NanoString) and human IPF fibroblasts (total RNA sequencing).

To further elucidate a potential relationship between EphB4 signaling and ELN, we queried physical protein-protein interactions (PPIs) between the two entities in the Integrated Interaction Database (**Figure 7D; Supplementary Table 10)** (27). We identified 4 proteins that can physically interact with EphB4 and ELN, including amyloid precursor protein (APP), fibronectin (FN1), perlecan (HSPG2), and TGFβ1. It is notable that a key pro-fibrotic cytokine, namely TGFβ1, linked EphB4 to ELN(13). Indeed, GO analysis also indicated that *ELN*, among 16 other DEGs from the human IPF total RNA sequencing data, was enriched for TGFβ1 regulation of ECM (**Supplementary Figure 9**), and the downregulated genes in our sequencing analysis were enriched for ECM-related pathways (**Supplementary Table 11**). Overall, PPI analysis coupled with our sequencing data suggests that ELN may be a crucial signaling partner involved in EphB4 signaling during lung fibrosis.

### Inhibition of EphB4 mitigates fibrotic upregulation of ELN in lung fibroblasts and fibrotic lung tissue

To further investigate the regulatory relationship between EphB4 and ELN, and their potential link to TGFβ1 signaling, lung fibroblasts from IPF patients were subjected to TGFβ1 stimulation followed by EphB4 inhibition using NVP-BHG712 or *EPHB4* siRNA treatment. As shown in **Figure 8A&B**, TGFβ1 stimulation markedly increased the expression of ELN in lung fibroblasts, while subsequent treatment with NVP-BHG712 or *EPHB4* siRNA moderately or significantly attenuated -induced ELN expression, respectively. Stress fiber formation induced by TGFβ1, as indicated by rhodamine-phalloidin staining, and αSMA staining was also reduced by NVP-BHG712 (**Supplementary Figure 10**).

**Figure 8.**
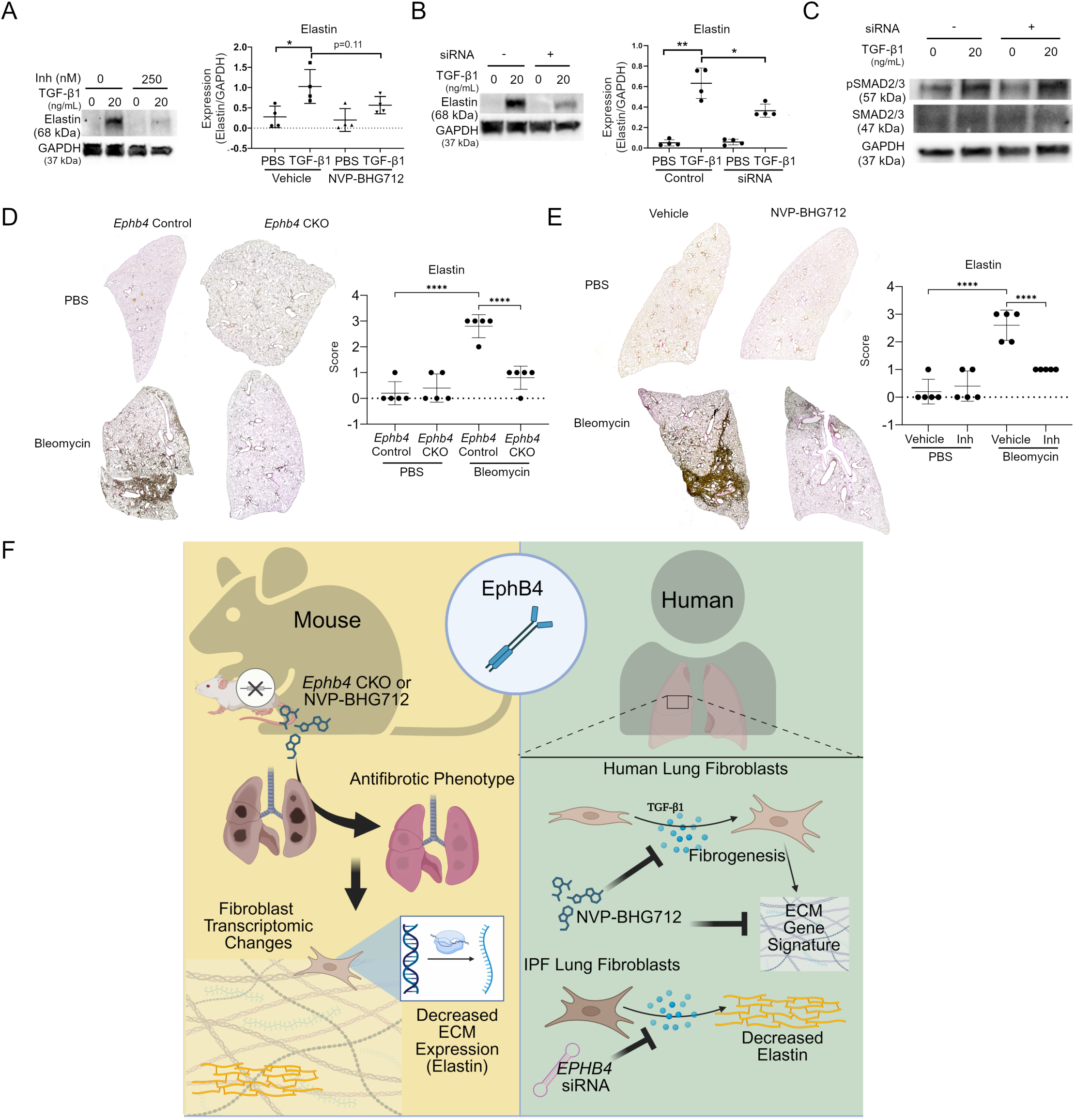
EphB4 signaling in PF. (A/B) Human IPF lung fibroblasts were treated with TGFβ1 or PBS for 24 h. Cells (total 16 samples; n=4 patients/group) were then treated with either NVP-BHG712 (250 nM) (A) or *Ephb4* siRNA (B), or their respective controls, for an additional 24 h. Protein lysates were collected and Western blotting was performed to detect ELN protein levels. ImageJ densitometry was performed and normalized to GAPDH (mean ± SD). *, p<0.05; **, p<0.01, as determined by paired two-way ANOVA followed by Tukey’s post-hoc test adjusted for multiple comparisons. (C) Western blotting of siRNA treated IPF cells to detect p-SMAD 2/3 and total SMAD 2/3 (n=4/group). (D/E) Verhoeff Van Gieson staining was performed on lung sections from (D) mice with *Ephb4* conditional knockout or respective control mice to visualize ELN (n=5/group) following bleomycin or PBS instillation, or (E) mice with bleomycin or PBS instillation treated with EphB4 inhibitor NVP-BHG712 (10 mg/kg) or vehicle (n=5/group). ELN staining was semi-quantitatively scored (0–3). ****, p<0.0001 as determined by 2-way ANOVA followed by Tukey’s post-hoc tests adjusted for multiple comparisons. (F) Schematic depicting the role of EphB4 signaling in PF.

As a key regulatory cytokine for fibrogenesis, TGFβ1 typically relies on canonical signaling associated with phosphorylation of the transcription factors SMAD2/3. To assess whether SMAD2/3 activity impacts the regulatory effect of EphB4 on ELN expression, human lung fibroblasts were treated with TGFβ1 or control for 24 h, then *EPHB4* siRNA or control for an additional 24 h (**Figure 8C**). As predicted, TGFβ1 treatment increased phosphorylation of SMAD2/3 in human lung fibroblasts; however, silencing of *EPHB4* did not decrease phosphorylation levels of SMAD2/3, suggesting that EphB4 likely does not modify fibrotic pathways through canonical TGFβ1 signaling.

To assess the link between EphB4 and ELN *in vivo*, ELN staining, assessed using Verhoeff’s Van Gieson ELN stain, was examined in the lungs of bleomycin-challenged *Ephb4* CKO and *Ephb4* control mice (**Figure 8D**) and NVP-BHG712-treated mice (**Figure 8E**). Both *Ephb4* CKO and NVP-BHG712 mouse lung tissue showed qualitative decreases in ELN protein after bleomycin challenge with no changes observed in PBS controls. Thus, our data suggest that EphB4 expression is crucial for ELN expression in the context of fibrosis in both mouse tissue and human lung fibroblasts. Overall, our data supports a mechanism involving EphB4-ELN signaling in lung fibrosis.

## DISCUSSION

Ephrins and Ephs are key regulators of embryogenesis, prenatal development, and wound healing (28). While Ephrin-B2 is known to participate in the dysregulation of tissue repair during fibrosis (7, 9, 29), the role of EphB4 signaling in fibrosis remains unclear. To comprehensively explore the role of EphB4 receptor in PF development, we employed a multi-faceted approach that included the generation of *Ephb4* CKO mice in conjunction with a preclinical animal model of lung fibrosis, total and snRNA sequencing, NanoString, previously published single cell data, advanced computational biology analysis, and functional assays using mouse and human (HLF and IPF) lung fibroblasts.

To our knowledge, we are the first to demonstrate that pharmacologic inhibition or genetic deletion of EphB4 reduces the expression of fibrotic ECM-related transcriptome and attenuates the development of PF. We showed that *Col1a2-*driven deletion of *Ephb4* CKO or EphB4 pharmacological inhibition using NVP-BHG712 attenuated bleomycin-induced PF *in vivo*. Our results indicated that fibrotic injury caused transcriptomic changes in lung fibroblasts, along with the transcriptomes of specific fibroblast subpopulations. Consistent with the reduced pathological histomorphometry, we also found that genetic or pharmacological targeting of EphB4 ameliorated fibrotic injury-induced changes through reduction of select ECM genes in fibroblasts. A subset of genes identified in our mouse studies were also elevated in human lung fibroblasts from IPF patients when compared to HLFs, with ELN identified as a prominent interaction partner of EphB4 signaling in mouse and human lung fibroblasts, and across genetic and pharmacological EphB4 targeting.

To date, the role of EphB4 in injury and fibrosis remain elusive, particularly in lung tissue. EphB4 mutations causing increased expression in lung tumors has been linked to critical cell processes including tumor cell survival and proliferation (30, 31). Interestingly, we found that reduction of EphB4 in lung tissue markedly protects against development of PF, and that this antifibrotic phenotype garners profound changes in the transcriptome (317 DEGs) of mouse pulmonary fibroblasts as determined though total RNA sequencing. A subset of the 214 downregulated DEGs were linked to ECM organization, protein transport, or ER cargo regulation in fibroblasts. While the exact mechanisms of EphB4 signaling remains to be fully elucidated, downstream targets of EphB4 mediate cytoskeletal rearrangements via actin GTPase activity, which is typically associated with cell motility (29, 32). EphB4 deficiency has also been linked to accumulation of type IV collagen in endothelial cell ER through impaired protein trafficking, illuminating the potential of EphB4 as a regulator of protein transport and secretion(33). However, our in vivo data also suggests that the role of EphB4 may be context specific, as we did not observe protection from bleomycin-induced skin fibrosis in *Ephb4* CKO mice. Overall, the cytoskeletal role of EphB4 signaling may intersect with ECM secretion and protein trafficking genes observed in our study, however further investigation is required to parse the biological and pathological signaling mechanisms for EphB4 in various tissues.

Recent studies have identified distinct fibroblast cell subsets and transcriptomic profiles that may play a key role in PF development (23–25, 34–36). Using snRNA sequencing, we identified four fibroblast subclusters within mouse lung tissues. Mapping of transcriptomic profiles associated with peribronchiolar, fibrotic, alveolar and adventitial fibroblasts, identified by previous studies (23, 24), showed enrichment of these transcriptomic profiles in distinct fibroblast subsets. Fibroblast subcluster 2 from our dataset was enriched with genes associated with the fibrotic fibroblasts. It is interesting to note that subcluster 2 showed increased cellular proportionality among all fibroblasts with bleomycin treatment and was attenuated with EphB4 inhibition. Furthermore, *ELN* was identified in all fibroblast subsets with higher expression in subcluster 2 and 1, and its expression was reduced in response to NVP-BHG712 treatment under bleomycin condition.

Among the ECM genes identified in mouse lung tissue, a subset was also elevated in lung fibroblasts from IPF patients compared to donor controls; however, when IPF fibroblasts were treated with EphB4 inhibitor, *ELN* was the only ECM gene that was significantly downregulated. *ELN* also emerged as a gene that was consistently responsive to EphB4 deletion and inhibition across total RNA sequencing of IPF and control donor fibroblasts, total and snRNA sequencing of mouse lung fibroblasts, and NanoString in mouse lung fibroblasts. Our PPI results highlighted ELN as a key signaling interaction partner of EphB4 during PF. PF pathological impairment of tissue function is largely driven by excessive ECM accumulation, to which elastin reorganization and protein deposition are known indicators of disease in PF patients(19). PPI and GO analyses suggested that EphB4 and elastin signaling both interact with TGFβ1. TGFβ1 is well established to increase elastin expression in lung fibroblasts (37); however, we showed that this induction is ameliorated by *EPHB4* silencing. Furthermore, both genetic and pharmacological blockade of EphB4 was sufficient to reduce elastin in mouse lung tissue following bleomycin-induced PF, further demonstrating a crucial link between ephB4 and elastin signaling during PF

In this study, we used the bleomycin-induced model of lung fibrosis to assess the role of EphB4 in the development of pulmonary fibrosis. However, the results from this study do not indicate that EphB4 is targetable to reverse or ameliorate existing fibrosis as a therapeutic, which would require future investigation. Further, while the bleomycin preclinical model of lung fibrosis is important in understanding mechanisms of fibrosis, it does not fully capture all clinical manifestations of human IPF. For example, strong acute inflammatory phase observed in the bleomycin model is typically not present in human IPF patients (1, 2); thus, future studies should be directed towards understanding the role of Ephb4 signaling in other relevant animal models to further understand its role in PF.

Together, our *Ephb4* CKO, small molecule EphB4 inhibitor NVP-BHG712, and *EPHB4* siRNA data coupled with transcriptomic data demonstrate an important role of EphB4 signaling in PF (**Figure 8F**). Targeting EphB4 signaling requires extensive additional investigation due to the vital functions of this molecule in developmental and physiological processes, including angiogenesis (38–40), and blood pressure and heart rate regulation (41). Additional studies are also warranted to elucidate the fibrotic signaling pathways from EphB4 to the expression of its effector molecules (e.g. elastin). Signaling molecules in these pathways may provide alternative therapeutic targets for the treatment of PF and other pathological fibrotic conditions in which EphB4 signaling is implicated.

## METHODS

### Sex as a biological variable

For our animal studies, we used male and female animals for the conditional knockout mice subjected to the bleomycin model of pulmonary fibrosis. For our human cell studies involving RNA sequencing and functional assays, we obtained lung fibroblasts from both male and female subjects.

### Generation of *Ephb4* CKO mouse

*Ephb4^tm1.1Jwu^* mice, kindly donated by Dr. Jianping Wu (University of Montreal, Montreal, Canada), were crossed with B6.Cg-Tg(*Col1a2*-cre/ERT,-ALPP)7Cpd mice (The Jackson Laboratory; strain #029567) to generate *Ephb4^loxP/loxP^;Col1a2-Cre^ERT^* mice on a C57Bl/6J background. Mouse lines were backcrossed for 5 generations prior to the initiation of mouse experiments. To induce conditional *Ephb4* knockdown, 4-hydroxytamoxifen was administered by intraperitoneal (IP) injection (1 mg in 100 μL 20% ethanol/80% corn oil vehicle) for 5 consecutive days. Control mice were administered vehicle comprising 20% ethanol and 80% corn oil. Lung fibroblasts were isolated and cultured from *Ephb4* CKO mice and *Ephb4* control mice, and qRT-PCR and Western blotting was performed to assess successful deletion of EphB4.

### Mouse genotyping

Ear punches were collected, and alkaline lysis was performed at 90°C for 1 h to extract DNA. DNA was subjected to PCR against the *Ephb4* and *Cre* genes (100 V for 35 min) and gels were imaged using Sybr Safe (Thermofisher) in a BIO-RAD Chemidoc apparatus. Primers used can be found in **(Supplementary Table 12)**.

### Lung fibrosis model

To induce PF, bleomycin sulphate (1.2 U/kg in 50 μL PBS; Bioshop) was intratracheally instilled in 6–7-week-old mice that were anesthetized with isoflurane (1.5%) and xylazine/ketamine cocktail (10 mg/kg xylazine:100 mg/kg ketamine). Bleomycin control mice were administered PBS only. Instillation was initiated 7 min following xylazine/ketamine administration. Mice were sacrificed by anesthetization and CO_2_ asphyxiation 14 days following bleomycin challenge. For experiments assessing the effects of NVP-BHG712 on the development of bleomycin-induced PF, mice were intratracheally administered bleomycin sulphate (1.2 U/kg) at 7 weeks of age. At 5 days post-bleomycin challenge, the treatment with NVP-BHG712 (Sigma Aldritch; SML0333; 10 mg/kg in 150 μL N-methyl-2-pyrrolidone and polyethylene glycol 300 at ratio of 1:9) or vehicle control via oral gavage was initiated. Mice were given NVP-BHG712 once daily for a total of 5 days. Mice were sacrificed and lungs collected 2 weeks after bleomycin challenge.

### Skin Fibrosis Model

To induce skin fibrosis, mice were treated with bleomycin sulphate by subcutaneous injection. Briefly, bleomycin sulphate was reconstituted to 1 U/mL in PBS, aliquoted, and stored at -80°C. *Ephb4* CKO and *Ephb4* control mice at 8 weeks of age were anesthetized with xylazine and ketamine (10 mg/kg xylazine:100 mg/kg ketamine), and back hair was removed using Nair. Bleomycin stock solution was diluted to a working concentration (0.1 U/mL in 100 μL volume) and mice were subcutaneously treated with bleomycin daily for 21 days. Mice were sacrificed 1 day following treatment end point and four skin sections were collected for each animal.

### Lung histology

After mice were sacrificed, the heart was carefully removed and the lungs were inflated with 1 mL 10% formalin via the trachea, which was then tied off. The entire lung was excised, rinsed in PBS under gentle shaking for 1 min, and fixed in 10% formalin at room temperature with shaking for 48 h. After rinsing the lungs in PBS for 30 min, lobes were individually excised, dehydrated in reagent alcohol for 1 min with vigorous agitation, and incubated for 24 h in a minimum of 50 mL JFC solution (Milestone) per mouse lung for completion of tissue dehydration and tissue clearing. Processed lobes were paraffin-embedded with the ventral side facing the surface of the paraffin block. For staining, 4 μm lung sections were collected after removal of 80-150 μm of tissue. Gomori’s One Step Trichrome stain (Thermo) was performed according to manufacturer’s instructions to assess lung collagen content. To assess the degree of PF in affected lung tissue, Ashcroft scoring was performed across total lung sections with a minimum of 50 non-overlapping fields of view(12). For quantification of lung collagen staining across total lung tissue, lung sections were imaged at 4X magnification and stitched together using Autostitch(42) to form full lobe images. Lobes were colour corrected in GIMP image editing software V.2.10.34. Quantification of lung collagen staining was performed in Fiji V. 2.9.0 (43). Colour deconvolution was performed selecting the preset Mason’s Trichrome filter to separate blue, green and red colour channels(44). The blue colour channel was selected and thresholding was performed using the filter “mean”. Normalization to total lung tissue was performed by converting total lobe images to RGB stacks and quantifying the total pixel count using the threshold filter “default”. Lung collagen was expressed as percent lung collagen (%) by normalizing the blue colour channel to the total lobe. Lung sections from C57BL/6J mice treated with bleomycin, then NVP-BHG712 or vehicle were normalized to total tissue and collagen content was expressed as a percentage. Verhoeff Van Gieson’s stain (Abcam) was performed according to manufacturer’s instructions to assess elastin in lung tissue. Lung sections were assigned elastin expression scores on a scale of 0–3 (absent, mild, moderate, or severe staining) based on the level of staining of elastin fibers against pink and yellow counterstain.

### Immunohistochemistry

Lung sections (4 μm) were deparaffinized in xylene followed by rehydration using a graded series of alcohol washes. Antigen retrieval was performed using proteinase K (20 ug/mL) for 15 min, and endogenous peroxide was blocked for 30 min using 3% H_2_O_2_. Nonspecific IgG binding was blocked by incubating sections with 2% goat serum in PBS for 30 min. Sections were then incubated with primary antibodies for αSMA (Sigma-Aldrich; A2547) at 1:1000 overnight at 4°C. Sections were incubated with biotinylated secondary anti-mouse antibody (Vector Laboratories; BA-9200) for 30 min, followed by incubation using the Vectastain Elite ABC kit (Vector Laboratories), as per manufacturer’s direction, and counterstained with Harris hematoxylin. Lung sections were assigned αSMA positivity expression scores on a scale of 0–3 (absent, mild, moderate, or severe staining) based on the level of brown staining of αSMA against the blue counterstain.

### Skin histology

Mouse epidermis and dermis sections were collected using surgical scissors and forceps. Mouse skin was fixed in 10% formalin at room temperature for 48 h, then incubated for 24 h in a minimum of 15 mL JFC solution per 4 mouse skin sections. Skin sections were embedded in paraffin and 4 μm sections were collected. Mouse skin sections were stained with Gomori’s Onse Step Trichrome. Skin fibrosis scoring and dermal thickness analyses were performed as previously described (7).

### Lung fibroblast isolation

Mouse lungs were perfused with 10 mL PBS at 37°C through the right ventricle of the heart and out the abdominal aorta until completely blanched. Lungs were excised and rinsed in PBS. After blotting away excess PBS, lung tissue was minced by cutting with scissors 50 times. 1 mL collagenase (1 mg/mL in DMEM; Sigma) was then added, and tissue pieces were cut an additional 50 times until they were no larger than 1 mm in diameter. The lung tissue + collagenase slurry was collected and volume adjusted with collagenase to 5 mL. Tissue pieces were digested by incubating at room temperature for 15 min. Tissue digest was deactivated with 1 mL FBS and the cells were immediately passed through a 70-μm cell filter and rinsed with 10 mL DMEM. Cells were centrifuged at 300 g for 8 min and resuspended in 1 mL complete DMEM (10% FBS, 1% penicillin/streptomycin). Cells were seeded in a 6 cm plate in 4 mL complete DMEM for 24 h, at which point 50% of the media was changed. Cells were lifted with trypsin (0.05%) and passaged when they reached 80% confluence (3 ̶ 7 days). For the preparation of RNA for NanoString gene analysis, mouse lung fibroblasts from C57Bl/6J mice were treated with TGFβ1 (20 ng/mL) or PBS (control), and NVP-BHG712 (250 nM) or DMSO (vehicle control), for 24 h.

### Skin fibroblast isolation

*Ephb4* control (n=6) and *Ephb4* CKO (n=6) mice were used to isolate dermal fibroblast cells. After sacrifice, mouse skin was sterilized with 70% ethanol and fur was removed with Nair. The skin was excised, minced, and digested in collagenase for 2 h at 37°C. Cell suspension was collected and centrifuged at 300 g for 8 min. Cell pellets were resuspended in DMEM (10% FBS, 1% penicillin/streptomycin) and cultured in 60 mm culture dishes for 24 h. Mouse fibroblasts were passaged to P1 and cell supernatants were collected for RNA and protein collection.

### Primary human lung fibroblast culture and treatment

Primary human lung fibroblasts from either IPF patients (n=4; Lonza, CC-7231) or donors with no history of lung fibrosis (n=5; Lonza, CC-2512) were purchased at P2 (**Supplementary Table 7**). Fibroblasts were seeded, expanded and passaged to P4. The cells were starved overnight and subsequently treated with TGFβ1 or PBS (control) for 48 h. After 24 h of TGFβ1 treatment, NVP-BHG712 (250 nM) or DMSO vehicle was spiked into the fibroblast cultures for an additional 24 h, and RNA was isolated for qRT-PCR. For RNA sequencing, P4 primary IPF lung fibroblasts were treated with NVP-BHG712 or DMSO vehicle for 24 h. IPF lung fibroblasts were alternatively treated with silencer select siRNA (Thermo) targeted to EphB4 using RNAiMax lipofectamine according to manufacturer’s instruction. siRNA treatment was performed for 24 h at the recommended concentration of 25 pmol per well in a 6-well plate.

### Western blot

Western blot methodology can be found in the **Supplementary Methods**.

### Immunofluorescence

Human IPF fibroblasts (n=4) were cultured in 6-well plates with 150,000 cells placed on coverslips (22mm x 22mm) inside each well. When cells reached 80% confluency, they were co-treated with NVP-BHG712 (250 nM) and TGFβ1 (20 ng/mL), or DMSO and HCl as a control, for 24h. Cells were fixed in 4% paraformaldehyde. Fixed cells were permeabilized in 0.2% Triton-X and blocked with 0.5% BSA in PBS. Subsequently, cells were immunostained in the following order in the dark room. Cells were first incubated with primary anti-αSMA (Sigma, A2547; clone 1A4) in a 1:400 dilution, followed by secondary anti-mouse (Sigma, red channel, AF568) in a 1:300 dilution. Lastly, 3.5µL of Rhodamine Phalloidin (Sigma, A11055) counterstain in 500µL PBS and DAPI (1µg/mL) were applied. Imaging was done by confocal microscopy with 63x oil-immersion magnification using Zen Blue v3.12 (Zeiss). HALO v4.1 (Indica Labs) was utilized for the area quantification of fluorescence stains. To ensure the specificity in analyzing the expression of fibrotic markers, artificial lines were drawn to produce layers to capture single cells for the consecutive quantification. The five most representative single cell layers per treatment for each cell line, resulting in a total of 80 quantification data points, were obtained.

### Flow cytometry

Lungs were recovered from *Ephb4* control and *Ephb4* CKO mice after vascular perfusion with PBS, then minced and digested for 30 min at 37°C in 0.5 mg/ml collagenase VIII (Sigma). Single cells were filtered through a 70 µm filter and 1 x 10^6^ cells were labeled with a live/dead marker (eFlour780, Fisher) in PBS according to the manufacturer’s instructions. Cells were washed once, resuspended in 50 µl of FACS buffer and labeled with CD45-PE-Cy7 (Biolegend, 30F11), F4/80-APC (Biolegend, BM8), CD31-PE (Biolegend, 390) and goat anti-EphB4 (R&D, AF446), followed by donkey anti-goat-AF488 secondary (Invitrogen, A11055). At least 100,000 events were acquired using a BD FACS-Canto II flow cytometer. Single cells, live cells, and either macrophages (CD45^+^/F4/80^+^) or endothelial cells (CD45^-^ /CD31^+^) were gated. Geometric mean fluorescence intensity (gMFI) of EphB4 was determined for each gated population using FlowJo_v10.8.4. Cell sorting of similarly labeled cell populations was performed on a BD FACS ARIA III cell sorter, and cells were lysed in RIPA buffer for Western blot analysis. Total EphB4 was detected by incubating the membrane with an anti-EphB4 antibody (R&D, AF446) at 1/250 dilution, followed by a donkey anti-goat HRP secondary antibody (1:10,000).

### RNA isolation and qRT-PCR

RNA isolation and qRT-PCR methodology can be found in the Supplementary Methods. Primer sequences can be found in **Supplementary Table 13**.

### NanoString

Mouse lung fibroblasts treated with TGFβ1 and NVP-BHG712, or human lung fibroblasts from IPF donors treated with NVP-BHG712 were lysed in Trizol reagent and frozen at -80°C. Total RNA was isolated using RNeasy MinElute columns (74204; Qiagen), per the manufacturer’s instruction. RNA was quantified by Nanodrop and 100 ng of total RNA was hybridized overnight at 65°C on the nCounter instrument (NanoString Technologies, Seattle, WA), using custom designed panels for mouse or human transcripts of interest. Primers used for the NanoString gene panels can be found in **Supplementary Table 14**. Results were assessed using the nCounter analysis system on log transformed expression counts with normalization performed using the standard housekeeping controls *Acad9* and *Rplp0*. Paired 2-tailed t-test and generalized least squared regression were performed for all analyses. The resulting p-values were further adjusted for multiple comparisons according to the Benjamini-Hochberg method with q<0.05 being determined as significant.

### Total RNA sequencing and analyses

Quality of total RNA extracted from either cultured lung fibroblasts of *Ephb4* CKO mice treated with bleomycin (n=4) or PBS (n=5), *Ephb4* control mice treated with bleomycin (n=4) or PBS (n=4), or human IPF lung fibroblasts treated with NVP-BHG712 or vehicle (n=4 per treatment group) were assessed using an RNA Nano chip on an Agilent Bioanalyzer (Agilent, Santa Clara, CA, USA). RNA integrity numbers for all samples were >9.5. Samples were fluorometrically quantified using the Qubit RNA BR assay (ThermoFisher) on the Denovix DS-11 spectrophotometer (Denovix). For each sample, 200 ng RNA was used to prepare sequencing libraries using the TruSeq Stranded Total RNA with RiboZero following the low sample protocol as per manufacturer’s recommendations (Illumina), as previously described (45). Library quality was assessed on a high sensitivity DNA Bioanalyzer chip (Agilent). In total, 17 mouse libraries or 8 human libraries were volumetrically pooled and sequenced on an Illumina NextSeq 550 sequencer for 75 paired end read-cycles at the Centre for Arthritis Diagnostic and Therapeutic Innovation (CADTI, Schroeder Arthritis Institute, Krembil, Toronto, ON, Canada). Sequencing data are deposited in the gene expression omnibus (GEO) data repository under the accession codes GSE205612 and GSE205613 for the mouse and human lung fibroblasts, respectively.

Demultiplexing of samples was performed using bcl2fastq conversion tool (v2.19.1.403). Quality assessment of each sample revealed high quality of reads. To maintain minimum read length of 25 bp post trimming of adapters, Cutadapt (v2.5) software was executed (very few reads being contaminated ∼3%) along with trimming of Ns. Splice-aware alignment of reads using a Hierarchical Graph FM index (HGFM) method was performed using HISAT2 software (v2.1.0 with parameters –rna-strandness RF –dta) against mouse reference genome (vGRCm38) and human reference genome (vGRCh38) for mouse and human samples respectively. To populate the abundance of transcripts based on the reference genome and transcriptome, StringTie (v2.0.3 with parameters -e -B -G referencetranscripts.gtf) was run to generate table format output files. These files were processed using the script ‘prepDE.py’ from StringTie software to collect the raw gene expression levels for downstream processing.

For mouse lung fibroblast RNA sequencing analyses, raw gene expression levels comprised 55,421 genes measured across fibroblasts obtained from 17 mice. Lowly expressed genes were filtered out based on genes with less than 2 samples with a count per million less or equal to 10. This filter kept 10,481 genes for further analysis. For human IPF fibroblast RNA sequencing analyses, 60,675 genes were measured across 8 samples in the original matrix of counts. Low-expressed genes were filtered out based on genes with less than 2 samples with a count per million less or equal to 10. This resulted in retaining 9,619 genes for further analysis.

Changes in gene expression were evaluated using the method proposed by Anders and Huber (46), which assumes a negative binomial distribution. Function DESeq in R package DESeq2 was used with default parameters to perform differential expression analyses(47). Pairwise comparisons among groups of interest were calculated and tested for statistical differences in expression levels. All analyses include batch number as a categorical covariate in the model to control for potential batch effects. P-values were adjusted for multiple comparisons via FDR correction proposed by Benjamini and Hochberg (48). An FDR of 5% (q<0.05) was statistically significant. All analyses were performed in R version 3.5.0 (49).

### Computational analyses

Identified DEGs were analyzed using pathDIP (14) (https://ophid.utoronto.ca/pathDIP) v.5 selecting mouse as the organism, Reactome as a source, and Ortholog Pathways members as annotation, retaining only pathways with q-value<0.01(48). Up- and down-regulated genes were subjected to pathway analysis and unique pathways were further considered. GO pathway enrichment analysis was performed using Enrichr web portal (June 1, 2022) and BioPlanet 2019 pathway source (50). For network annotation, each gene was annotated with the pathway with the lowest q-value it belonged to, and a network with the gene-pathway interactions was built using NAViGaTOR v.3.0.16(15). After validation through qRT-PCR of genes involved in pathways linked to ECM, PPIs were retrieved using mouse Integrated Interaction Database (IID; https://ophid.utoronto.ca/iid) v.2021-05 (27), and shortest paths between *Ephb4* and the confirmed genes were calculated using igraph_1.2.6 in R 4.0.3 (49). Node color depicts Gene Ontology Biological Process. Network image was exported in SVG format and finalized in Adobe Illustrator version 26.3. Gene Ontology and Reactome GSEA were performed using clusterProfiler_4.6.2 and ReactomePA_1.42.0 in R 4.2.3. GSEA was performed using all deregulated genes combined, and also upregulated and downregulated genes individually.

EPHB4 and ELN were used to query IID v.2021-05 to retrieve their direct, human, physical protein interactions (27). We used NAViGaTOR v.3.0.16 to visualize the resulting network, and highlight connecting proteins and those confirmed in RNAseq data (red label)(15). Node color depicts Gene Ontology Cellular Component. Network image was exported in SVG format and finalized in Adobe Illustrator version 26.3.

### snRNA Sequencing and bioinformatics analysis

C57Bl/6J mice were subjected to intratracheal instillation of bleomycin or PBS control, then treated with NPV-BHG712 or vehicle control as described above. Mouse lungs were perfused with PBS, isolated, snap frozen, and stored at -80°C. Nuclei were isolated from frozen mouse lungs (n=12; n=3/group) by physical dissociation using a glass dounce homogenizer and were sorted using fluorescence-activated nuclei sorting (FANS) based on DAPI-positive fluorescence to obtain a single nuclei population. Nuclei concentration was determined using DAPI staining and hemocytometer, then adjusted to 1,000 nuclei/μL. SnRNA sequencing libraries were generated using the 10X Genomics Chromium Next GEM Single Cell 3’ Reagent Kits (v3.1 Dual Index). Sequencing was performed on the Illumina NextSeq 550 system (2x150 bp High Output Kit) at the CADTI. For data analysis of the single nuclei sequencing experiments, sequencing data were processed using Cell Ranger pipeline (v 7.2.0) by 10x Genomics (https://www.10xgenomics.com). Reads were aligned to the mouse transcriptome (mm10), followed by filtering and correction of cell barcodes and Unique Molecular Identifiers (UMIs). Reads associated with retained barcodes were quantified and used to build expression matrices. The standard procedures of filtering, normalization, dimensionality reduction and clustering were performed using R package Seurat (v5.0.3) (51). Low quality cells expressing less than 200 genes and low expressed genes present in less than 3 cells were removed. DecontX from Celda suite (celda_1.22.1) was used to remove ambient RNA contamination in individual cells. Cells with high mitochondrial DNA, high number of unique feature counts (potential doublets or multiplets) were excluded. We used DoubletFinder for the detection of doublets(52). Samples were merged for subsequent clustering and visualization. The batch correction between samples was done with the Harmony R package (v1.0) (53).

Data were log-normalized and highly variable genes were detected using variance-stabilizing transformation (vst). The normalized data were centered and scaled. Dimensionality reduction was performed using principal component analysis (PCA) and the most significant PCs were chosen for subsequent clustering. Graph-based clustering was implemented by calculating k-nearest neighbors, followed by modularity optimization to clusters cells. Non-linear dimensionality reduction and visualization was performed using UMAP (Uniform Manifold Approximation and Projection). Clusters were annotated on the basis of canonical markers and differential gene expression testing was used to determine a gene set signature for each cluster using the Wilcoxon Rank Sum test. Pathway enrichment analysis was performed on upregulated and downregulated genes using the Reactome database through pathDIP, and the enriched pathways were subsequently visualized using dotplots.

To explore potential cell-cell interactions across fibroblast subclusters in bleomycin samples, we employed CellChat v2.0 R package where each fibroblast subcluster was chosen as ‘receiver’ cell type and the other subclusters as ‘sender’ cell type (26). The “netVisual_chord_gene” function in CellChat was used to generate a chord diagram (circle plots) to visualize the complex ligand-receptor interactions between source and target cell types. The total number of interactions is defined as the sum of ligand–receptor pair interactions across all cell groups for each condition within the inferred pathway. The interaction strength for a given pathway is defined as the sum of communication probabilities across all cell-group pairs in the inferred network.

### Statistical Analysis

qRT-PCR and Western blot data are presented as interleaved scatter plots with horizontal lines representing the mean. The upper and lower bounds in the scatter plots correspond to the standard deviation of the data set. Statistical analysis identifying the effects of the following utilized 2-Way Anova followed by Tukey’s honest significant difference post-hoc test: determining the severity of fibrosis using either the Ashcroft scoring system or through ImageJ computational analysis, quantification of IHC scoring for αSMA in lung tissue, evaluating the mRNA expression of *Col1a1* and *Acta2* after treatment with PBS vs bleomycin and *Ephb4* control vs *Ephb4* CKO in mouse lung fibroblasts via qRT-PCR, evaluating the protein expression of ELN via Western blot after PBS vs TGFβ1 and Vehicle vs NVP-BHG712/*EPHB4* siRNA, and quantification of ELN stain scoring for the Van Gieson stain in lung tissue. Statistical analysis identifying the effects of the following utilized 2-tailed student’s T test: determining EphB4 expression or phosphorylation of EphB4 either through qRT-PCR or Western blot comparing *Ephb4* control to *Ephb4* CKO. In all comparison tests, a value of p<0.05 was considered statistically significant. Graphs and analysis were generated in GraphPad Prism version 10.

### Study Approval

Animal experiments were approved by the Animal Resources Centre under the animal use protocol number #3730, University Health Network.

### Data Availability

All datasets for RNA sequencing can be accessed online as part of the GEO super series GSE273468. Datasets for total RNA sequencing of mouse lung fibroblasts can be accessed online under accession number GSE205612. Datasets for snRNA sequencing of mouse lung fibroblasts can be accessed online under accession number GSE273467. Datasets for RNA sequencing of human lung fibroblasts can be accessed online under accession number GSE205613. Full access to all data points, means, p-values, and sample size numbers can be found in the **Supporting Data Values** file.

## Acknowledgments and Affiliations

This study is supported by grants to MK by National Science and Engineering Research Council (NSERC), Canada Research Chairs Program, Tony and Shari Fell Platinum Chair in Arthritis Research and University Health Network Foundation. BW is the recipient of the Ontario Graduate Scholarship, The Arthritis Society PhD Salary Award, and the CIHR Frederick Banting and Charles Best Canada Graduate Scholarship. IJ was supported in part by funding from Natural Sciences Research Council (NSERC RGPIN-2024-04314), Canada Foundation for Innovation (CFI #225404, #30865), and Ontario Research Fund (RDI #34876, RE010-020). The research of BH is supported by a foundation grant from the Canadian Institutes of Health Research (#375597) and support from the John Evans Leadership funds (#36050, #38861, and 38430) and innovation funds (‘Fibrosis Network, #36349’) from the Canada Foundation for Innovation (CFI) and the Ontario Research Fund (ORF). The funders had no role in study design, data collection and analysis, decision to publish, or preparation of the manuscript. The authors would like to kindly thank Dr. Evgeny Rosomacha for their help with the histology protocols and KP and JG for assisting in animal care and lab management for the duration of this study. Biorender was used to create some schematics.

## Author Contributions

BW was involved in the conception and design of the study, acquisition of data, analysis and interpretation of data, drafting the article, revising it critically for important intellectual content, and approved the final version of the manuscript. SSL and KDS prepared the cDNA libraries necessary for the sequencing experiments. SV performed the computational analysis required for the single nuclei sequencing. EPT aided in performing the intratracheal instillations on the mouse model. NF, AM, EPT, PK, AN, LT and SN aided in the acquisition and analysis of data from lung fibroblast isolations of mice and subsequent biochemical assays such as Western blotting and qRT-PCR. JK performed immunofluorescence and related analysis. HG aided in histology preparations in lung and skin samples. PP performed biostatistics on raw sequencing data. OEG performed computational analysis on RNA sequencing and NanoString data. CP and IJ performed computational analysis, gene prediction, and pathway enrichment analysis. SN aided in development of the Ephb4 CKO mouse strain, and SN and PD aided in the breeding and genotyping of the mice. JSR was involved in the interpretation of data, drafting the article, revising it critically for important intellectual content, and approved the final version of the manuscript. MBB, RG, JW, BH, and IJ were involved in the interpretation of the data and provided crucial revision on the manuscript, revising it critically for important intellectual content, and approved the final version. MK was involved in the conception and design of the study, interpretation of data, drafting the article, revising it critically for important intellectual content, and approved the final version of the manuscript.

