## Supplementary material for "Identification of Ephrin type-B receptor 4 as a critical mediator of tissue fibrosis": Supplerment.pdf

### SUPPLEMENTAL MATERIAL

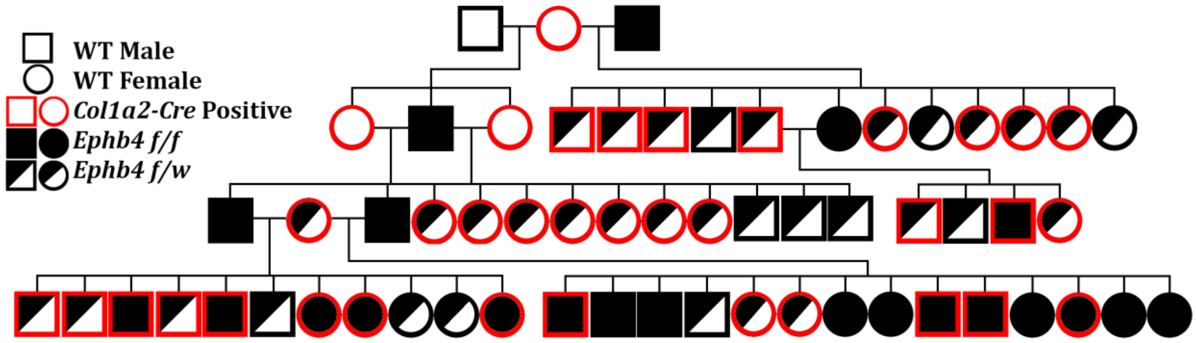


**Supplementary Figure 1. Pedigree for the generation of *Col1a2*-driven *Ephb4* CKO mouse lines.** *Ephb4*$^{tm1.1Jwu}$ mice were crossed with B6.Cg-Tg(*Col1a2*-Cre/ERT,-ALPP)7Cpd mice to generate a *Ephb4* conditional knockout (CKO) mouse line when treated with 4-hydroxytamoxifen at 4-5 weeks of age. Mice of F5 and greater were used for experiments in this study.

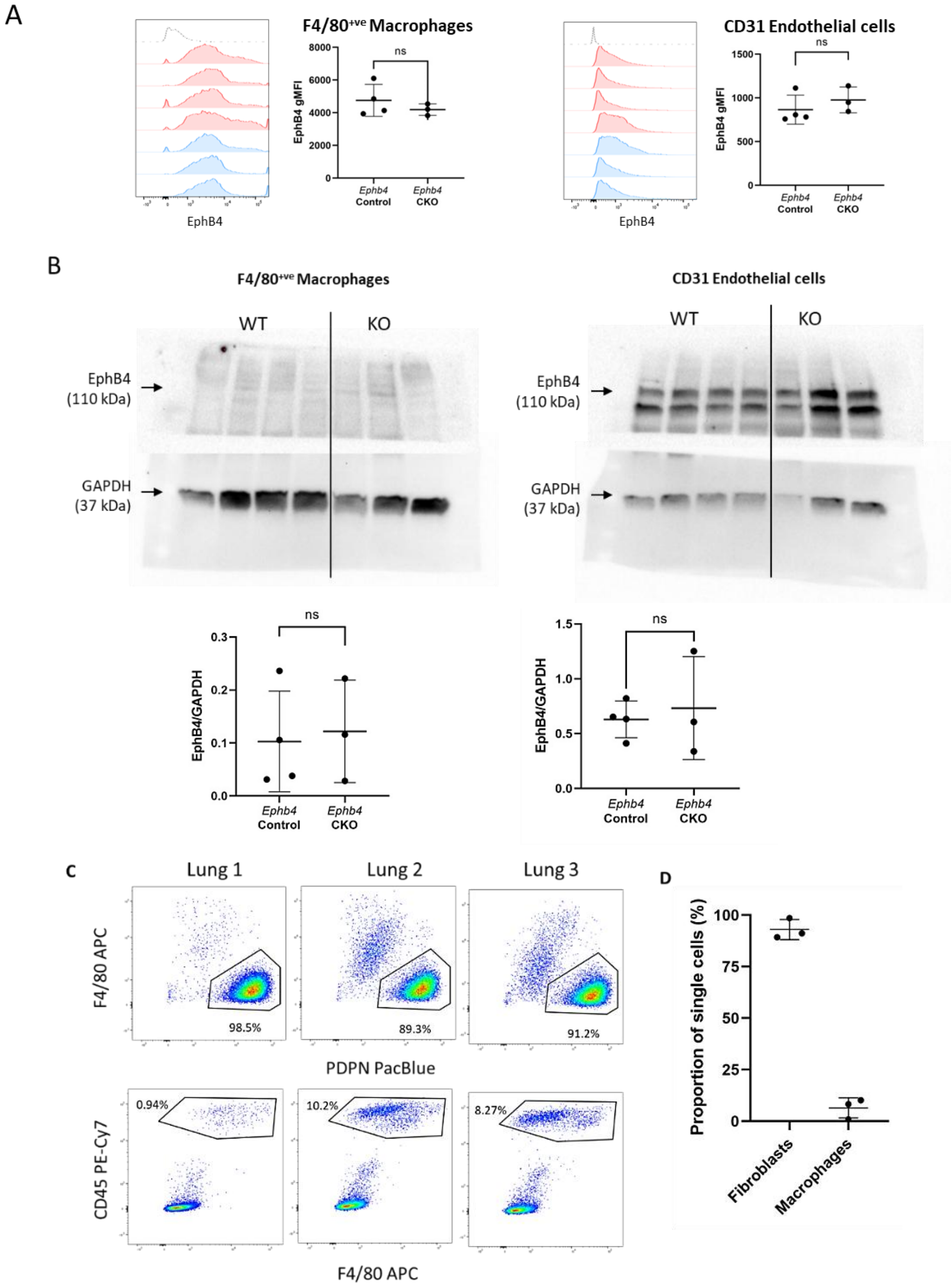


**Supplementary Figure 2: EphB4 expression is not reduced in macrophages and endothelial cells of *Ephb4* conditional knockout (CKO) mice**. Lungs were recovered from *Ephb4* control (n=4) and *Ephb4* CKO (n=3) mice and digested with collagenase to produce a single cell suspension. (A) *Ephb4* control (n=4) and *Ephb4* CKO (n=3) cells were labeled with anti-EphB4, F4/80, CD31 and a live/dead marker (eFluor780), and analyzed by flow cytometry. Relative surface expression of EphB4 was determined based on geometric mean fluorescence intensity (gMFI) after gating on macrophage and endothelial cell populations. Histograms of EphB4 expression (FITC channel) are shown including: unstained controls (dotted line), *Ephb4* control (red) and *Ephb4* CKO (blue). (B) Macrophages (F4/80+ve) and endothelial (CD31+ve) cells were sorted on a BD FACS ARIA III and lysed in RIPA buffer. The equivalent of 120,000 sorted macrophages and 240,000 sorted endothelial cells were loaded per lane. EphB4 expression was determined by densitometry and mean expression relative to GAPDH is shown. (C) Mouse lungs were digested and recovered cells were cultured in standard culture media, passaged once, and grown to confluence. Cells were processed for flow cytometry using standard procedures. (C) Scatterplots are shown indicating percentages of PDPN+ve and F4/80+ve populations for 3 independent mice. (D) Mean percentages (±StDev) of gated populations are shown for PDPN+ve fibroblasts and F40/80+ve, CD45+ve macrophages (n=3).

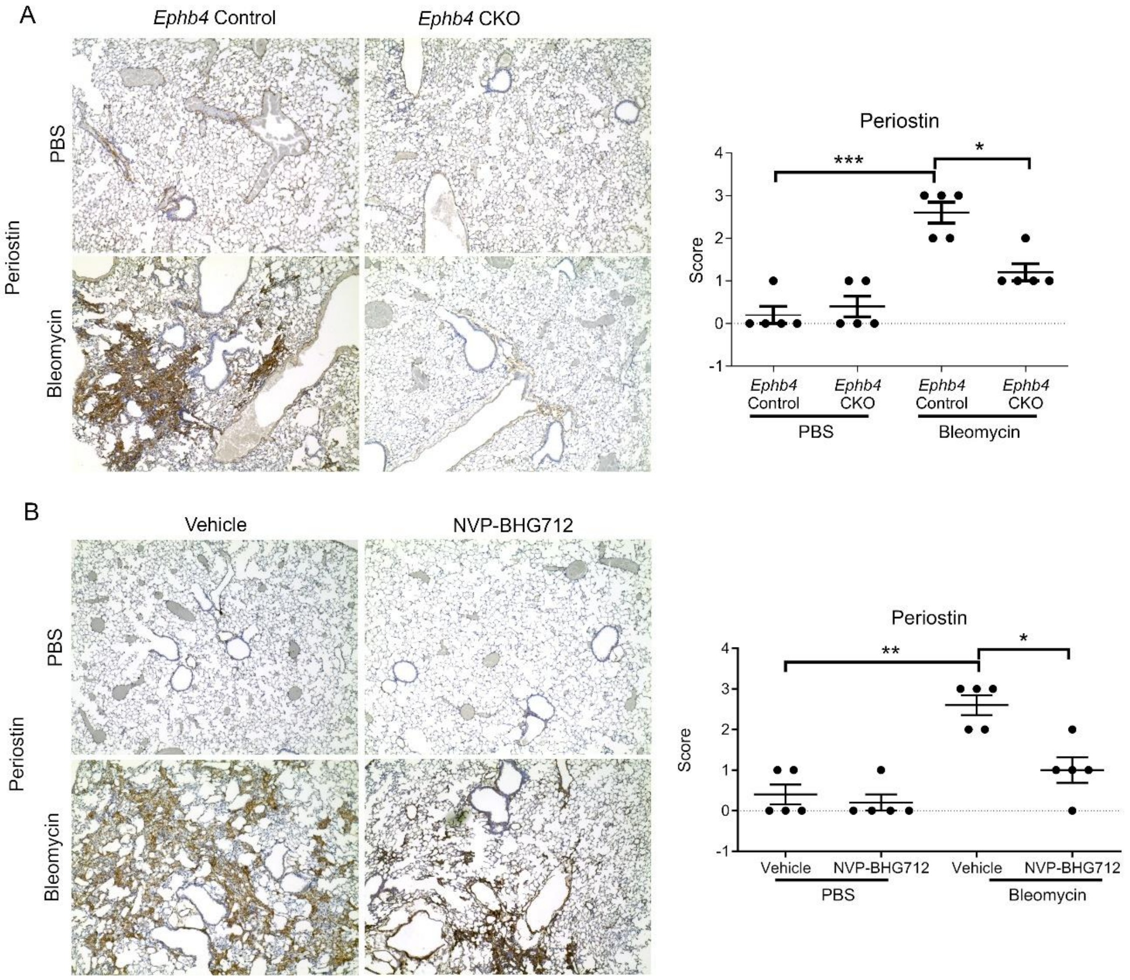


**Supplementary Figure 3: EphB4 conditional knockout or inhibition by NVP-BHG712 reduces the expression of periostin in mouse lung tissue.** PBS or bleomycin intratracheal instillation were performed on (A) *Ephb4* control and *Ephb4* CKO mice, or (B) mice treated with EphB4 inhibitor NVP-BHG712 or vehicle (n=6/group). Periostin staining was performed in lung tissue sections to assess myofibroblast populations in mouse lungs. Semi-quantitative scoring (0-3) of the staining intensity was assigned to each mouse lung. For all graphs, data are expressed as mean ± SD where significant differences (*, $p < 0.05$; **, $p < 0.01$; ***, $p < 0.001$) were identified using 2-way ANOVA with Tukey's post-hoc tests.

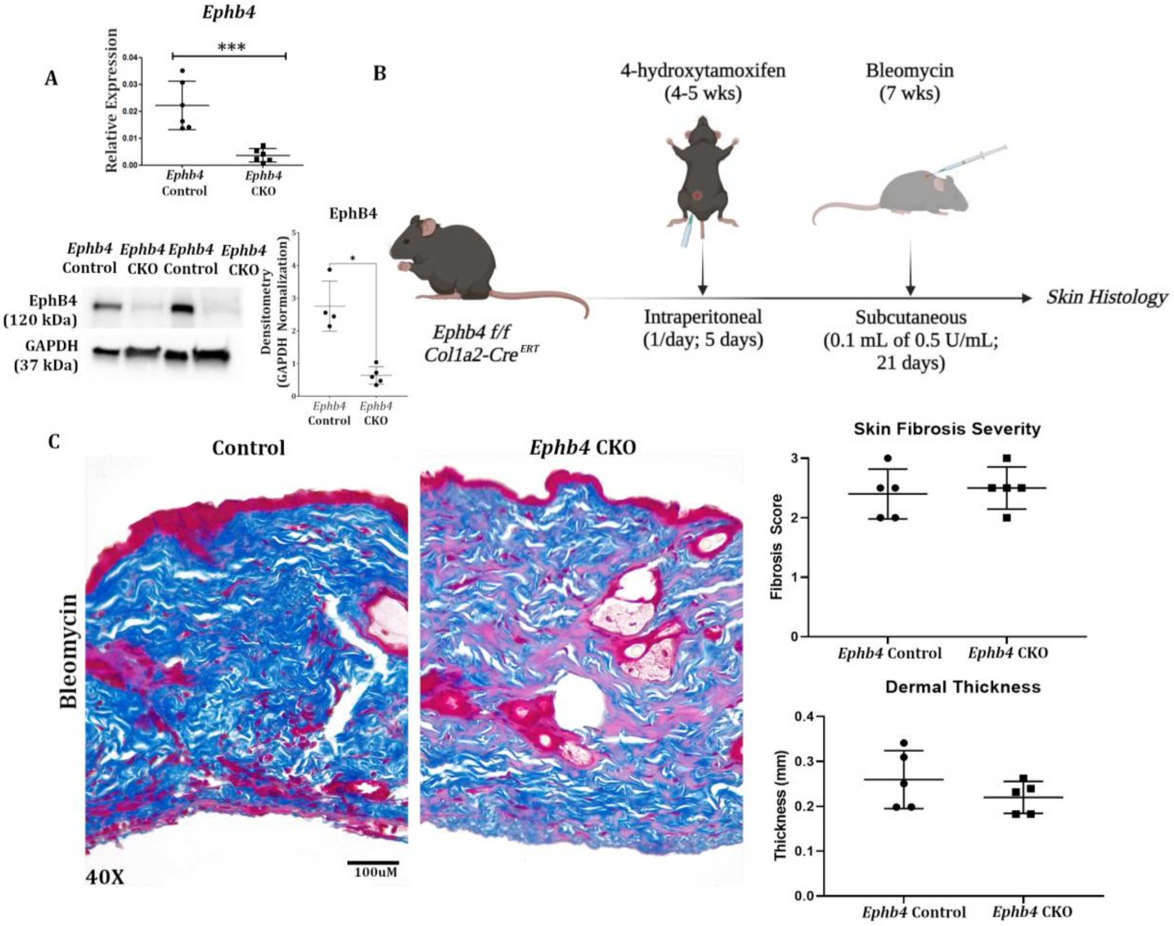


**Supplementary Figure 4. Conditional knockdown of *Ephb4* in mouse fibroblasts does not confer protection in a skin fibrosis model.** (A) Effective knockdown of *Ephb4* expression was observed at both the transcript (qRT-PCR, n=6) and protein levels (Western blotting; n=4, *Ephb4* control; n=5, *Ephb4* CKO) in P1 mouse skin fibroblasts isolated 7 days after final treatment with 4-hydroxytamoxifen (1 mg/day for 5 days; *i.p.*). Significance was determined using unpaired student's T tests. *, $p<0.05$; ***, $p<0.001$. (B) *Ephb4*$^{f/f}$*;Col1a2-Cre*$^{ERT}$ mice were treated with either 4-hydroxytamoxifen (*Ephb4* CKO) or corn oil (*Ephb4* control) at 4–5 weeks of age. To induce skin fibrosis, mice received subcutaneous injections of bleomycin (100 µL of 0.5 U/mL dissolved in PBS for 21 days). (C) Representative images of dermis sections stained with Gomori's One Step Trichrome from *Ephb4* CKO and *Ephb4* control mice (n=5, 4 sections/animal). To assess skin fibrosis severity, skin fibrosis scoring (0-3) and dermal thickness were quantified with no significant differences between groups found ($p< 0.05$).

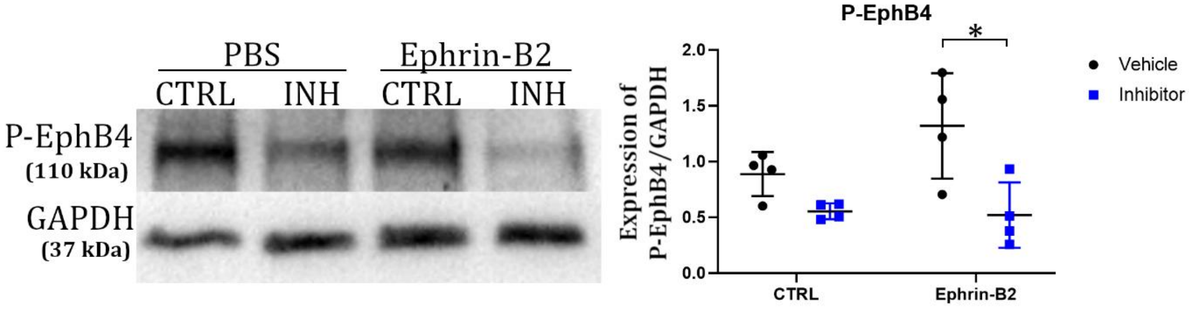


**Supplementary Figure 5. NVP-BHG712 treatment of human lung fibroblasts inhibits EphB4 phosphorylation.** Primary human lung fibroblasts were treated with pre-clustered Ephrin-B2 (1 µg/mL; or PBS control) and NVP-BHG712 (250 nM; or DMSO vehicle) for 24h (n=4). Western blotting was performed on protein lysates to detect phospho(P)-EphB4. GAPDH was used as a housekeeping control. Significance was determined by 2-way ANOVA followed by Tukey's post-hoc tests adjusted for multiple comparisons. *, $p < 0.05$.

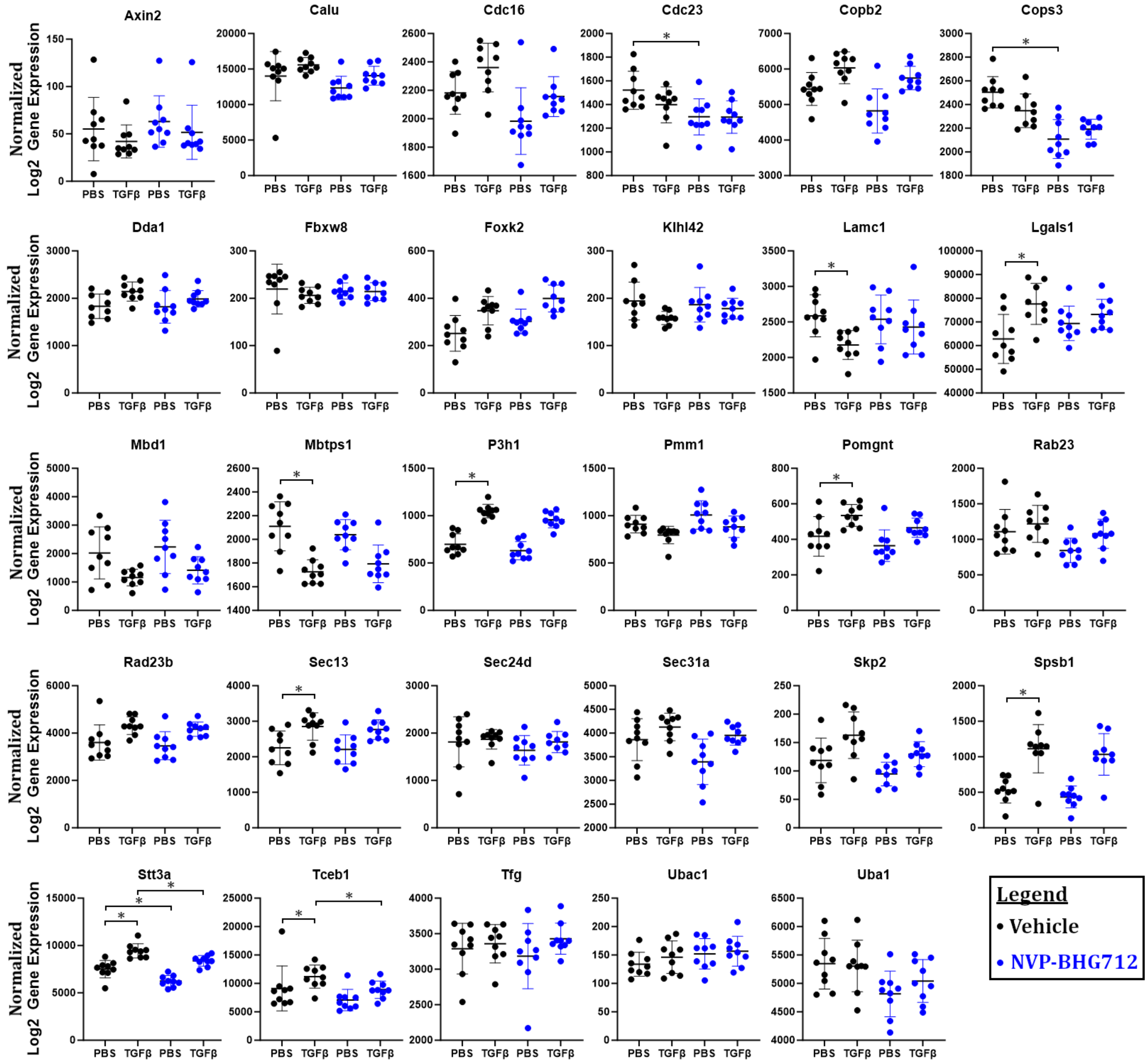


**Supplementary Figure 6. Expression of remaining genes in the NanoString codeset.** Transcript levels of the remaining genes not included in **Figure 4** in response to 24 h co-treatment of P4 mouse lung fibroblasts (n=9) with TGF-β1 (20 ng/mL; or PBS control) and NVP-BHG712 (250 nM; or DMSO vehicle). RNA was isolated and NanoString was performed to examine the transcript levels of downregulated genes identified by RNAseq that were linked to significantly enriched pathways. Wilcoxon signed rank test and FDR q≤0.05 performed to identify significant differences in gene expression between treatment groups. *, q < 0.05.

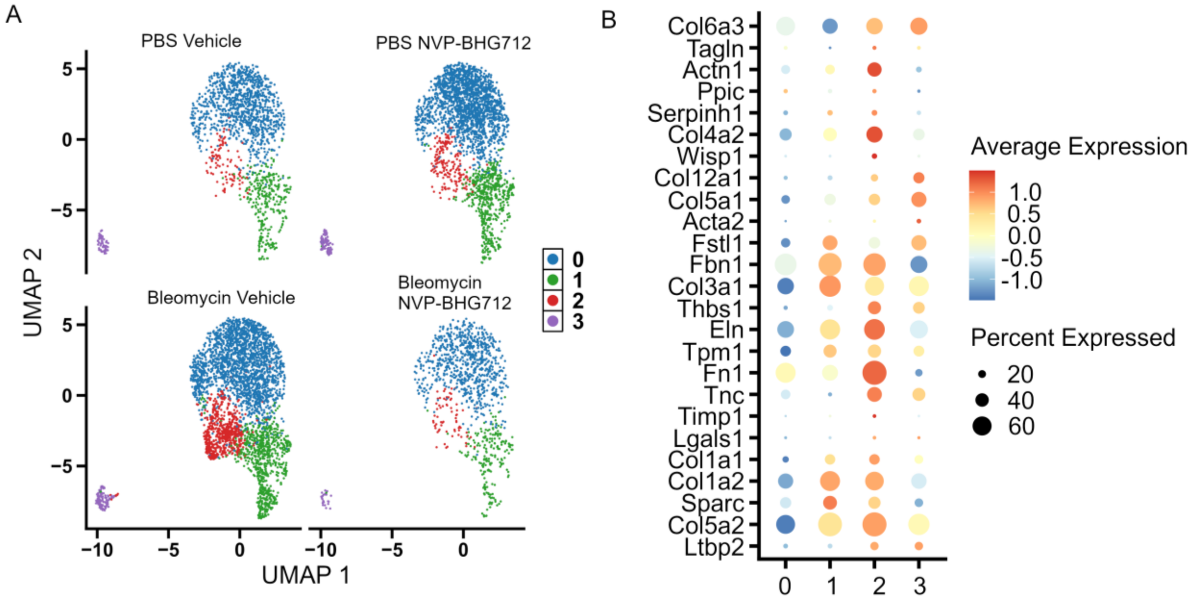


**Supplementary Figure 7. Identification of fibroblast subclusters in mouse lungs and association of fibroblast subclusters with an activated fibroblast gene profile.** (A) UMAP projection of distinct fibroblast populations identified in lung tissue from bleomycin- or PBS-treated mice that were administered NVP-BHG712 or vehicle. (B) Expression of key marker genes associated with activated fibroblasts from Peyser et al.[23]. were mapped to our snRNAseq dataset.

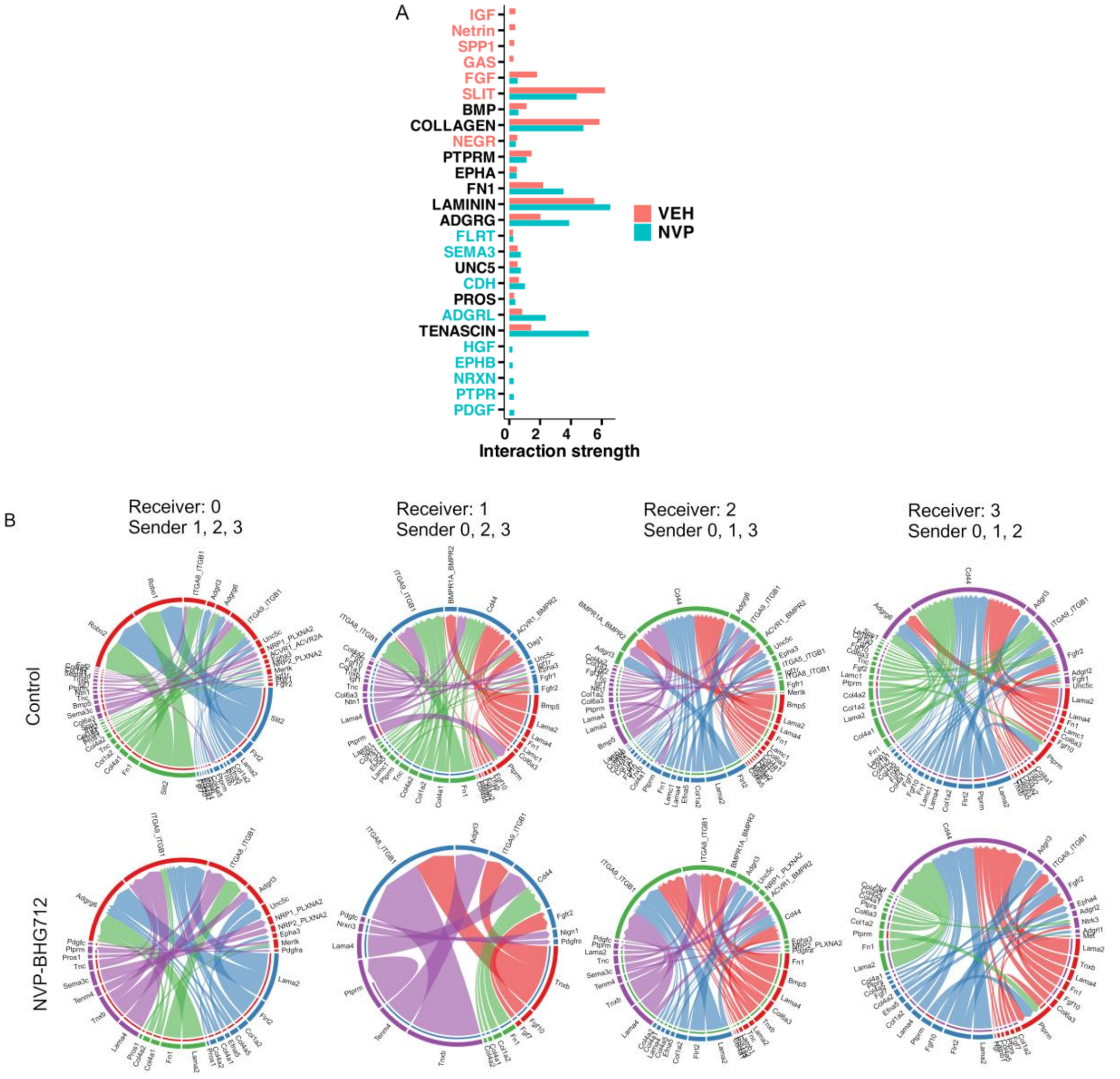


**Supplementary Figure 8. Comparison of cell-cell interactions in lung fibroblasts from bleomycin-treated treated with vehicle compared to EphB4 inhibitor.** (A) Cell-cell communication between fibroblast subclusters in bleomycin-treated vehicle and NVP-BHG712 groups showing significant interactions across various signaling pathways. (B) Comparison of the ligand-receptor pair interactions between vehicle and NVP-BHG712 that contribute to signaling between fibroblast subclusters.

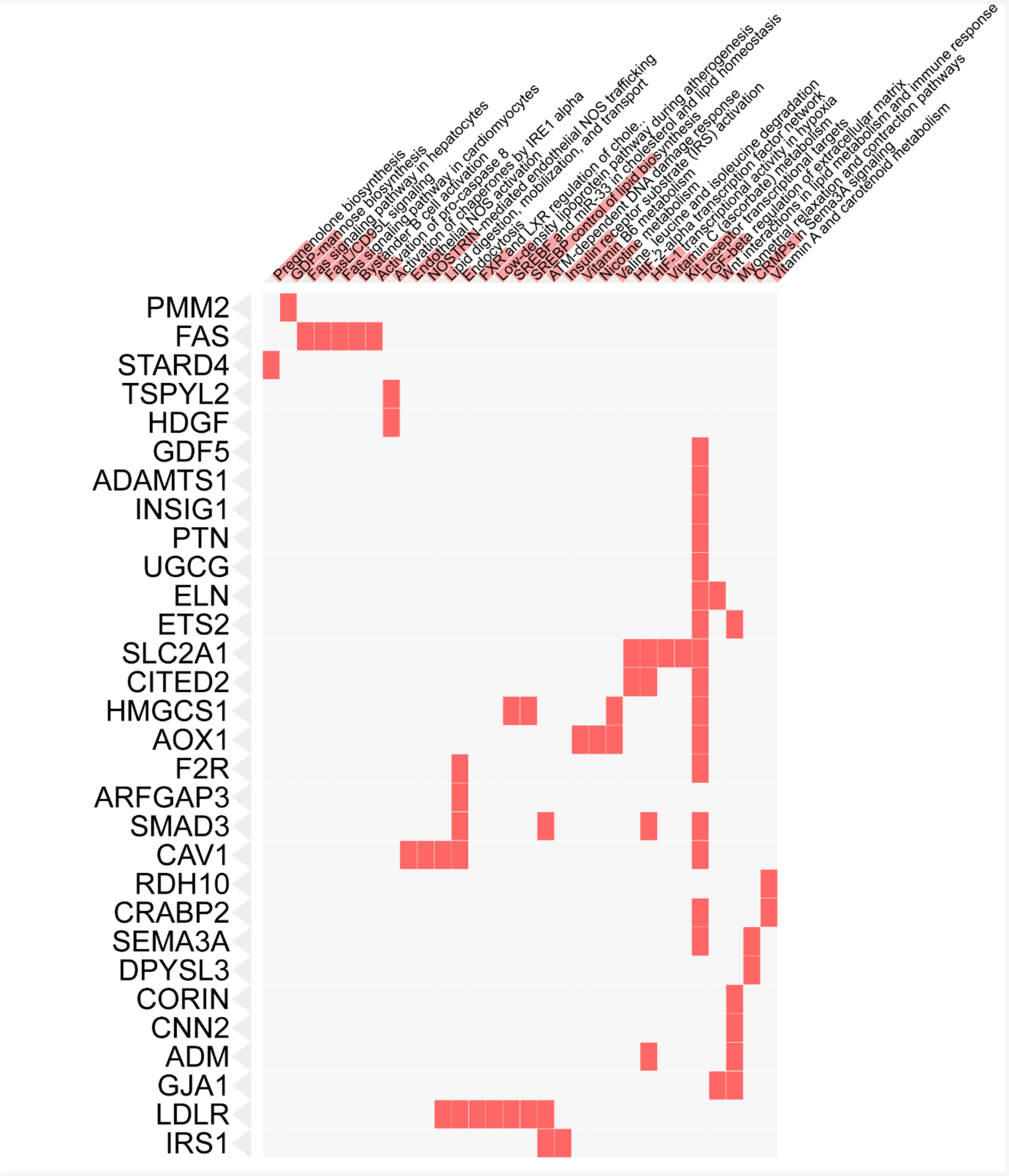


**Supplementary Figure 9. GO enrichment analysis of transcriptomes of IPF fibroblasts treated with NVP-BHG712 compared to vehicle controls**. Primary IPF patient fibroblasts were treated with NVP-BHG712 (250 nM) for 24 h and RNA was harvested for total RNA sequencing. DEGs were identified and pathway enrichment analysis was performed considering down regulated genes, using Enrichr web portal (June 1, 2022) and BioPlanet 2019 pathway sources. Highlighted association of input genes with enriched pathways is presented using combined scores.

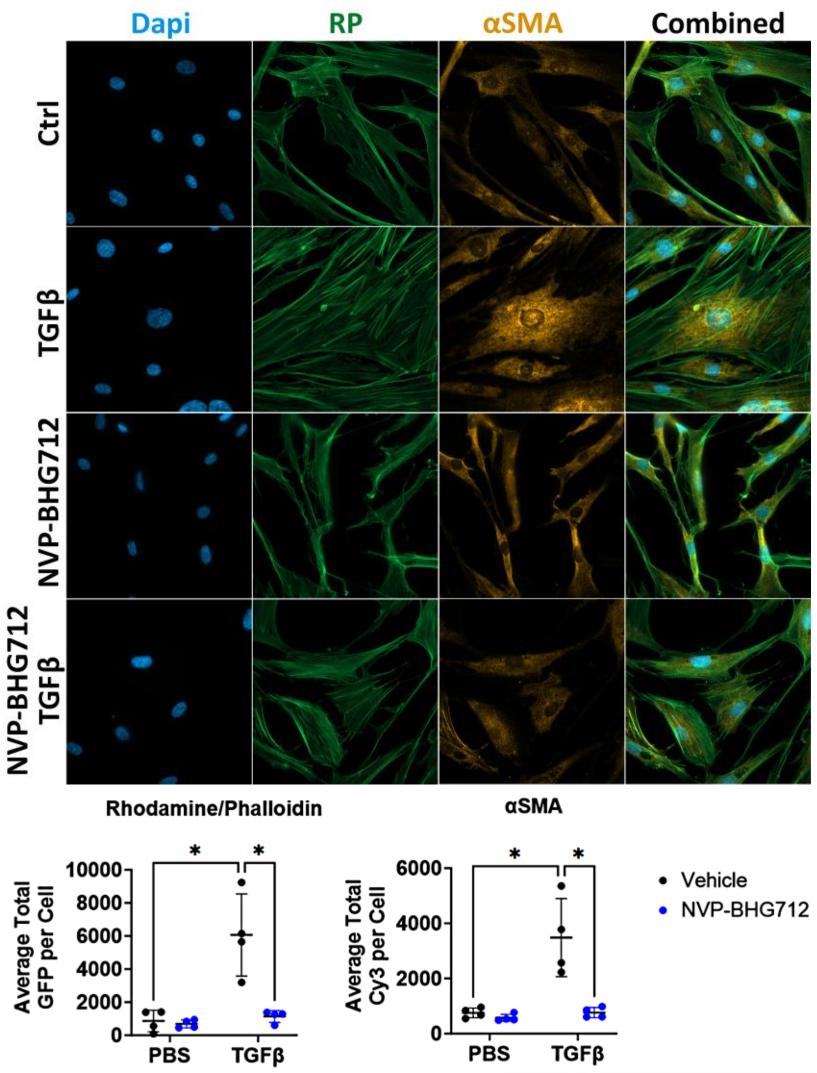


**Supplementary Figure 10. Immunofluorescence of α-SMA and rhodamine/phalloidin in human lung fibroblasts from IPF patients.** Human IPF fibroblasts (n=4) were cultured and co-treated with NVP-BHG712 (250nM) and TGFβ (20ng/mL) or respective vehicle (as a control) for 24h. Subsequently, cells were immunostained with α-SMA and Rhodamine Phalloidin to observe fibrogenic changes. Imaging was done by confocal microscopy with 63x oil-immersion magnification. Quantification in HALO was performed by averaging total intensity per single cell for 5 cells per patient per group (n=4 patients/group). Two-way ANOVA was conducted on log-transformed data to infer statistical significance followed by Tukey's post-hoc tests for multiple comparisons. *, $p < 0.05$.

**SUPPLEMENTARY TABLES**

*The following Supplementary Tables can be accessed in attached supplementary files:*

**Supplementary Table 4.** Pathway and Gene Ontology GSEA results comparing *Ephb4* CKO vs control mice treated with bleomycin**.**

**Supplementary Table 6.** Genes associated with each fibroblast subcluster identified using single nuclei RNA sequencing.

**Supplementary Table 10.** PPIs between EphB4 and Elastin in the Integrated Interaction Database ver. 2021-05 (14)(https://ophid.utoronto.ca/iid).

**Supplementary Table 1. Fibrotic lobe counts in *Ephb4* CKO mice compared to control mice.**

| PBS-Challenged Mice | | | | Bleomycin-Challenged Mice | | | |
|---|---|---|---|---|---|---|---|
| ***Ephb4* Control Animal** | **# of Affected Lobes** | ***Ephb4* CKO Animals** | **# of Affected Lobes** | ***Ephb4* Control Animal** | **# of Affected Lobes** | ***Ephb4* CKO Animals** | **# of Affected Lobes** |
| *Ephb4* Control 1 | 0/5 | *Ephb4* CKO 1 | 0/5 | *Ephb4* Control 1 | 4/5 | *Ephb4* CKO 1 | 1/5 |
| *Ephb4* Control 2 | 0/5 | *Ephb4* CKO 2 | 0/5 | *Ephb4* Control 2 | 2/5 | *Ephb4* CKO 2 | 0/5 |
| *Ephb4* Control 3 | 0/5 | *Ephb4* CKO 3 | 0/5 | *Ephb4* Control 3 | 4/5 | *Ephb4* CKO 3 | 2/5 |
| *Ephb4* Control 4 | 0/5 | *Ephb4* CKO 4 | 0/5 | *Ephb4* Control 4 | 3/5 | *Ephb4* CKO 4 | 0/5 |
| *Ephb4* Control 5 | 0/5 | *Ephb4* CKO 5 | 0/5 | *Ephb4* Control 5 | 3/5 | *Ephb4* CKO 5 | 2/5 |
| *Ephb4* Control 6 | 0/5 | *Ephb4* CKO 6 | 0/5 | *Ephb4* Control 6 | 5/5 | *Ephb4* CKO 6 | 2/5 |
| *Ephb4* Control 7 | 0/5 | *Ephb4* CKO 7 | 0/5 | *Ephb4* Control 7 | 3/5 | *Ephb4* CKO 7 | 5/5 |
| *Ephb4* Control 8 | 0/5 | *Ephb4* CKO 8 | 0/5 | *Ephb4* Control 8 | 2/5 | *Ephb4* CKO 8 | 0/5 |

**Supplementary Table 2A. Top 50 upregulated DEGs comparing Ephb4 CKO to control mouse fibroblasts following bleomycin treatment.**

| geneSymbol | baseMean | log2FoldChange | lfcSE | stat | pvalue | padj |
|---|---|---|---|---|---|---|
| Rin2 | 1.09E+04 | 1.27E+00 | 2.39E-01 | 5.32E+00 | 1.01E-07 | 2.73E-04 |
| Ralgapa2 | 3.57E+03 | 9.67E-01 | 1.88E-01 | 5.16E+00 | 2.52E-07 | 4.29E-04 |
| Pank2 | 3.38E+03 | 3.98E-01 | 8.95E-02 | 4.45E+00 | 8.63E-06 | 4.08E-03 |
| Rbm47 | 1.96E+03 | 1.40E+00 | 3.15E-01 | 4.43E+00 | 9.24E-06 | 4.11E-03 |
| Cyb5r4 | 6.63E+03 | 1.03E+00 | 2.35E-01 | 4.36E+00 | 1.29E-05 | 4.80E-03 |
| Ly6i | 1.28E+03 | 4.96E+00 | 1.15E+00 | 4.33E+00 | 1.51E-05 | 5.15E-03 |
| Cd36 | 1.26E+05 | 2.79E+00 | 6.47E-01 | 4.30E+00 | 1.67E-05 | 5.42E-03 |
| Lpl | 9.06E+04 | 2.77E+00 | 6.55E-01 | 4.23E+00 | 2.35E-05 | 6.59E-03 |
| Grk3 | 1.37E+03 | 1.26E+00 | 2.98E-01 | 4.22E+00 | 2.40E-05 | 6.63E-03 |
| Cmip | 2.44E+03 | 6.97E-01 | 1.69E-01 | 4.12E+00 | 3.73E-05 | 8.26E-03 |
| Gm6545 | 5.50E+02 | 5.00E+00 | 1.22E+00 | 4.11E+00 | 4.04E-05 | 8.26E-03 |
| Clcc1 | 3.00E+03 | 5.37E-01 | 1.31E-01 | 4.09E+00 | 4.32E-05 | 8.49E-03 |
| Slc6a12 | 4.54E+03 | 2.25E+00 | 5.64E-01 | 3.98E+00 | 6.75E-05 | 1.09E-02 |
| Usp6nl | 3.49E+03 | 5.19E-01 | 1.32E-01 | 3.93E+00 | 8.52E-05 | 1.30E-02 |
| A930007I19Rik | 1.76E+03 | 2.34E+00 | 5.97E-01 | 3.92E+00 | 8.68E-05 | 1.31E-02 |
| Coro2a | 6.03E+03 | 1.62E+00 | 4.18E-01 | 3.89E+00 | 9.99E-05 | 1.42E-02 |
| Alas1 | 6.25E+04 | 2.19E+00 | 5.64E-01 | 3.89E+00 | 1.01E-04 | 1.42E-02 |
| Pik3cb | 5.52E+03 | 1.14E+00 | 2.95E-01 | 3.87E+00 | 1.07E-04 | 1.45E-02 |
| Gm13571 | 3.06E+03 | 3.82E+00 | 9.86E-01 | 3.88E+00 | 1.06E-04 | 1.45E-02 |
| Mlxip | 3.04E+03 | 6.25E-01 | 1.62E-01 | 3.87E+00 | 1.10E-04 | 1.48E-02 |
| Cbl | 5.31E+03 | 9.92E-01 | 2.57E-01 | 3.87E+00 | 1.11E-04 | 1.48E-02 |
| Lpin2 | 5.61E+03 | 9.39E-01 | 2.43E-01 | 3.86E+00 | 1.11E-04 | 1.48E-02 |
| Gm20056 | 7.94E+02 | 3.40E+00 | 8.95E-01 | 3.80E+00 | 1.47E-04 | 1.65E-02 |
| Zfand5 | 1.04E+04 | 7.89E-01 | 2.08E-01 | 3.79E+00 | 1.49E-04 | 1.66E-02 |
| Sft2d2 | 5.45E+03 | 8.43E-01 | 2.22E-01 | 3.79E+00 | 1.50E-04 | 1.66E-02 |
| Cracr2b | 1.09E+03 | 1.29E+00 | 3.39E-01 | 3.79E+00 | 1.50E-04 | 1.66E-02 |
| Tmem106c | 1.45E+03 | 9.62E-01 | 2.55E-01 | 3.77E+00 | 1.61E-04 | 1.72E-02 |
| Ly6c2 | 9.69E+02 | 3.57E+00 | 9.48E-01 | 3.77E+00 | 1.62E-04 | 1.72E-02 |
| Galm | 2.19E+03 | 1.38E+00 | 3.69E-01 | 3.75E+00 | 1.75E-04 | 1.81E-02 |
| Crybg1 | 3.75E+03 | 1.43E+00 | 3.83E-01 | 3.75E+00 | 1.78E-04 | 1.83E-02 |
| Cd200 | 1.62E+04 | 2.13E+00 | 5.69E-01 | 3.74E+00 | 1.87E-04 | 1.89E-02 |
| Prkcd | 2.48E+04 | 1.11E+00 | 3.00E-01 | 3.71E+00 | 2.06E-04 | 1.99E-02 |
| Ahcyl2 | 4.06E+03 | 9.28E-01 | 2.51E-01 | 3.69E+00 | 2.20E-04 | 2.05E-02 |
| Tns4 | 3.78E+02 | 1.56E+00 | 4.23E-01 | 3.68E+00 | 2.31E-04 | 2.09E-02 |
| Il1rn | 3.62E+04 | 2.69E+00 | 7.40E-01 | 3.64E+00 | 2.75E-04 | 2.28E-02 |
| Shtn1 | 9.94E+03 | 1.40E+00 | 3.85E-01 | 3.63E+00 | 2.83E-04 | 2.32E-02 |
| Nostrin | 1.06E+03 | 1.84E+00 | 5.08E-01 | 3.62E+00 | 3.00E-04 | 2.38E-02 |
| Met | 2.74E+04 | 2.71E+00 | 7.51E-01 | 3.61E+00 | 3.01E-04 | 2.39E-02 |
| Cdca7l | 1.84E+03 | 1.47E+00 | 4.07E-01 | 3.60E+00 | 3.24E-04 | 2.45E-02 |
| Ptk2b | 8.80E+03 | 1.14E+00 | 3.17E-01 | 3.58E+00 | 3.38E-04 | 2.48E-02 |
| Bcl2a1d | 1.02E+04 | 1.76E+00 | 4.91E-01 | 3.58E+00 | 3.44E-04 | 2.50E-02 |
| Vsig8 | 1.70E+03 | 3.28E+00 | 9.17E-01 | 3.58E+00 | 3.45E-04 | 2.50E-02 |
| Ttc5 | 2.16E+03 | 4.71E-01 | 1.32E-01 | 3.57E+00 | 3.54E-04 | 2.52E-02 |
| Nfya | 3.09E+03 | 7.34E-01 | 2.06E-01 | 3.57E+00 | 3.54E-04 | 2.52E-02 |
| Gpbp1l1 | 6.60E+03 | 4.56E-01 | 1.28E-01 | 3.55E+00 | 3.81E-04 | 2.63E-02 |
| Chd7 | 2.67E+03 | 8.45E-01 | 2.38E-01 | 3.55E+00 | 3.86E-04 | 2.65E-02 |
| Ezh1 | 3.36E+03 | 9.96E-01 | 2.81E-01 | 3.55E+00 | 3.89E-04 | 2.65E-02 |
| Parp4 | 1.30E+03 | 7.59E-01 | 2.14E-01 | 3.54E+00 | 3.97E-04 | 2.69E-02 |
| Parp6 | 1.47E+03 | 6.85E-01 | 1.94E-01 | 3.53E+00 | 4.13E-04 | 2.69E-02 |
| Prdx1 | 2.31E+05 | 2.13E+00 | 6.03E-01 | 3.53E+00 | 4.09E-04 | 2.69E-02 |
| Hsd17b11 | 1.39E+03 | 9.29E-01 | 2.63E-01 | 3.52E+00 | 4.24E-04 | 2.72E-02 |

**Supplementary Table 2B. Top 50 downregulated DEGs comparing *Ephb4* CKO to control mouse fibroblasts following bleomycin treatment**.

| geneSymbol | baseMean | log2FoldChange | lfcSE | stat | pvalue | padj |
|---|---|---|---|---|---|---|
| Rnu3a | 9.76E+02 | -2.65E+01 | 4.39E+00 | -6.03E+00 | 1.61E-09 | 1.10E-05 |
| Eln | 1.30E+03 | -7.53E+00 | 2.13E+00 | -3.53E+00 | 4.12E-04 | 2.69E-02 |
| Col11a1 | 2.92E+03 | -5.42E+00 | 1.70E+00 | -3.20E+00 | 1.39E-03 | 4.99E-02 |
| Tnfsf15 | 3.07E+03 | -5.29E+00 | 1.03E+00 | -5.13E+00 | 2.82E-07 | 4.44E-04 |
| Fibin | 8.44E+02 | -4.82E+00 | 1.51E+00 | -3.20E+00 | 1.39E-03 | 4.98E-02 |
| Nrep | 2.56E+03 | -4.81E+00 | 1.19E+00 | -4.04E+00 | 5.32E-05 | 9.79E-03 |
| Pappa | 2.03E+03 | -4.79E+00 | 1.14E+00 | -4.19E+00 | 2.77E-05 | 7.02E-03 |
| Axin2 | 2.75E+02 | -4.77E+00 | 1.04E+00 | -4.58E+00 | 4.71E-06 | 2.83E-03 |
| Timp3 | 3.70E+04 | -4.74E+00 | 9.40E-01 | -5.04E+00 | 4.57E-07 | 5.38E-04 |
| Cthrc1 | 1.11E+03 | -4.68E+00 | 1.44E+00 | -3.24E+00 | 1.18E-03 | 4.61E-02 |
| Gm21982 | 3.13E+02 | -4.63E+00 | 1.35E+00 | -3.43E+00 | 5.95E-04 | 3.24E-02 |
| Hs3st1 | 4.41E+02 | -4.62E+00 | 1.11E+00 | -4.15E+00 | 3.32E-05 | 7.53E-03 |
| Cdh6 | 2.79E+02 | -4.55E+00 | 1.18E+00 | -3.87E+00 | 1.07E-04 | 1.45E-02 |
| Has2 | 5.70E+03 | -4.52E+00 | 1.27E+00 | -3.56E+00 | 3.72E-04 | 2.60E-02 |
| Crispld2 | 1.93E+03 | -4.43E+00 | 1.33E+00 | -3.34E+00 | 8.40E-04 | 3.86E-02 |
| Gm21451 | 3.86E+03 | -4.42E+00 | 1.25E+00 | -3.54E+00 | 3.97E-04 | 2.69E-02 |
| Loxl2 | 1.13E+04 | -4.36E+00 | 1.24E+00 | -3.51E+00 | 4.46E-04 | 2.80E-02 |
| Dkk2 | 8.53E+02 | -4.31E+00 | 1.23E+00 | -3.50E+00 | 4.61E-04 | 2.86E-02 |
| Aldh1a2 | 3.16E+03 | -4.15E+00 | 8.24E-01 | -5.04E+00 | 4.74E-07 | 5.38E-04 |
| Tll1 | 6.67E+02 | -4.04E+00 | 1.09E+00 | -3.69E+00 | 2.21E-04 | 2.05E-02 |
| Fap | 2.34E+02 | -3.99E+00 | 1.22E+00 | -3.27E+00 | 1.06E-03 | 4.40E-02 |
| Nox4 | 6.87E+02 | -3.99E+00 | 1.18E+00 | -3.39E+00 | 7.08E-04 | 3.52E-02 |
| Aldh1l2 | 3.16E+03 | -3.67E+00 | 1.07E+00 | -3.42E+00 | 6.16E-04 | 3.29E-02 |
| Piezo2 | 1.67E+03 | -3.60E+00 | 1.08E+00 | -3.34E+00 | 8.42E-04 | 3.86E-02 |
| Prickle1 | 9.12E+02 | -3.57E+00 | 1.04E+00 | -3.42E+00 | 6.16E-04 | 3.29E-02 |
| Wnt5a | 1.17E+03 | -3.35E+00 | 1.02E+00 | -3.27E+00 | 1.08E-03 | 4.43E-02 |
| Csrp2 | 1.60E+03 | -3.33E+00 | 7.49E-01 | -4.45E+00 | 8.66E-06 | 4.08E-03 |
| Ramp2 | 2.58E+02 | -3.22E+00 | 9.18E-01 | -3.51E+00 | 4.53E-04 | 2.82E-02 |
| Lox | 2.43E+04 | -3.06E+00 | 8.13E-01 | -3.76E+00 | 1.67E-04 | 1.75E-02 |
| Tspan18 | 3.43E+02 | -3.06E+00 | 9.09E-01 | -3.36E+00 | 7.71E-04 | 3.67E-02 |
| Ccdc80 | 7.74E+03 | -2.86E+00 | 6.63E-01 | -4.31E+00 | 1.62E-05 | 5.39E-03 |
| Pxdn | 5.07E+03 | -2.83E+00 | 8.67E-01 | -3.27E+00 | 1.09E-03 | 4.45E-02 |
| Zfp9 | 9.10E+02 | -2.73E+00 | 8.49E-01 | -3.21E+00 | 1.32E-03 | 4.85E-02 |
| Creb3l1 | 1.28E+03 | -2.65E+00 | 7.55E-01 | -3.51E+00 | 4.42E-04 | 2.79E-02 |
| Spsb1 | 3.14E+03 | -2.62E+00 | 7.77E-01 | -3.37E+00 | 7.57E-04 | 3.64E-02 |
| Mex3b | 3.06E+02 | -2.61E+00 | 7.96E-01 | -3.28E+00 | 1.03E-03 | 4.31E-02 |
| Nxn | 3.07E+03 | -2.57E+00 | 8.03E-01 | -3.20E+00 | 1.39E-03 | 4.98E-02 |
| Lgalsl | 1.12E+03 | -2.55E+00 | 6.91E-01 | -3.69E+00 | 2.25E-04 | 2.06E-02 |
| Lbh | 3.07E+03 | -2.52E+00 | 6.62E-01 | -3.81E+00 | 1.37E-04 | 1.61E-02 |
| Lrig3 | 6.14E+02 | -2.42E+00 | 4.93E-01 | -4.91E+00 | 9.07E-07 | 8.06E-04 |
| Zfp449 | 5.43E+02 | -2.17E+00 | 6.18E-01 | -3.51E+00 | 4.53E-04 | 2.82E-02 |
| P3h1 | 3.21E+03 | -2.10E+00 | 5.11E-01 | -4.12E+00 | 3.85E-05 | 8.26E-03 |
| Crim1 | 3.38E+03 | -2.10E+00 | 5.98E-01 | -3.52E+00 | 4.33E-04 | 2.75E-02 |
| Pdia5 | 6.23E+02 | -2.00E+00 | 5.09E-01 | -3.94E+00 | 8.16E-05 | 1.27E-02 |
| Nt5dc2 | 2.36E+03 | -1.91E+00 | 5.94E-01 | -3.22E+00 | 1.29E-03 | 4.82E-02 |
| Slc7a5 | 1.76E+03 | -1.91E+00 | 4.77E-01 | -4.01E+00 | 6.12E-05 | 1.02E-02 |
| Zfp970 | 5.07E+02 | -1.89E+00 | 5.54E-01 | -3.42E+00 | 6.35E-04 | 3.35E-02 |
| Dennd5b | 6.50E+02 | -1.88E+00 | 5.70E-01 | -3.30E+00 | 9.62E-04 | 4.17E-02 |
| Csrp1 | 2.53E+04 | -1.86E+00 | 5.00E-01 | -3.73E+00 | 1.93E-04 | 1.93E-02 |
| Spry1 | 7.19E+02 | -1.82E+00 | 4.76E-01 | -3.83E+00 | 1.29E-04 | 1.56E-02 |

**Supplementary Table 2C. Top 50 DEGs identified in comparison between bleomycin vs PBS challenge in *Ephb4* control mice**

| geneSymbol | baseMean | log2FoldChange | lfcSE | stat | pvalue | padj |
|---|---|---|---|---|---|---|
| Disp1 | 918.03 | 1.31 | 0.21 | 6.13 | 8.91E-10 | 1.10E-05 |
| Marco | 37247.68 | -5.56 | 0.92 | -6.04 | 1.57E-09 | 1.10E-05 |
| Ralgapa2 | 3567.91 | -1.08 | 0.19 | -5.78 | 7.50E-09 | 3.83E-05 |
| Cemip | 6761.34 | 6.67 | 1.25 | 5.33 | 9.72E-08 | 2.73E-04 |
| Rin2 | 10929.57 | -1.27 | 0.24 | -5.31 | 1.07E-07 | 2.73E-04 |
| Aacs | 2826.58 | 1.28 | 0.24 | 5.25 | 1.50E-07 | 3.26E-04 |
| Nars | 16204.76 | 0.58 | 0.11 | 5.24 | 1.59E-07 | 3.26E-04 |
| Usp6nl | 3491.61 | -0.67 | 0.13 | -5.09 | 3.49E-07 | 5.09E-04 |
| Aars | 18765.35 | 0.70 | 0.14 | 5.05 | 4.31E-07 | 5.38E-04 |
| Atf5 | 2300.43 | 1.59 | 0.32 | 4.96 | 7.08E-07 | 7.61E-04 |
| Eno3 | 796.68 | 2.16 | 0.44 | 4.92 | 8.68E-07 | 8.06E-04 |
| Mbd1 | 4984.71 | 1.10 | 0.22 | 4.91 | 8.95E-07 | 8.06E-04 |
| Akt1s1 | 2217.73 | 1.05 | 0.21 | 4.91 | 9.01E-07 | 8.06E-04 |
| Galnt10 | 2426.05 | 1.12 | 0.23 | 4.84 | 1.30E-06 | 1.02E-03 |
| Slc25a4 | 12978.09 | 1.00 | 0.21 | 4.85 | 1.26E-06 | 1.02E-03 |
| Lpl | 90620.39 | -3.10 | 0.66 | -4.73 | 2.29E-06 | 1.67E-03 |
| Akap1 | 1337.69 | 1.12 | 0.25 | 4.50 | 6.84E-06 | 3.88E-03 |
| Epb41l3 | 2256.18 | 4.08 | 0.91 | 4.48 | 7.35E-06 | 3.99E-03 |
| Ccdc80 | 7744.35 | 2.97 | 0.66 | 4.47 | 7.66E-06 | 3.99E-03 |
| Ext2 | 3728.73 | 0.64 | 0.14 | 4.48 | 7.56E-06 | 3.99E-03 |
| Ipo4 | 3426.73 | 0.68 | 0.15 | 4.44 | 8.93E-06 | 4.08E-03 |
| Zfp143 | 1675.10 | -0.42 | 0.09 | -4.45 | 8.42E-06 | 4.08E-03 |
| A930007I19Rik | 1757.99 | -2.65 | 0.60 | -4.44 | 8.98E-06 | 4.08E-03 |
| Chpf | 1652.09 | 1.13 | 0.25 | 4.43 | 9.50E-06 | 4.13E-03 |
| Vsig8 | 1701.89 | -4.04 | 0.92 | -4.40 | 1.08E-05 | 4.41E-03 |
| Pmp22 | 4449.79 | 2.11 | 0.48 | 4.36 | 1.32E-05 | 4.80E-03 |
| Il1rn | 36150.73 | -3.23 | 0.74 | -4.36 | 1.29E-05 | 4.80E-03 |
| Ppp1r8 | 2266.49 | 0.54 | 0.12 | 4.35 | 1.38E-05 | 4.95E-03 |
| Aldh1a2 | 3163.68 | 3.57 | 0.82 | 4.33 | 1.49E-05 | 5.15E-03 |
| Ly6i | 1278.31 | -4.94 | 1.15 | -4.31 | 1.63E-05 | 5.39E-03 |
| Wls | 15662.32 | 1.37 | 0.32 | 4.27 | 1.99E-05 | 5.97E-03 |
| Pop1 | 1141.08 | 0.93 | 0.22 | 4.27 | 1.96E-05 | 5.97E-03 |
| R3hcc1 | 668.25 | 0.80 | 0.19 | 4.27 | 1.97E-05 | 5.97E-03 |
| Arhgef2 | 8126.47 | 0.65 | 0.15 | 4.25 | 2.17E-05 | 6.35E-03 |
| Slc39a6 | 3655.31 | 1.05 | 0.25 | 4.24 | 2.23E-05 | 6.41E-03 |
| Ano8 | 939.32 | 0.65 | 0.15 | 4.24 | 2.26E-05 | 6.42E-03 |
| Mvk | 969.95 | 1.24 | 0.30 | 4.19 | 2.77E-05 | 7.02E-03 |
| Snta1 | 812.62 | 1.07 | 0.25 | 4.20 | 2.69E-05 | 7.02E-03 |
| Utp4 | 2725.18 | 0.56 | 0.13 | 4.19 | 2.82E-05 | 7.02E-03 |
| Cd200 | 16157.63 | -2.39 | 0.57 | -4.19 | 2.78E-05 | 7.02E-03 |
| Sipa1l1 | 2824.14 | 1.68 | 0.40 | 4.17 | 3.03E-05 | 7.26E-03 |
| Lgals1 | 58452.51 | 1.14 | 0.27 | 4.17 | 3.11E-05 | 7.26E-03 |
| Chd7 | 2668.65 | -0.99 | 0.24 | -4.17 | 3.05E-05 | 7.26E-03 |
| Gm13571 | 3060.38 | -4.11 | 0.99 | -4.16 | 3.13E-05 | 7.26E-03 |
| Tll1 | 666.53 | 4.50 | 1.09 | 4.11 | 3.92E-05 | 8.26E-03 |
| Timp3 | 36965.03 | 3.86 | 0.94 | 4.11 | 4.01E-05 | 8.26E-03 |
| Lrig3 | 614.49 | 2.03 | 0.49 | 4.12 | 3.82E-05 | 8.26E-03 |
| Adpgk | 2646.32 | 0.83 | 0.20 | 4.12 | 3.73E-05 | 8.26E-03 |
| Acadl | 5663.69 | -1.09 | 0.27 | -4.11 | 3.97E-05 | 8.26E-03 |
| Xxylt1 | 1144.04 | 0.76 | 0.19 | 4.09 | 4.23E-05 | 8.45E-03 |

**Supplementary Table 2D. No genes express significant changes when comparing *Ephb4* CKO and control mice after PBS challenge. Table is the expression of genes related to ECM, protein trafficking, and ER cargo concentration in comparison of Ephb4 CKO and control mice after PBS challenge.**

| geneSymbol | baseMean | log2FoldChange | lfcSE | stat | pvalue | padj | Gene_group |
|---|---|---|---|---|---|---|---|
| **Col11a1** | 2.92E+03 | -1.47E+00 | 1.62E+00 | -9.11E-01 | 3.62E-01 | 1.00E+00 | ECM |
| **Eln** | 1.30E+03 | -2.43E+00 | 2.03E+00 | -1.20E+00 | 2.31E-01 | 1.00E+00 | ECM |
| **Itga5** | 9.27E+03 | 1.57E-01 | 3.64E-01 | 4.31E-01 | 6.67E-01 | 1.00E+00 | ECM |
| **Itgb1** | 6.63E+04 | -1.50E-01 | 2.77E-01 | -5.41E-01 | 5.88E-01 | 1.00E+00 | ECM |
| **Lamc1** | 6.49E+03 | -9.96E-02 | 4.19E-01 | -2.37E-01 | 8.12E-01 | 1.00E+00 | ECM |
| **Lox** | 2.43E+04 | -7.89E-01 | 7.73E-01 | -1.02E+00 | 3.08E-01 | 1.00E+00 | ECM |
| **Loxl2** | 1.13E+04 | -7.75E-01 | 1.18E+00 | -6.55E-01 | 5.12E-01 | 1.00E+00 | ECM |
| **P3h1** | 3.21E+03 | -4.23E-01 | 4.87E-01 | -8.70E-01 | 3.84E-01 | 1.00E+00 | ECM |
| **P4hb** | 6.91E+04 | -2.03E-01 | 2.54E-01 | -8.01E-01 | 4.23E-01 | 1.00E+00 | ECM |
| **Pxdn** | 5.07E+03 | -7.72E-01 | 8.24E-01 | -9.37E-01 | 3.49E-01 | 1.00E+00 | ECM |
| **Tll1** | 6.67E+02 | 3.65E-01 | 1.04E+00 | 3.50E-01 | 7.26E-01 | 1.00E+00 | ECM |
| **Cnih1** | 4.79E+03 | -4.24E-02 | 1.44E-01 | -2.95E-01 | 7.68E-01 | 1.00E+00 | ER Cargo |
| **Lman1** | 1.19E+04 | -3.99E-01 | 2.17E-01 | -1.84E+00 | 6.58E-02 | 1.00E+00 | ER Cargo |
| **Sec23a** | 8.04E+03 | -7.46E-01 | 3.98E-01 | -1.87E+00 | 6.11E-02 | 1.00E+00 | ER Cargo |
| **Sec24d** | 6.86E+03 | -6.17E-01 | 3.41E-01 | -1.81E+00 | 7.01E-02 | 1.00E+00 | ER Cargo |
| **Arcn1** | 1.80E+04 | -2.88E-01 | 1.14E-01 | -2.52E+00 | 1.18E-02 | 1.00E+00 | Protein Transport |
| **Arf4** | 1.64E+04 | -3.27E-01 | 1.65E-01 | -1.97E+00 | 4.83E-02 | 1.00E+00 | Protein Transport |
| **Copb2** | 1.92E+04 | -2.83E-01 | 1.52E-01 | -1.86E+00 | 6.33E-02 | 1.00E+00 | Protein Transport |
| **Copg1** | 1.26E+04 | -2.24E-01 | 1.81E-01 | -1.24E+00 | 2.15E-01 | 1.00E+00 | Protein Transport |
| **Kdelr2** | 8.71E+03 | -5.41E-01 | 2.84E-01 | -1.91E+00 | 5.66E-02 | 1.00E+00 | Protein Transport |
| **Tuba1a** | 2.30E+04 | -3.31E-01 | 4.63E-01 | -7.15E-01 | 4.75E-01 | 1.00E+00 | Protein Transport |
| **Tubb6** | 1.55E+04 | 2.01E-01 | 3.72E-01 | 5.39E-01 | 5.90E-01 | 1.00E+00 | Protein Transport |

**Supplementary Table 3. Pathway enrichment analysis of DEGs from RNA sequencing of lung fibroblasts from bleomycin-challenged *EphB4* CKO versus control mice using pathDIP ver. 5 (14)(https://ophid.utoronto.ca/pathDIP).**

| 8 pathways included exclusively in "Upregulated DEGs" |
|---|
| Cytokine Signaling in Immune system |
| Signaling by Interleukins |
| Metabolism of lipids |
| Metabolism of vitamins and cofactors |
| InlB-mediated entry of Listeria monocytogenes into host cell |
| Metabolism of water-soluble vitamins and cofactors |
| Phospholipid metabolism |
| Effects of PIP2 hydrolysis |

| 11 pathways included exclusively in "Downregulated" DEGs |
|---|
| Post-translational protein modification |
| Crosslinking of collagen fibrils |
| COPI-mediated anterograde transport |
| Collagen formation |
| COPI-dependent Golgi-to-ER retrograde traffic |
| Class I MHC mediated antigen processing & presentation |
| Elastic fibre formation |
| Extracellular matrix organization |
| Translation |
| Cargo concentration in the ER |
| Fibronectin matrix formation |

| Pathway Category | Number of Occurrences |
|---|---|
| Protein Trafficking | **7** |
| ECM Related | **5** |
| Lipid Synthesis | **3** |
| Translation | **3** |
| MHC Presentation | **2** |

| 10 common pathways in "Downregulated DEGs" and "All DEGs" |
|---|
| Cytosolic tRNA aminoacylation |
| tRNA Aminoacylation |
| Metabolism of proteins |
| ER to Golgi Anterograde Transport |
| Transport to the Golgi and subsequent modification |
| Asparagine N-linked glycosylation |
| COPII-mediated vesicle transport |
| Membrane Trafficking |
| Antigen Presentation: Folding, assembly and peptide loading of class I MHC |
| Vesicle-mediated transport |

| 2 common pathways in "Upregulated DEGs" and "All DEGs" |
|---|
| Metabolism |
| MET interacts with TNS proteins |

#### Supplementary Table 5. Expression data of NanoString mouse targets after housekeeping gene normalization

| Mouse ID | 1 | 2 | 3 | 4 | 5 | 6 | 7 | 8 | 9 |
|---|---|---|---|---|---|---|---|---|---|
| **Genes** | **Vehicle and PBS** | | | | | | | | |
| **Arcn1** | 4701.71 | 6987.46 | 5932.61 | 6348.86 | 5614.27 | 5880.49 | 6128.28 | 6453.05 | 6495.28 |
| **Arf4** | 8122.53 | 11923.59 | 10599.98 | 10215.55 | 10047.65 | 8763.44 | 10695.54 | 9719.66 | 6123.83 |
| **Axin2** | 128.44 | 60.33 | 37.21 | 44.25 | 72.88 | 42.64 | 37.67 | 65.13 | 7.63 |
| **Calu** | 15113.83 | 17449.65 | 14633.3 | 15463.92 | 13394.62 | 13938.35 | 15681.3 | 15052.95 | 5309.7 |
| **Cdc16** | 2071.58 | 2174.22 | 2155.49 | 2308.03 | 2140.44 | 2323.28 | 2163.66 | 2402.36 | 1895.41 |
| **Cdc23** | 1359.84 | 1825.63 | 1384.04 | 1617.19 | 1512.88 | 1431.15 | 1459.18 | 1702.19 | 1399.3 |
| **Cnih1** | 6792.15 | 7861.17 | 6925.91 | 7179.58 | 6370.04 | 5991.09 | 6887.16 | 7130.67 | 6764.96 |
| **Col11a1** | 1662.69 | 1765.3 | 1983.74 | 1420.21 | 1564.17 | 1720.97 | 1501.03 | 1613.26 | 2284.67 |
| **Copb2** | 4587.41 | 6308.15 | 5758 | 5442.49 | 5422.63 | 5298.17 | 5427.98 | 5561.25 | 5136.69 |
| **Copg1** | 3314.77 | 4366.32 | 3920.25 | 3713.97 | 3581.8 | 3744.43 | 4046.92 | 4304.96 | 3025.03 |
| **Cops3** | 2433.34 | 2786.49 | 2391.65 | 2596.35 | 2525.07 | 2363.92 | 2388.26 | 2518.84 | 2528.91 |
| **Dda1** | 1562.52 | 1801.05 | 2062.46 | 2123.9 | 1610.05 | 1724.97 | 1902.79 | 2205.71 | 1480.71 |
| **Eln** | 24.75 | 42.46 | 120.23 | 69.94 | 68.83 | 175.23 | 104.63 | 87.68 | 198.45 |
| **Fbxw8** | 189.72 | 239.1 | 229 | 246.93 | 245.62 | 255.18 | 246.92 | 234.22 | 89.05 |
| **Foxk2** | 214.46 | 397.75 | 246.18 | 308.31 | 225.38 | 197.22 | 265.05 | 280.57 | 129.75 |
| **Itga5** | 499.63 | 4341.74 | 1152.17 | 1347.42 | 600.56 | 658.27 | 1569.39 | 1531.85 | 618.23 |
| **Itgb1** | 36137.2 | 75928 | 45626 | 45110.02 | 35056.79 | 34448.12 | 41562.91 | 41626.7 | 28932.37 |
| **Kdelr2** | 12826.6 | 20195.92 | 16805.96 | 15492.47 | 13162.49 | 13421.99 | 14449.51 | 15224.54 | 14239.75 |
| **Klhl42** | 187.36 | 270.38 | 154.58 | 194.12 | 237.53 | 169.23 | 203.67 | 189.13 | 142.47 |
| **Lamc1** | 1971.42 | 2366.4 | 2539.07 | 2572.09 | 2643.83 | 2564.47 | 2877.9 | 2960.99 | 2783.33 |
| **Lgals1** | 57411.57 | 54462.85 | 65244.4 | 66568.82 | 55960.49 | 60013.54 | 80762.67 | 75870.96 | 49107.68 |
| **Lman1** | 3324.19 | 4180.86 | 3741.34 | 3632.61 | 3409.05 | 3525.9 | 3607.49 | 3976.79 | 3622.91 |
| **Lox** | 46568.17 | 20575.8 | 58431.57 | 30455.4 | 21999.57 | 34564.05 | 34207.02 | 34185.39 | 40716.99 |
| **Loxl2** | 2823.38 | 11043.18 | 5461.72 | 6415.94 | 4581.84 | 4010.94 | 5397.29 | 6235.11 | 2241.42 |
| **Mbd1** | 719.99 | 1888.2 | 887.39 | 3334.29 | 2743.7 | 2898.94 | 1498.24 | 1664.62 | 2569.62 |
| **Mbtps1** | 2280.15 | 2031.21 | 1895 | 2215.25 | 2363.12 | 2300.63 | 2031.13 | 2139.33 | 1732.58 |
| **P3h1** | 592.72 | 569.81 | 797.22 | 665.15 | 635.65 | 640.95 | 869.09 | 847.97 | 664.03 |
| **P4hb** | 19245.21 | 19581.42 | 18961.45 | 20924.97 | 18861.78 | 21345.93 | 21519.41 | 23173.12 | 20106.62 |
| **Pmm1** | 780.08 | 875.95 | 865.92 | 926.35 | 977.1 | 975.42 | 813.29 | 898.07 | 1083.82 |
| **Pomgnt1** | 361.76 | 612.27 | 446.56 | 431.06 | 363.04 | 360.45 | 527.31 | 429.62 | 221.34 |
| **Pxdn** | 1168.95 | 2046.85 | 1087.76 | 1530.12 | 1166.04 | 1215.28 | 1372.69 | 1360.25 | 407.07 |
| **Rab23** | 841.36 | 1812.23 | 857.33 | 1120.47 | 1056.72 | 983.41 | 1177.39 | 1313.91 | 798.87 |
| **Rad23b** | 3060.24 | 5354 | 3583.9 | 3497.01 | 3270.04 | 3033.52 | 3957.64 | 3741.32 | 2933.43 |
| **Sec13** | 1797.02 | 2788.73 | 2619.22 | 2208.11 | 2104 | 1848.23 | 2901.62 | 2459.98 | 1539.23 |
| **Sec23a** | 2888.2 | 4131.69 | 3596.78 | 3252.93 | 2894.86 | 2816.99 | 3719.09 | 3716.27 | 3114.07 |
| **Sec24d** | 1286.78 | 2397.68 | 1764.75 | 2018.27 | 1882.67 | 1816.92 | 2153.89 | 2300.9 | 712.37 |
| **Sec31a** | 3293.56 | 4440.06 | 4013.28 | 3892.39 | 3834.17 | 3797.07 | 4137.6 | 4278.65 | 3063.19 |
| **Skp2** | 71.88 | 189.94 | 114.5 | 139.88 | 107.97 | 110.6 | 139.5 | 135.27 | 58.52 |
| **Spsb1** | 160.26 | 741.87 | 500.94 | 737.94 | 516.89 | 510.36 | 552.42 | 655.08 | 396.89 |
| **Stt3a** | 7113.85 | 8819.79 | 7814.73 | 7924.66 | 7170.34 | 7672.09 | 7470.28 | 8141.47 | 5495.42 |
| **Tceb1** | 6527.01 | 19152.38 | 8654.88 | 9490.46 | 7218.92 | 6690.01 | 8830.41 | 9092.14 | 6446.94 |
| **Tfg** | 2539.4 | 3528.36 | 2932.67 | 3421.36 | 3337.52 | 3229.41 | 3629.81 | 3641.11 | 3348.14 |
| **Tll1** | 809.54 | 692.71 | 1235.19 | 396.8 | 446.71 | 483.71 | 398.97 | 353.21 | 483.39 |
| **Tuba1a** | 23750.13 | 70947.17 | 41400.89 | 42716.34 | 27856.76 | 29510.4 | 42196.25 | 44114.23 | 35992.46 |
| **Tubb6** | 1229.04 | 5615.44 | 2152.63 | 2643.46 | 1634.35 | 1394.5 | 2690.97 | 2646.6 | 2060.78 |
| **UBAC1** | 135.51 | 107.26 | 123.09 | 142.74 | 129.56 | 119.93 | 152.06 | 176.61 | 117.03 |
| **Uba1** | 4819.55 | 6100.34 | 5151.14 | 5365.41 | 4803.17 | 5537.36 | 5418.22 | 5874.38 | 5045.1 |
| **Genes** | **Vehicle and TGFβ1** | | | | | | | | |
| **Arcn1** | 5174.43 | 6400.55 | 6135.6 | 6505.47 | 6533.52 | 7294.82 | 6299.91 | 6742.94 | 6283.01 |
| **Arf4** | 8548.71 | 13064.05 | 11860.99 | 11392.2 | 12116.59 | 10709.18 | 11162.21 | 10807.17 | 11088.52 |
| **Axin2** | 84.33 | 28.64 | 29.3 | 36.34 | 49.48 | 48.56 | 33.33 | 34.98 | 33.65 |
| **Calu** | 15775.11 | 16639.87 | 14046.83 | 17274.09 | 16354.52 | 15280.33 | 15200.86 | 14921.92 | 14677.99 |
| **Cdc16** | 2028.55 | 2525.33 | 2327.66 | 2428.71 | 2419.87 | 2495.01 | 2266.64 | 2549.49 | 2200.4 |
| **Cdc23** | 1051.75 | 1499.58 | 1374.8 | 1465.2 | 1449.85 | 1450.19 | 1424.71 | 1574 | 1291.98 |
| **Cnih1** | 6456.91 | 8139.65 | 7178.71 | 7350.6 | 7295.27 | 7725.15 | 7246.14 | 7674.71 | 7365.43 |
| **Col11a1** | 3087.33 | 4093.91 | 4936.61 | 4231.49 | 4801.76 | 4175.89 | 3684.9 | 3953.46 | 4302.75 |
| **Copb2** | 5044.43 | 6431.79 | 6287.97 | 6164.38 | 6496.7 | 6277.66 | 5734.33 | 5902.51 | 5953.29 |
| **Copg1** | 3547.61 | 4238.4 | 4160.72 | 4293.61 | 4409.38 | 4417.36 | 4196.72 | 4327.53 | 4305.63 |
| **Cops3** | 2269.82 | 2215.53 | 2368.68 | 2400.58 | 2476.25 | 2633.28 | 2237.6 | 2189.03 | 2336.9 |
| **Dda1** | 1784.93 | 2373.03 | 2154.2 | 2167.32 | 2017.13 | 2268.37 | 2067.71 | 2437.76 | 2014.87 |
| **Eln** | 245.96 | 1166.34 | 1252.91 | 2700.65 | 3310.49 | 8505.56 | 2196.74 | 1737.23 | 4561.34 |
| **Fbxw8** | 181.54 | 187.45 | 225.03 | 212.16 | 232.44 | 194.26 | 203.22 | 202.09 | 217.25 |
| **Foxk2** | 238.93 | 433.47 | 371.53 | 365.71 | 347.5 | 266.43 | 368.81 | 384.76 | 351.83 |
| **Itga5** | 1322.3 | 5024.63 | 2212.8 | 3318.38 | 2632.74 | 2970.54 | 3644.04 | 4221.63 | 2353.25 |
| **Itgb1** | 47478.76 | 71670.03 | 49837.25 | 60847.87 | 59989.21 | 52775.61 | 52343.36 | 53691.91 | 48034.86 |
| **Kdelr2** | 17659.6 | 18731.73 | 21389.63 | 20082.58 | 19988.35 | 20798.49 | 16491.17 | 18622.77 | 17853.14 |
| **Klhl42** | 159.29 | 179.64 | 157.05 | 175.82 | 156.49 | 136.25 | 159.14 | 141.85 | 158.61 |
| **Lamc1** | 1766.19 | 2058.02 | 2048.72 | 2207.17 | 2355.43 | 2119.31 | 2398.89 | 2273.56 | 2361.9 |
| **Lgals1** | 62364.91 | 88164.1 | 72950.95 | 78521.66 | 74888.13 | 70732.4 | 88835.33 | 84655.05 | 77591.74 |
| **Lman1** | 4711.8 | 4076.98 | 4894.42 | 4454.2 | 4810.97 | 5116.83 | 4293.49 | 4630.67 | 4918.94 |

| | | | | | | | | | |
|---|---|---|---|---|---|---|---|---|---|
| **Lox** | 79975.31 | 41653.7 | 86364.89 | 58216.38 | 56861.68 | 62192.44 | 52043.36 | 52615.37 | 68993.93 |
| **Loxl2** | 4941.36 | 14078.09 | 9686.87 | 12213.88 | 8803.8 | 8656.65 | 7475.17 | 8970.84 | 7485.59 |
| **Mbd1** | 607.86 | 1205.39 | 925.91 | 1580.07 | 1415.33 | 1258.63 | 993.53 | 1069.74 | 1351.58 |
| **Mbtps1** | 1923.14 | 1623.24 | 1693.59 | 1821.53 | 1769.74 | 1744.95 | 1624.71 | 1705.17 | 1630.35 |
| **P3h1** | 941.66 | 989.31 | 1026.7 | 1020.95 | 1062.07 | 1197.93 | 1062.35 | 1055.16 | 1082.42 |
| **P4hb** | 24563.91 | 25464.22 | 23166.44 | 28592.44 | 28687.44 | 35025.23 | 27918.97 | 29939.06 | 27842.9 |
| **Pmm1** | 565.7 | 838.31 | 866.13 | 838.09 | 828.48 | 832.34 | 746.23 | 808.38 | 835.36 |
| **Pomgnt1** | 462.63 | 607.9 | 521.56 | 526.3 | 451.06 | 481.6 | 580.64 | 561.59 | 618.11 |
| **Pxdn** | 1144.28 | 2172.57 | 1454.49 | 1764.1 | 1722.56 | 1739.56 | 1782.77 | 1899.49 | 1482.31 |
| **Rab23** | 972.11 | 1634.96 | 787.61 | 1447.61 | 1380.81 | 1026.6 | 1264.5 | 1275.72 | 1169.89 |
| **Rad23b** | 3685.82 | 4815.06 | 3918.11 | 4288.92 | 4283.96 | 4281.11 | 4554.78 | 4811.39 | 4228.73 |
| **Sec13** | 2119.9 | 3310.27 | 2985.17 | 2878.82 | 2948.03 | 2331.78 | 3176.3 | 2882.75 | 3028.07 |
| **Sec23a** | 3356.71 | 4342.53 | 4355.28 | 4068.56 | 4249.44 | 4561.71 | 4061.23 | 4198.31 | 3982.64 |
| **Sec24d** | 1366.81 | 1933.05 | 1763.91 | 1880.14 | 1954.99 | 2041.06 | 1902.12 | 2019 | 1962 |
| **Sec31a** | 3560.5 | 3837.47 | 4297.85 | 3973.61 | 4331.14 | 4481.44 | 4111.77 | 4240.09 | 4325.82 |
| **Skp2** | 85.5 | 216.09 | 127.75 | 151.21 | 156.49 | 166.6 | 192.47 | 210.84 | 159.57 |
| **Spsb1** | 337.31 | 1615.43 | 1051.32 | 1343.29 | 1142.62 | 1147.34 | 1160.2 | 1175.64 | 1052.62 |
| **Stt3a** | 8741.96 | 8759.26 | 9378.62 | 9553.08 | 9917.66 | 11061.27 | 8621.39 | 9517.85 | 9311.09 |
| **Tceb1** | 7364.61 | 14195.24 | 9964.64 | 12808.17 | 12248.92 | 10708.51 | 11063.29 | 12567.72 | 9898.44 |
| **Tfg** | 2787.49 | 3630.49 | 3182.07 | 3442.63 | 3461.23 | 3631.56 | 3316.08 | 3553.16 | 3212.64 |
| **Tll1** | 1096.26 | 1871.87 | 2400.33 | 1182.71 | 930.89 | 1071.79 | 1064.5 | 1079.45 | 1148.75 |
| **Tuba1a** | 29116.43 | 60174.55 | 48197.58 | 48595.31 | 44336.6 | 44322.67 | 47805.79 | 53547.14 | 43507.17 |
| **Tubb6** | 1657.27 | 4889.26 | 3205.51 | 3436.77 | 3313.94 | 2917.93 | 3399.95 | 3790.23 | 2806.98 |
| **UBAC1** | 113.61 | 118.46 | 159.4 | 137.14 | 159.94 | 108.6 | 187.09 | 180.72 | 150.92 |
| **Uba1** | 4526.75 | 5303.2 | 4832.3 | 5294.64 | 5306.91 | 6118.47 | 5388.1 | 5683.89 | 5297.69 |
| **Genes** | **NVP-BHG712 and PBS** | | | | | | | | |
| **Arcn1** | 3638.85 | 6292.71 | 4869.9 | 5306.5 | 4655.52 | 4740.6 | 5362.05 | 5881.7 | 4812.23 |
| **Arf4** | 6760.65 | 11213.36 | 8140.14 | 9173.06 | 8121.29 | 7190.5 | 8208.34 | 8347.55 | 7116.06 |
| **Axin2** | 127.31 | 55.1 | 40.82 | 64.93 | 77.92 | 51.68 | 61.9 | 51.15 | 36.53 |
| **Calu** | 10864.59 | 16043.09 | 11847.62 | 13248.42 | 11099.76 | 11114.19 | 12624.36 | 13071.41 | 11062.53 |
| **Cdc16** | 1674.57 | 2538.85 | 1882.21 | 1941.56 | 1987.58 | 1909.74 | 1945.8 | 2075.19 | 1893.48 |
| **Cdc23** | 1039.3 | 1591.08 | 1244.06 | 1355.91 | 1392.12 | 1227.04 | 1238.03 | 1354.8 | 1228.63 |
| **Cnih1** | 6451.46 | 7966.44 | 6462.05 | 6601.29 | 6180.23 | 6076 | 5951.87 | 6585.76 | 5930.05 |
| **Col11a1** | 667.75 | 1016.64 | 995.9 | 887.39 | 840.86 | 780.24 | 768.51 | 1004.41 | 1073.98 |
| **Copb2** | 3959.74 | 6092.96 | 4765.69 | 5232.65 | 4597.37 | 4468.85 | 4722.02 | 5196.14 | 4347.08 |
| **Copg1** | 2764.54 | 3946.72 | 3191.81 | 3232.53 | 2916.83 | 3166.79 | 3334.49 | 3511.61 | 3495.93 |
| **Cops3** | 1886.33 | 2373.54 | 2015.43 | 2243.29 | 2065.5 | 1996.44 | 2126.84 | 2299.36 | 1973.84 |
| **Dda1** | 1317.31 | 2489.26 | 1711.39 | 1858.8 | 1758.47 | 1534.63 | 1830.18 | 2191.63 | 1698.65 |
| **Eln** | 23.38 | 41.33 | 17.19 | 39.47 | 34.89 | 58.35 | 35.04 | 19.59 | 37.75 |
| **Fbxw8** | 189.67 | 245.21 | 204.12 | 235.53 | 202.36 | 214.23 | 218.41 | 216.55 | 210.66 |
| **Foxk2** | 250.73 | 427.04 | 305.11 | 304.28 | 274.47 | 253.41 | 297.83 | 294.9 | 312.94 |
| **Itga5** | 281.91 | 2195.83 | 820.78 | 971.42 | 383.79 | 496.82 | 929.69 | 985.91 | 444.45 |
| **Itgb1** | 28109.2 | 58076.66 | 37625.98 | 37542.71 | 27315.58 | 26976.48 | 31311.54 | 36471.98 | 29891.35 |
| **Kdelr2** | 8960.07 | 14085.58 | 12302.06 | 11136.26 | 9609.94 | 9206.11 | 9837.63 | 11702.46 | 9539.22 |
| **Klhl42** | 137.71 | 267.25 | 157.93 | 193.52 | 203.53 | 182.56 | 176.36 | 193.7 | 165.6 |
| **Lamc1** | 1939.6 | 2565.02 | 2127.16 | 2672.35 | 2514.42 | 2419.9 | 2667.6 | 2939.22 | 2986.94 |
| **Lgals1** | 65733.19 | 84415.66 | 66477.97 | 67832.27 | 69722.37 | 64181.88 | 72756.21 | 74459.82 | 59014.36 |
| **Lman1** | 2418.97 | 3295.13 | 3031.74 | 3070.84 | 2713.3 | 2862.53 | 2986.45 | 3310.29 | 3224.39 |
| **Lox** | 24740.56 | 16054.11 | 33722.97 | 16735.58 | 11125.34 | 11199.21 | 17581.14 | 21938.04 | 17717.09 |
| **Loxl2** | 1768.11 | 8184.1 | 4089.94 | 5781.38 | 3625.1 | 3549.4 | 4497.77 | 5093.85 | 4733.08 |
| **Mbd1** | 731.41 | 2512.67 | 1241.92 | 3053.02 | 2786.57 | 3816.15 | 1823.17 | 1918.49 | 2201.55 |
| **Mbtps1** | 2144.86 | 2007.11 | 1797.34 | 2210.19 | 2099.23 | 2138.98 | 1916.6 | 2006.63 | 2031.07 |
| **P3h1** | 520.95 | 786.59 | 597.32 | 586.92 | 552.43 | 550.17 | 762.67 | 686.65 | 627.1 |
| **P4hb** | 15997.44 | 18404.24 | 17467.4 | 15806.18 | 15283.1 | 17759.53 | 17367.4 | 18592.92 | 18327.14 |
| **Pmm1** | 843.13 | 1023.53 | 864.83 | 1031.25 | 1123.47 | 1122.84 | 836.25 | 941.29 | 1274.9 |
| **Pomgnt1** | 270.22 | 577.2 | 330.89 | 393.4 | 327.97 | 329.27 | 398.27 | 350.4 | 301.98 |
| **Pxdn** | 613.19 | 1666.85 | 729.46 | 1231.14 | 871.09 | 823.58 | 907.5 | 1088.2 | 864.55 |
| **Rab23** | 636.57 | 1168.17 | 640.3 | 975.23 | 744.33 | 781.9 | 847.93 | 963.05 | 841.41 |
| **Rad23b** | 2830.8 | 4712.64 | 3410.97 | 3454.06 | 2867.98 | 2984.23 | 3753.79 | 3835.89 | 3331.54 |
| **Sec13** | 1648.59 | 2967.27 | 2217.4 | 2448.27 | 2081.79 | 1813.05 | 2530.95 | 2305.89 | 1852.08 |
| **Sec23a** | 1886.33 | 3923.3 | 2664.32 | 2779.29 | 2331.83 | 2162.32 | 2662.92 | 3173.18 | 2213.72 |
| **Sec24d** | 1057.49 | 2131.09 | 1477.19 | 1809.15 | 1447.95 | 1486.28 | 1695.86 | 1888.02 | 1726.66 |
| **Sec31a** | 2535.89 | 3988.04 | 3347.59 | 3535.54 | 2870.31 | 3200.13 | 3725.76 | 3938.18 | 3393.65 |
| **Skp2** | 66.26 | 126.74 | 98.84 | 114.58 | 76.76 | 100.03 | 93.44 | 107.73 | 68.19 |
| **Spsb1** | 133.81 | 690.16 | 554.35 | 453.24 | 380.3 | 470.14 | 416.96 | 473.37 | 331.21 |
| **Stt3a** | 5556.36 | 7248.73 | 6639.31 | 6257.54 | 5383.57 | 6068.5 | 5996.25 | 6819.73 | 6201.59 |
| **Tceb1** | 5183.51 | 11436.52 | 7519.18 | 7678.38 | 5585.93 | 5949.3 | 6612.93 | 7568.4 | 6057.91 |
| **Tfg** | 2170.84 | 3833.76 | 2959.76 | 3250.36 | 3269.22 | 3164.29 | 3399.9 | 3517.05 | 3091.66 |
| **Tll1** | 455.99 | 479.39 | 1046.39 | 304.28 | 307.03 | 511.82 | 259.28 | 264.43 | 312.94 |
| **Tuba1a** | 17406.99 | 59693.92 | 34986.37 | 40831.26 | 31412.87 | 32365.61 | 41323.2 | 44968.61 | 27412.18 |
| **Tubb6** | 700.23 | 3078.85 | 1443.89 | 1554.52 | 1018.8 | 956.96 | 1400.37 | 1599.65 | 672.15 |
| **UBAC1** | 105.23 | 185.97 | 129.99 | 138.77 | 184.92 | 161.72 | 164.68 | 162.14 | 135.16 |
| **Uba1** | 4337.78 | 5513.01 | 4695.86 | 4826.52 | 4136.82 | 4909.82 | 5007 | 5024.2 | 4882.85 |
| **Genes** | **NVP-BHG712 and TGFβ1** | | | | | | | | |

| | | | | | | | | | |
|---|---|---|---|---|---|---|---|---|---|
| **Arcn1** | 4659.98 | 6568.48 | 5544.66 | 6096.87 | 5772.66 | 6051.22 | 5735.22 | 5987.01 | 5832.21 |
| **Arf4** | 9077.15 | 11523.99 | 10007.26 | 10987.73 | 10114.17 | 10080.82 | 9384.67 | 9065.79 | 9567.84 |
| **Axin2** | 125.83 | 39.31 | 50.84 | 38.53 | 55.95 | 37.82 | 40.29 | 41.91 | 34.31 |
| **Calu** | 14306.66 | 15985.52 | 12254.9 | 16167.57 | 14088.58 | 13400.3 | 13164.43 | 12967.14 | 14136.16 |
| **Cdc16** | 2060.68 | 2490.86 | 2156.86 | 2135.9 | 2052.48 | 2153.99 | 2106.34 | 2225.03 | 2022.23 |
| **Cdc23** | 1022.06 | 1505.52 | 1258.78 | 1442.34 | 1316.3 | 1264.76 | 1266.21 | 1346.91 | 1228.44 |
| **Cnih1** | 6923.75 | 7448.99 | 6793.75 | 7094.82 | 6578.53 | 6710.39 | 6115 | 6566.9 | 6503.62 |
| **Col11a1** | 1690.93 | 3355.65 | 3243.77 | 3176.23 | 3687.79 | 3209.18 | 2966.2 | 2981.32 | 3742.5 |
| **Copb2** | 5373 | 6358.83 | 5756.48 | 6154.67 | 5668.61 | 5829.05 | 5451.46 | 5497.76 | 5630.9 |
| **Copg1** | 3358.68 | 4187.68 | 3779.96 | 4120.24 | 3898.83 | 3913.52 | 3651.17 | 3937.41 | 4265.21 |
| **Cops3** | 2059.58 | 2249.77 | 2208.91 | 2290.02 | 2160.45 | 2275.31 | 2064.34 | 2207.48 | 2210.95 |
| **Dda1** | 1774.81 | 2365.07 | 1945.05 | 2058.83 | 1877.76 | 1934.44 | 1883.45 | 2128.54 | 1914.71 |
| **Eln** | 132.45 | 841.21 | 185.19 | 1514.26 | 1658.87 | 2472.8 | 1114.47 | 791.38 | 2307.03 |
| **Fbxw8** | 210.81 | 230.61 | 222.71 | 232.47 | 202.2 | 198.01 | 188.6 | 197.85 | 243.63 |
| **Foxk2** | 323.4 | 479.57 | 347.37 | 461.09 | 351.4 | 388.67 | 462.08 | 370.35 | 407.19 |
| **Itga5** | 1209.7 | 3524.68 | 1239.41 | 1988.19 | 1311.39 | 1527.38 | 2005.18 | 2379.99 | 1055.72 |
| **Itgb1** | 42534.77 | 62982.94 | 44481.97 | 52473.95 | 46558.18 | 42857 | 42834.98 | 45681.77 | 43807.33 |
| **Kdelr2** | 12954.58 | 17412.42 | 17465.51 | 16222.79 | 15867.2 | 15980.77 | 13515.92 | 14855.93 | 14644 |
| **Klhl42** | 151.21 | 220.13 | 159.77 | 188.8 | 192.39 | 164.4 | 189.46 | 158.86 | 178.43 |
| **Lamc1** | 2031.98 | 2181.63 | 2035.83 | 2657.35 | 2462.78 | 2334.67 | 2390.1 | 2481.35 | 3275.83 |
| **Lgals1** | 76651.33 | 85377.08 | 66527.48 | 78998.62 | 66567.63 | 70103.52 | 72545.83 | 74074.02 | 67520.48 |
| **Lman1** | 3842.11 | 4046.17 | 4243.52 | 4357.85 | 4258.08 | 4385.7 | 3938.36 | 4025.13 | 4972.07 |
| **Lox** | 70128.23 | 36128.58 | 68637.14 | 51627.55 | 45441.14 | 52058.05 | 41244.72 | 44445 | 64368.18 |
| **Loxl2** | 3421.59 | 11489.92 | 7452.19 | 10003.91 | 6153.51 | 6561.75 | 5213.99 | 6107.86 | 7403.78 |
| **Mbd1** | 639.06 | 1529.11 | 1096.59 | 2229.66 | 1805.12 | 1775.82 | 1183.05 | 1112.03 | 1346.25 |
| **Mbtps1** | 2142.36 | 1799.03 | 1707.82 | 1891.87 | 1703.04 | 1748.5 | 1593.69 | 1696.79 | 1858.67 |
| **P3h1** | 1068.42 | 1025.96 | 803.68 | 947.86 | 969.8 | 916.53 | 892.43 | 936.6 | 1047.72 |
| **P4hb** | 23558.2 | 24292.75 | 20144.04 | 26281.93 | 24187.05 | 26939.78 | 23891.61 | 24381.74 | 27718.72 |
| **Pmm1** | 683.21 | 960.44 | 982.81 | 932.45 | 883.42 | 850.88 | 750.12 | 854.73 | 1040.85 |
| **Pomgnt1** | 503.3 | 545.08 | 451.46 | 540.72 | 385.76 | 439.1 | 456.07 | 423.95 | 436.93 |
| **Pxdn** | 1003.3 | 2058.46 | 1143.79 | 1745.45 | 1398.75 | 1289.97 | 1431.66 | 1621.75 | 1071.74 |
| **Rab23** | 873.06 | 1371.87 | 697.17 | 1280.51 | 1053.23 | 1048.37 | 1081.03 | 1079.86 | 1228.44 |
| **Rad23b** | 3827.77 | 4722.28 | 3875.58 | 4285.92 | 3808.52 | 4110.48 | 4382.43 | 4258.06 | 4200.01 |
| **Sec13** | 2439.26 | 2967.8 | 2522.39 | 3287.97 | 2699.34 | 2769.56 | 2790.46 | 2462.83 | 2957.85 |
| **Sec23a** | 3334.39 | 4457.6 | 3588.72 | 3880.06 | 3613.19 | 3570.02 | 3554.29 | 3773.68 | 3556.06 |
| **Sec24d** | 1479.01 | 2234.04 | 1594.05 | 1967.64 | 1688.31 | 1744.3 | 1788.29 | 1940.44 | 1860.95 |
| **Sec31a** | 3602.6 | 4170.65 | 3878 | 4242.25 | 3799.69 | 3843.66 | 3870.63 | 4025.13 | 4126.81 |
| **Skp2** | 93.82 | 170.34 | 125.88 | 127.15 | 126.62 | 146.02 | 138.02 | 131.57 | 106.37 |
| **Spsb1** | 424.94 | 1396.77 | 1032.44 | 1432.07 | 1056.18 | 1099.31 | 969.59 | 945.37 | 949.35 |
| **Stt3a** | 7407.19 | 8624.32 | 8278.87 | 8598.81 | 8346.35 | 9182.14 | 7702.68 | 8362.13 | 8958.2 |
| **Tceb1** | 6424.86 | 11634.05 | 8753.33 | 10163.17 | 8838.12 | 8784.54 | 8433.95 | 9767.51 | 7320.29 |
| **Tfg** | 3112.54 | 3887.63 | 3320.02 | 3645.02 | 3401.17 | 3399.84 | 3356.26 | 3405.28 | 3339.88 |
| **Tll1** | 803.52 | 1312.91 | 1807.07 | 765.48 | 616.43 | 692.26 | 655.82 | 736.8 | 643.96 |
| **Tuba1a** | 26649.77 | 56794.45 | 45360.69 | 47259.44 | 40777.67 | 43742.55 | 43935.74 | 45892.29 | 26245.51 |
| **Tubb6** | 1248.33 | 3377.92 | 2362.62 | 2155.16 | 2023.03 | 1996.94 | 2237.51 | 2450.16 | 937.91 |
| **UBAC1** | 149 | 154.61 | 127.09 | 208.07 | 147.24 | 119.23 | 174.03 | 161.78 | 168.14 |
| **Uba1** | 4487.8 | 5393.15 | 4773.66 | 5220.94 | 4589.86 | 5514.43 | 4953.38 | 5002.66 | 5452.47 |

#### Supplementary Table 7. Characteristics of Human Lung Fibroblast Donors

| Cell LOT Number | Sex | Age (years) | Condition |
|---|---|---|---|
| **LOT 70002722** | Male | 14 | Normal lung |
| **LOT 70014625** | Female | 29 | Normal lung |
| **LOT 70041445** | Female | 20 | Normal lung |
| **LOT 19TL148581** | Male | 29 | Normal lung |
| **LOT 20TL292907** | Male | 37 | Normal lung |
| **LOT 20TL114088** | Male | 47 | IPF |
| **LOT 8F5049** | Male | 52 | IPF |
| **LOT 8F5060** | Male | 55 | IPF |
| **LOT 8F5061** | Male | 83 | IPF |

**Supplementary Table 8A. Top 50 DEGs from bulk RNA sequencing comparing primary human lung fibroblasts from control donors to IPF patients.**

| geneSymbol | baseMean | log2FoldChange | lfcSE | stat | pvalue | padj |
|---|---|---|---|---|---|---|
| AC007952.4 | 6.34E+04 | -6.54E+00 | 2.16E-01 | -3.03E+01 | 4.64E-202 | 4.47E-198 |
| NEAT1 | 1.94E+04 | -2.52E+00 | 1.27E-01 | -1.99E+01 | 9.19E-88 | 4.43E-84 |
| SNORD3D | 3.34E+04 | 9.38E+00 | 5.51E-01 | 1.70E+01 | 4.64E-65 | 1.49E-61 |
| MT-ATP6 | 1.62E+04 | -1.84E+00 | 1.15E-01 | -1.60E+01 | 2.60E-57 | 6.27E-54 |
| MT-ND5 | 7.74E+03 | -1.63E+00 | 1.11E-01 | -1.47E+01 | 7.25E-49 | 1.40E-45 |
| MT-ND4L | 3.38E+03 | -1.92E+00 | 1.37E-01 | -1.40E+01 | 1.25E-44 | 2.01E-41 |
| SNORA53 | 1.20E+03 | -3.43E+00 | 2.60E-01 | -1.32E+01 | 1.09E-39 | 1.50E-36 |
| MT-ATP8 | 2.51E+03 | -2.01E+00 | 1.58E-01 | -1.28E+01 | 2.98E-37 | 3.59E-34 |
| MT-ND4 | 1.68E+04 | -1.80E+00 | 1.43E-01 | -1.27E+01 | 1.02E-36 | 1.10E-33 |
| MT-CYB | 1.41E+04 | -1.49E+00 | 1.25E-01 | -1.20E+01 | 4.16E-33 | 4.01E-30 |
| MTATP6P1 | 2.77E+03 | -1.66E+00 | 1.52E-01 | -1.09E+01 | 9.11E-28 | 7.99E-25 |
| MT-ND1 | 7.99E+03 | -1.41E+00 | 1.35E-01 | -1.05E+01 | 1.44E-25 | 1.16E-22 |
| TPP1 | 2.19E+03 | -1.29E+00 | 1.24E-01 | -1.04E+01 | 2.36E-25 | 1.75E-22 |
| MT-ND2 | 2.93E+03 | -1.78E+00 | 1.74E-01 | -1.02E+01 | 1.25E-24 | 8.62E-22 |
| SELENOP | 5.68E+03 | -3.00E+00 | 3.10E-01 | -9.68E+00 | 3.78E-22 | 2.43E-19 |
| AFF4 | 1.13E+04 | -6.36E-01 | 6.80E-02 | -9.34E+00 | 9.22E-21 | 5.56E-18 |
| MT-CO2 | 6.08E+04 | -1.21E+00 | 1.37E-01 | -8.81E+00 | 1.20E-18 | 6.79E-16 |
| SDCBP | 9.48E+03 | -9.95E-01 | 1.13E-01 | -8.78E+00 | 1.57E-18 | 8.41E-16 |
| LINC00632 | 4.98E+02 | -2.72E+00 | 3.12E-01 | -8.72E+00 | 2.88E-18 | 1.46E-15 |
| PDXP | 8.61E+02 | 2.62E+00 | 3.05E-01 | 8.58E+00 | 9.87E-18 | 4.76E-15 |
| RCN3 | 1.01E+04 | 1.31E+00 | 1.55E-01 | 8.45E+00 | 2.95E-17 | 1.30E-14 |
| RNU2-2P | 2.02E+04 | -1.72E+00 | 2.04E-01 | -8.45E+00 | 2.89E-17 | 1.30E-14 |
| FZD7 | 4.08E+03 | 1.30E+00 | 1.56E-01 | 8.35E+00 | 6.65E-17 | 2.79E-14 |
| WWP1 | 1.15E+03 | -9.58E-01 | 1.15E-01 | -8.34E+00 | 7.73E-17 | 3.11E-14 |
| MT-ND3 | 5.21E+03 | -1.30E+00 | 1.56E-01 | -8.29E+00 | 1.11E-16 | 4.27E-14 |
| PPDPF | 2.31E+03 | 8.73E-01 | 1.07E-01 | 8.18E+00 | 2.84E-16 | 1.05E-13 |
| BDNF | 3.77E+03 | 1.13E+00 | 1.39E-01 | 8.17E+00 | 3.07E-16 | 1.09E-13 |
| SNORD17 | 5.84E+03 | -1.25E+00 | 1.55E-01 | -8.09E+00 | 5.90E-16 | 2.03E-13 |
| MT-RNR1 | 7.89E+05 | -1.11E+00 | 1.39E-01 | -8.04E+00 | 8.64E-16 | 2.87E-13 |
| SCARNA5 | 1.77E+03 | -1.22E+00 | 1.52E-01 | -8.03E+00 | 9.37E-16 | 3.01E-13 |
| H3-2 | 2.67E+03 | 1.49E+00 | 1.89E-01 | 7.89E+00 | 3.10E-15 | 9.64E-13 |
| OTULINL | 4.65E+02 | 2.74E+00 | 3.51E-01 | 7.81E+00 | 5.77E-15 | 1.74E-12 |
| CENPX | 2.25E+03 | 9.54E-01 | 1.23E-01 | 7.78E+00 | 7.06E-15 | 2.00E-12 |
| LRBA | 4.27E+03 | -9.76E-01 | 1.25E-01 | -7.79E+00 | 6.97E-15 | 2.00E-12 |
| PAPPA | 3.02E+03 | -1.86E+00 | 2.39E-01 | -7.76E+00 | 8.23E-15 | 2.27E-12 |
| BCAP29 | 2.15E+03 | -8.95E-01 | 1.16E-01 | -7.71E+00 | 1.28E-14 | 3.43E-12 |
| C5orf51 | 2.52E+03 | -9.68E-01 | 1.26E-01 | -7.69E+00 | 1.46E-14 | 3.80E-12 |
| TPI1 | 4.76E+04 | 4.79E-01 | 6.29E-02 | 7.62E+00 | 2.57E-14 | 6.52E-12 |
| GSK3B | 4.39E+03 | -5.07E-01 | 6.66E-02 | -7.62E+00 | 2.64E-14 | 6.52E-12 |
| MT-ND6 | 8.06E+02 | -1.83E+00 | 2.42E-01 | -7.55E+00 | 4.40E-14 | 1.06E-11 |
| EHBP1L1 | 3.83E+03 | 9.09E-01 | 1.23E-01 | 7.41E+00 | 1.24E-13 | 2.92E-11 |
| CA12 | 3.50E+03 | -1.80E+00 | 2.44E-01 | -7.38E+00 | 1.60E-13 | 3.67E-11 |
| HEXB | 1.28E+04 | -7.45E-01 | 1.01E-01 | -7.37E+00 | 1.73E-13 | 3.89E-11 |
| TULP4 | 2.85E+03 | -6.78E-01 | 9.21E-02 | -7.36E+00 | 1.77E-13 | 3.89E-11 |
| RYBP | 1.61E+03 | -9.15E-01 | 1.25E-01 | -7.34E+00 | 2.18E-13 | 4.67E-11 |
| SEC63 | 2.13E+03 | -1.16E+00 | 1.58E-01 | -7.32E+00 | 2.47E-13 | 5.17E-11 |
| MICAL1 | 4.04E+03 | 7.12E-01 | 9.80E-02 | 7.26E+00 | 3.75E-13 | 7.70E-11 |
| RHOQ | 6.85E+03 | -1.16E+00 | 1.61E-01 | -7.23E+00 | 4.71E-13 | 9.30E-11 |
| SEC24B | 3.51E+03 | -5.43E-01 | 7.51E-02 | -7.23E+00 | 4.73E-13 | 9.30E-11 |
| CPD | 1.80E+04 | -1.09E+00 | 1.51E-01 | -7.21E+00 | 5.67E-13 | 1.09E-10 |
| GFPT2 | 2.16E+03 | -9.11E-01 | 1.27E-01 | -7.19E+00 | 6.49E-13 | 1.23E-10 |

**Supplementary Table 8B. Top 50 DEGs from RNA sequencing comparing primary human lung fibroblasts from IPF patients treated with NVP-BHG712 compared to vehicle.**

| geneSymbol | baseMean | log2FoldChange | lfcSE | stat | pvalue | padj |
|---|---|---|---|---|---|---|
| ABHD15 | 1449.14 | 3.79E-01 | 1.05E-01 | 3.62E+00 | 2.93E-04 | 3.66E-02 |
| AC010735.2 | 743.79 | -7.22E-01 | 1.79E-01 | -4.05E+00 | 5.21E-05 | 1.11E-02 |
| ADAMTS1 | 34534.25 | -5.63E-01 | 1.35E-01 | -4.17E+00 | 3.01E-05 | 7.42E-03 |
| ADM | 10737.52 | -9.95E-01 | 2.23E-01 | -4.45E+00 | 8.49E-06 | 2.63E-03 |
| AHRR | 2758.23 | -1.13E+00 | 3.12E-01 | -3.63E+00 | 2.87E-04 | 3.66E-02 |
| AL109918.1 | 1124.52 | -1.01E+00 | 1.67E-01 | -6.03E+00 | 1.66E-09 | 2.66E-06 |
| ANO6 | 10987.96 | 3.42E-01 | 7.33E-02 | 4.66E+00 | 3.10E-06 | 1.24E-03 |
| AOX1 | 5601.04 | -3.08E-01 | 8.41E-02 | -3.66E+00 | 2.52E-04 | 3.50E-02 |
| AP000944.5 | 1236.08 | -1.45E+00 | 4.11E-01 | -3.54E+00 | 4.07E-04 | 4.71E-02 |
| ARFGAP3 | 6331.13 | -1.81E-01 | 4.56E-02 | -3.98E+00 | 7.01E-05 | 1.37E-02 |
| BRD2 | 13272.91 | 2.50E-01 | 5.65E-02 | 4.42E+00 | 9.89E-06 | 2.97E-03 |
| CAMK2G | 3208.97 | 3.22E-01 | 6.54E-02 | 4.93E+00 | 8.42E-07 | 3.85E-04 |
| CAV1 | 53700.96 | -4.79E-01 | 1.09E-01 | -4.38E+00 | 1.17E-05 | 3.30E-03 |
| CHD3 | 8583.68 | -4.14E-01 | 8.98E-02 | -4.61E+00 | 4.07E-06 | 1.50E-03 |
| CHKA | 823.15 | 5.63E-01 | 1.35E-01 | 4.16E+00 | 3.13E-05 | 7.51E-03 |
| CHML | 2458.77 | -5.38E-01 | 1.19E-01 | -4.52E+00 | 6.17E-06 | 2.12E-03 |
| CITED2 | 16810.67 | -6.42E-01 | 9.52E-02 | -6.75E+00 | 1.52E-11 | 4.88E-08 |
| CNN2 | 22849.00 | -4.74E-01 | 1.11E-01 | -4.27E+00 | 1.92E-05 | 5.13E-03 |
| COL16A1 | 9510.50 | -5.04E-01 | 1.15E-01 | -4.40E+00 | 1.08E-05 | 3.14E-03 |
| CORIN | 872.68 | -1.48E+00 | 2.84E-01 | -5.22E+00 | 1.80E-07 | 1.33E-04 |
| CRABP2 | 1269.05 | -1.50E+00 | 3.00E-01 | -5.01E+00 | 5.53E-07 | 3.05E-04 |
| CYP2U1 | 3069.62 | 3.05E-01 | 8.62E-02 | 3.54E+00 | 3.99E-04 | 4.68E-02 |
| DPYSL3 | 16214.37 | -3.49E-01 | 5.92E-02 | -5.90E+00 | 3.71E-09 | 5.10E-06 |
| DSEL | 22244.54 | -3.74E-01 | 7.95E-02 | -4.71E+00 | 2.52E-06 | 1.10E-03 |
| ELN | 3800.96 | -7.75E-01 | 1.84E-01 | -4.21E+00 | 2.60E-05 | 6.58E-03 |
| ENDOD1 | 6649.01 | 4.15E-01 | 6.49E-02 | 6.40E+00 | 1.57E-10 | 3.78E-07 |
| ETS2 | 7968.66 | -3.33E-01 | 8.51E-02 | -3.91E+00 | 9.13E-05 | 1.72E-02 |
| F2R | 31246.90 | -3.25E-01 | 8.93E-02 | -3.64E+00 | 2.72E-04 | 3.58E-02 |
| F2RL2 | 4062.20 | -7.79E-01 | 2.13E-01 | -3.65E+00 | 2.62E-04 | 3.58E-02 |
| FAM219A | 1512.03 | 4.32E-01 | 1.15E-01 | 3.75E+00 | 1.76E-04 | 2.91E-02 |
| FAS | 4058.36 | -2.78E-01 | 7.64E-02 | -3.63E+00 | 2.80E-04 | 3.63E-02 |
| FENDRR | 2535.20 | -9.25E-01 | 1.87E-01 | -4.94E+00 | 7.89E-07 | 3.79E-04 |
| FEZ1 | 3150.73 | 2.40E-01 | 6.08E-02 | 3.96E+00 | 7.57E-05 | 1.45E-02 |
| FOXL1 | 4729.39 | -2.43E-01 | 6.55E-02 | -3.71E+00 | 2.06E-04 | 3.20E-02 |
| GALNT5 | 19937.73 | -9.55E-01 | 1.36E-01 | -7.02E+00 | 2.25E-12 | 1.08E-08 |
| GARS1 | 16971.62 | -2.54E-01 | 6.92E-02 | -3.67E+00 | 2.40E-04 | 3.44E-02 |
| GDE1 | 7968.85 | 3.00E-01 | 6.00E-02 | 5.00E+00 | 5.86E-07 | 3.05E-04 |
| GDF5 | 1821.18 | -1.23E+00 | 3.37E-01 | -3.65E+00 | 2.65E-04 | 3.58E-02 |
| GFPT2 | 1934.82 | 2.99E-01 | 7.69E-02 | 3.89E+00 | 9.89E-05 | 1.83E-02 |
| GJA1 | 6089.47 | -7.14E-01 | 1.42E-01 | -5.03E+00 | 4.87E-07 | 3.02E-04 |
| GPD1L | 1476.13 | 3.84E-01 | 9.30E-02 | 4.13E+00 | 3.69E-05 | 8.64E-03 |
| HDGF | 25251.47 | -2.84E-01 | 6.26E-02 | -4.53E+00 | 5.77E-06 | 2.05E-03 |
| HMGCS1 | 1604.19 | -8.45E-01 | 2.12E-01 | -3.99E+00 | 6.74E-05 | 1.35E-02 |
| HNRNPL | 18040.30 | -1.84E-01 | 5.01E-02 | -3.67E+00 | 2.39E-04 | 3.44E-02 |
| ICMT | 21339.85 | 2.26E-01 | 6.29E-02 | 3.59E+00 | 3.32E-04 | 4.08E-02 |
| INSIG1 | 5636.83 | -1.14E+00 | 2.80E-01 | -4.08E+00 | 4.54E-05 | 1.04E-02 |
| IRS1 | 12787.83 | -2.73E-01 | 6.45E-02 | -4.24E+00 | 2.28E-05 | 5.91E-03 |
| ITPRID2 | 26802.20 | -3.47E-01 | 4.86E-02 | -7.14E+00 | 9.41E-13 | 9.04E-09 |
| JARID2 | 1300.79 | -4.19E-01 | 9.07E-02 | -4.62E+00 | 3.93E-06 | 1.50E-03 |
| LANCL2 | 1938.28 | -3.75E-01 | 7.99E-02 | -4.69E+00 | 2.74E-06 | 1.14E-03 |
| LDLR | 13996.78 | -5.41E-01 | 1.47E-01 | -3.68E+00 | 2.36E-04 | 3.44E-02 |

#### Supplementary Table 9. Protein-protein interaction comparison of DEGs identified in lung fibroblasts from bleomycin-treated *Ephb4* CKO mice to human lung fibroblasts from IPF patients treated with EphB4 inhibitor.

| uniprot1 | uniprot2 | symbol1 | symbol2 | species where gene1 or gene2 is DE | species where protein1 or protein2 is partner | methods | Evidence type |
|---|---|---|---|---|---|---|---|
| P41250 | P41250 | GARS1 | GARS1 | human & mouse | none | Cosedimentation in solution \| molecular sieving \| x-ray crystallography | exp |
| P01130 | P01130 | LDLR | LDLR | human & mouse | none | surface plasmon resonance | exp |
| P15502 | P15502 | ELN | ELN | human & mouse | none | atomic force microscopy \| biophysical \| electron microscopy\| light microscopy\| transmission electron microscopy | exp |
| P01130 | Q9NVV0 | LDLR | TMEM38B | human & mouse | none | | ortho |

#### Supplementary Table 11. Gene Ontology enrichment analysis of human DEGs.

**GO analysis biological functions associated with differentially-expressed genes upregulated in primary human IPF fibroblasts treated with EphB4 inhibitor compared to DMSO vehicle**

| Name | P-value | Adjusted p-value | Odds Ratio | Combined score |
|---|---|---|---|---|
| chloride channel complex (GO:0034707) | 0.006235 | 0.1865 | 208.03 | 1056.3 |
| multimeric ribonuclease P complex (GO:0030681) | 0.01243 | 0.1865 | 92.44 | 405.55 |
| sarcoplasmic reticulum membrane (GO:0033017) | 0.03444 | 0.1926 | 30.78 | 103.7 |
| intercalated disc (GO:0014704) | 0.03806 | 0.1926 | 27.7 | 90.54 |
| ion channel complex (GO:0034702) | 0.04287 | 0.1926 | 24.44 | 76.97 |
| sarcoplasmic reticulum (GO:0016529) | 0.05479 | 0.1926 | 18.87 | 54.82 |
| cell-cell contact zone (GO:0044291) | 0.05716 | 0.1926 | 18.05 | 51.66 |
| late endosome (GO:0005770) | 0.02311 | 0.1926 | 9.2 | 34.67 |
| autophagosome (GO:0005776) | 0.08282 | 0.2384 | 12.2 | 30.39 |
| tertiary granule membrane (GO:0070821) | 0.08741 | 0.2384 | 11.52 | 28.07 |
| specific granule membrane (GO:0035579) | 0.1078 | 0.2695 | 9.21 | 20.51 |
| mitochondrial outer membrane (GO:0005741) | 0.1462 | 0.3102 | 6.62 | 12.72 |
| organelle outer membrane (GO:0031968) | 0.1633 | 0.3102 | 5.86 | 10.62 |
| endoplasmic reticulum membrane (GO:0005789) | 0.05779 | 0.1926 | 3.71 | 10.56 |
| endocytic vesicle membrane (GO:0030666) | 0.18 | 0.3102 | 5.26 | 9.02 |
| specific granule (GO:0042581) | 0.182 | 0.3102 | 5.19 | 8.85 |
| tertiary granule (GO:0070820) | 0.1861 | 0.3102 | 5.06 | 8.51 |
| endocytic vesicle (GO:0030139) | 0.2114 | 0.3232 | 4.39 | 6.81 |
| nuclear membrane (GO:0031965) | 0.2262 | 0.3232 | 4.06 | 6.03 |
| axon (GO:0030424) | 0.2262 | 0.3232 | 4.06 | 6.03 |
| neuron projection (GO:0043005) | 0.1525 | 0.3102 | 3.05 | 5.73 |
| early endosome (GO:0005769) | 0.2846 | 0.3648 | 3.1 | 3.89 |
| secretory granule membrane (GO:0030667) | 0.2918 | 0.3648 | 3.01 | 3.7 |
| bounding membrane of organelle (GO:0098588) | 0.2487 | 0.3391 | 2.18 | 3.04 |
| mitochondrial matrix (GO:0005759) | 0.3554 | 0.4265 | 2.36 | 2.44 |
| cytoplasmic vesicle membrane (GO:0030659) | 0.3811 | 0.4398 | 2.15 | 2.08 |
| mitochondrial membrane (GO:0031966) | 0.4477 | 0.4974 | 1.74 | 1.4 |
| intracellular organelle lumen (GO:0070013) | 0.6617 | 0.709 | 0.94 | 0.39 |
| intracellular membrane-bounded organelle (GO:0043231) | 0.8164 | 0.8445 | 0.71 | 0.14 |
| nucleus (GO:0005634) | 0.8445 | 0.8445 | 0.66 | 0.11 |

**GO analysis biological functions associated with differentially expressed genes downregulated in primary human IPF fibroblasts treated with EphB4 inhibitor NVP-BHg712 (250 nm) compared to DMSO vehicle**

| Name | P-value | Adjusted p-value | Odds Ratio | Combined score |
|---|---|---|---|---|
| cortical actin cytoskeleton (GO:0030864) | 0.000285 | 0.02041 | 26.39 | 215.47 |
| cortical cytoskeleton (GO:0030863) | 0.000739 | 0.02041 | 18.7 | 134.82 |
| caveola (GO:0005901) | 0.000816 | 0.02041 | 18.04 | 128.29 |
| plasma membrane raft (GO:0044853) | 0.002014 | 0.03777 | 13 | 80.72 |
| Golgi membrane (GO:0000139) | 0.003103 | 0.04655 | 4.56 | 26.33 |
| elastic fiber (GO:0071953) | 0.01516 | 0.0788 | 83.06 | 347.96 |
| MPP7-DLG1-LIN7 complex (GO:0097025) | 0.01516 | 0.0788 | 83.06 | 347.96 |
| CD95 death-inducing signaling complex (GO:0031265) | 0.01516 | 0.0788 | 83.06 | 347.96 |

| | | | | |
|---|---|---|---|---|
| sarcolemma (GO:0042383) | 0.011 | 0.0788 | 13.48 | 60.81 |
| basolateral plasma membrane (GO:0016323) | 0.01103 | 0.0788 | 6.92 | 31.17 |
| membrane raft (GO:0045121) | 0.01354 | 0.0788 | 6.39 | 27.51 |
| early endosome (GO:0005769) | 0.00882 | 0.0788 | 5.27 | 24.93 |
| cell-cell junction (GO:0005911) | 0.0094 | 0.0788 | 5.17 | 24.13 |
| actin cytoskeleton (GO:0015629) | 0.01576 | 0.0788 | 4.41 | 18.32 |
| bounding membrane of organelle (GO:0098588) | 0.008578 | 0.0788 | 3.27 | 15.57 |
| protein kinase complex (GO:1902911) | 0.02116 | 0.08965 | 55.37 | 213.49 |
| late endosome (GO:0005770) | 0.02002 | 0.08965 | 5.49 | 21.48 |
| nucleus (GO:0005634) | 0.02152 | 0.08965 | 1.82 | 6.99 |
| death-inducing signaling complex (GO:0031264) | 0.02415 | 0.09054 | 47.46 | 176.71 |
| lipid droplet (GO:0005811) | 0.02313 | 0.09054 | 8.98 | 33.82 |
| collagen-containing extracellular matrix (GO:0062023) | 0.02866 | 0.09844 | 3.65 | 12.97 |
| intracellular membrane-bounded organelle (GO:0043231) | 0.02888 | 0.09844 | 1.73 | 6.13 |
| integral component of plasma membrane (GO:0005887) | 0.03132 | 0.1021 | 2.22 | 7.67 |
| platelet dense tubular network (GO:0031094) | 0.03305 | 0.1033 | 33.22 | 113.25 |
| early endosome membrane (GO:0031901) | 0.0354 | 0.1062 | 7.08 | 23.66 |
| endolysosome membrane (GO:0036020) | 0.05062 | 0.146 | 20.75 | 61.91 |
| supramolecular fiber (GO:0099512) | 0.05641 | 0.1567 | 18.45 | 53.03 |
| connexin complex (GO:0005922) | 0.06216 | 0.1608 | 16.6 | 46.11 |
| adherens junction (GO:0005912) | 0.06147 | 0.1608 | 5.17 | 14.41 |
| endoplasmic reticulum membrane (GO:0005789) | 0.06565 | 0.1641 | 2.43 | 6.61 |
| gap junction (GO:0005921) | 0.07073 | 0.1658 | 14.43 | 38.23 |
| Golgi-associated vesicle membrane (GO:0030660) | 0.07073 | 0.1658 | 14.43 | 38.23 |
| endolysosome (GO:0036019) | 0.07357 | 0.1672 | 13.83 | 36.09 |
| endosome membrane (GO:0010008) | 0.0769 | 0.1696 | 3.15 | 8.08 |
| nuclear inner membrane (GO:0005637) | 0.08203 | 0.172 | 12.29 | 30.74 |
| secretory vesicle (GO:0099503) | 0.08483 | 0.172 | 11.85 | 29.24 |
| endocytic vesicle membrane (GO:0030666) | 0.08391 | 0.172 | 4.3 | 10.65 |
| exocytic vesicle (GO:0070382) | 0.08763 | 0.1729 | 11.44 | 27.86 |
| intercalated disc (GO:0014704) | 0.09041 | 0.1739 | 11.06 | 26.58 |
| neuromuscular junction (GO:0031594) | 0.1097 | 0.2008 | 8.96 | 19.81 |
| cytoplasmic vesicle membrane (GO:0030659) | 0.1098 | 0.2008 | 2.68 | 5.93 |
| focal adhesion (GO:0005925) | 0.1143 | 0.2041 | 2.63 | 5.71 |
| cell-substrate junction (GO:0030055) | 0.1189 | 0.2073 | 2.59 | 5.51 |
| cell-cell contact zone (GO:0044291) | 0.1339 | 0.2183 | 7.21 | 14.49 |
| Golgi-associated vesicle (GO:0005798) | 0.1339 | 0.2183 | 7.21 | 14.49 |
| axon (GO:0030424) | 0.1284 | 0.2183 | 3.31 | 6.8 |
| intracellular non-membrane-bounded organelle (GO:0043232) | 0.1404 | 0.224 | 1.78 | 3.49 |
| tertiary granule lumen (GO:1904724) | 0.1548 | 0.2419 | 6.14 | 11.45 |
| specific granule lumen (GO:0035580) | 0.1728 | 0.2645 | 5.43 | 9.54 |
| late endosome membrane (GO:0031902) | 0.1878 | 0.2799 | 4.94 | 8.27 |
| clathrin-coated endocytic vesicle membrane (GO:0030669) | 0.1903 | 0.2799 | 4.87 | 8.08 |
| clathrin-coated endocytic vesicle (GO:0045334) | 0.2291 | 0.3225 | 3.94 | 5.81 |
| tight junction (GO:0070160) | 0.2291 | 0.3225 | 3.94 | 5.81 |
| azurophil granule lumen (GO:0035578) | 0.2408 | 0.3225 | 3.72 | 5.29 |
| clathrin-coated vesicle membrane (GO:0030665) | 0.2408 | 0.3225 | 3.72 | 5.29 |
| neuron projection (GO:0043005) | 0.2404 | 0.3225 | 1.81 | 2.58 |
| apical junction complex (GO:0043296) | 0.2592 | 0.3295 | 3.41 | 4.6 |
| secretory granule lumen (GO:0034774) | 0.2508 | 0.3295 | 2.12 | 2.93 |
| intracellular organelle lumen (GO:0070013) | 0.2589 | 0.3295 | 1.59 | 2.15 |
| cytoskeleton (GO:0005856) | 0.2766 | 0.3458 | 1.68 | 2.15 |
| mitochondrial matrix (GO:0005759) | 0.287 | 0.3529 | 1.92 | 2.4 |
| azurophil granule (GO:0042582) | 0.3783 | 0.4561 | 2.14 | 2.08 |
| specific granule (GO:0042581) | 0.3878 | 0.4561 | 2.07 | 1.96 |
| vacuolar lumen (GO:0005775) | 0.3897 | 0.4561 | 2.06 | 1.94 |
| tertiary granule (GO:0070820) | 0.3953 | 0.4561 | 2.02 | 1.88 |
| endocytic vesicle (GO:0030139) | 0.4401 | 0.5002 | 1.75 | 1.44 |
| nuclear membrane (GO:0031965) | 0.4655 | 0.521 | 1.62 | 1.24 |
| lytic vacuole (GO:0000323) | 0.4896 | 0.54 | 1.51 | 1.08 |
| dendrite (GO:0030425) | 0.5641 | 0.6132 | 1.22 | 0.7 |
| endoplasmic reticulum lumen (GO:0005788) | 0.5839 | 0.6256 | 1.15 | 0.62 |
| lysosomal membrane (GO:0005765) | 0.6381 | 0.6741 | 0.99 | 0.45 |
| organelle inner membrane (GO:0019866) | 0.6557 | 0.6778 | 0.95 | 0.4 |
| nucleolus (GO:0005730) | 0.66 | 0.6778 | 0.89 | 0.37 |
| nuclear lumen (GO:0031981) | 0.6688 | 0.6778 | 0.88 | 0.35 |
| lysosome (GO:0005764) | 0.7712 | 0.7712 | 0.68 | 0.18 |

**Supplementary Table 12. Mouse genotyping primer sequences.**

| Gene Name | Forward Sequence | Reverse Sequence |
|---|---|---|
| *Cre* | GCATTACCGGTCGATGCAACGAGTGATGAG | GAGTGAACGAACCTGGTCGAAATCAGTGCG |
| *Ephb4* | GCCCTTAAAGGACCGACTTC | GCCTAACGCTGGAGAAAGTG |

**Supplementary Table 13. RT-qPCR primer sequences.**

| **Mouse** | | |
|---|---|---|
| **Gene** | **Forward Sequence** | **Reverse Sequence** |
| *Ephb4* | CTACGTCTCTAACCTCCCATCT | GCTGGTCACCCTTTCTCTTT |
| *Col1a1* | CCGTGCTTCTCAGAACATCA | GAGCAGCCATCGACTAGGAC |
| *Acta2* | CTGACAGAGGCACCACTGAA | CATCTCCAGAGTCCAGCACA |
| **Human** | | |
| **Gene** | **Forward Sequence** | **Reverse Sequence** |
| *COL1A1* | GAACGCGTGTCATCCCTTGT | GAACGAGGTAGTCTTTCAGCAACA |
| *ACTA2* | TTCAATGTCCCAGCCATGTA | GAAGGAATAGCCACGCTCAG |

#### Supplementary Table 14. Primers used for mouse NanoString CodeSet

| Gene | Accession# | Sequence |
|---|---|---|
| **Acad9** | NM_172678.3 | TGCAAAAACCGAGGTGGTCGATTCTGATGGTTCGAAAACAGACAAAATGACCGCATTCATAGTAGAAAGAGACTTCGGCGGAATCACTAATGGGAAACCT |
| **Arcn1** | NM_145985.4 | TGCTTTTCCCTTCATCAGTCTTCTTTCCAGCAGAAATGAGTCTAGTGAGTCTGTGCCATTTCTCATTAGCCCTAAATCACTGGAGAGATCCATCTTTACC |
| **Arf4** | NM_007479.3 | GTTGGTGGTCAAGATAAAATTAGGCCTCTCTGGAGGCATTACTTCCAGAATACCCAGGGTCTCATTTTTGTGGTAGATAGCAATGATCGTGAAAGAATCC |
| **Axin2** | NM_015732.4 | ACCGCGAGTGTGAGATCCACGGAAACAGCTGAAAACGGATTCAGGTCCTTCAAGAGAAGCGACCCAGTCAATCCTTATCACGTAGGTTCCGGCTATGTCT |
| **Calu** | NM_184053.3 | GTGGGCGGCCTGGTGTGGTAAAAGCCCAGTTGTGGTGTGACTTCACCTTAGCCATTGCATCAAGCTCTTGATAGCAGATACACTCTTACGTTTCTAACCC |
| **Cdc16** | NM_027276.2 | CTGGGAGATGTCACAGTCTTCCATAAAGAGCTCGATTTGTCTCCTGAGAGGGAAGATCTATGATGCTTTAGATAATCGCACCCTGGCTACCTACAGTTAC |
| **Cdc23** | NM_178347.4 | TATTATTATAGACGGGCCCACCAGCTTCGTCCCAATGATTCTCGCATGCTGGTTGCCTTAGGAGAATGTTATGAGAAACTCAATCAACTAGTGGAAGCCA |
| **Cnih1** | NM_009919.2 | CTTTTGGCATACCATATTTGGAGGTATATGAGTAGACCAGTGATGAGCGGCCCTGGCCTCTATGACCCGACGACCATCATGAATGCAGACATTCTAGCCT |
| **Col11a1** | NM_007729.2 | GTAAAGGGAGCAGATGGTGTCAGAGGTCTCAAGGGCTCTAAAGGCGAAAAGGGTGAAGATGGCTTTCCAGGATTCAAAGGTGACATGGGTCTTAAAGGTG |
| **Copb2** | NM_015827.2 | TCTGTGAACAGATTAAACTATTACGTGGGAGGAGAGATAGTCACCATCGCCCACCTGGACAGGACAATGTATCTTCTGGGCTATATTCCAAAAGACAATA |
| **Copg1** | NM_017477.2 | GCAACTGAGGCTTTCTTTGCCATGACCAAGCTCTTCCAGTCCAATGATCCCACACTCCGCCGCATGTGCTATTTGACCATCAAGGAGATGTCCTGCATCG |
| **Cops3** | NM_011991.1 | CACTTTCTGTGCTACTACTATTATGGAGGAATGATCTATACGGGGCTGAAGAACTTTGAAAGAGCGCTGTACTTTTATGAGCAGGCTATAACTACTCCTG |
| **Dda1** | NM_001294258.1 | CCCTTTCAGATACAGGCAGAAGTTTCTCCCACTCTCAGGTCTTGGTCTGAATTCCAACTGCCTTCCTATGATTGCCTTTTGCTGTGCTACTACTGTGCAT |
| **Eln** | NM_007925.3 | GTCCCTGCCTCCTGTTACCTAAAGCTACTTCCCACATCTGGGACACCCTGGAGTCAGATGGCTCCTCACACTGGGAATAGCTCCCTTGTTCTTATGGAAT |
| **Tceb1** | NM_001310470.1 | ACTAACAGCTCCACTGAAATTCCTGAATTCCCAATTGCACCTGAAATTGCACTGGAACTGCTGATGGCTGCAAACTTCCTAGATTGTTAAATAAAATAAA |
| **Fbxw8** | NM_172721.2 | ACATCTGCAGCTTGCCTTTTGCCTTAAGCGGCCACTTCTGCTCTGTTATTAAAGGTTCTACACTGATGAGCGTGTGTGCTGCTCCTGTAAGGACTCGTTT |
| **Foxk2** | NM_001080932.2 | TTTGGGTGGCTCTCCCTTAGGACACCTACCTGCCCGTTTCCGTTTAGAGCAGTTACCTGATGAATTTGACCTACCTTGTGCCATTGTAGAGGGTGGGTGG |
| **Itga5** | NM_001314041.1 | CCTCAGCAAGAACCTGAACAACTCACAAAGCAACGTGGTCTCCTTCCCACTCTCGGTGGAGGCTCAAGCCCAGGTCTCCCTTAATGGTGTCTCCAAGCCT |

| | | |
|---|---|---|
| **Itgb1** | NM_010578.1 | CTGTGATAGGTCTAATGGCTTAATTTGTGGAGGCAATGGCGTGTGCAGGTGTCGTGTTTGTGAATGCTATCCCAATTACACTGGCAGTGCATGTGACTGT |
| **Kdelr2** | NM_025841.4 | AAAGAGTCCTCGTTGTCCTGGAAACCCTTCCTCAGATGTCACACTACATGTCAGGTTCGGGAGGATGACTAGAAAGTCCTAAGGTTTCATTACCAAAACT |
| **Klhl42** | NM_001081237.1 | AGACCTTGACCCTGCTGTGTTCTGTTGTTTCTAACGCCAAGAACCCAAGCAAACTTAAATGCCTTAGGTTTCCGTTCAAGCAGAGGGAAGGCGAGGATAC |
| **Lamc1** | NM_010683.2 | GTTCTCAAGTCTTACTATTACGCAATCTCAGACTTTGCTGTGGGCGGCAGGTGTAAATGTAACGGACATGCCAGCGAGTGTGTAAAGAACGAGTTTGACA |
| **Lgals1** | NM_008495.2 | TCTCAGGAATCTCTTCGCTTCAGCTTCAATCATGGCCTGTGGTCTGGTCGCCAGCAACCTGAATCTCAAACCTGGGGAATGTCTCAAAGTTCGGGGAGAG |
| **Lman1** | NM_027400.3 | GGACATCAAGGAACACCTGCACGTGGTGAAGAGAGATATCGACAGCCTCGCACAGCGCAGCATGCCATCTAATGAAAAACCAAAATGCCCAGACCTACCA |
| **Lox** | NM_010728.2 | GATACTTGTTGCTATTCGATCCCACGCTGCTTAGCTTTTCTGTGGGCAGAAATGTCTAATGTGACAATCAGCACATCCCCATTGTGAGGTTTCACGGCAT |
| **Loxl2** | NM_033325.2 | TTCCACTTGGCGCTCTGGTCTTCCAAACTCCCACCACAATAACCCATTCAGTTCTGTTCTCTAAAAGCGCTCTAGGGCTTCTGGACCCAAAGTTCTAAGT |
| **Mbd1** | NM_013594.2 | GTGTGCTGTGAGAATTGTGGAATCCACTTTTCATGGGATGGTGTCAAAAGGCAGAGACTTAAGACGTTGTGCAAAGATTGTCGAGCACAGAGAATCGCCT |
| **Mbtps1** | NM_019709.3 | TGCTGTGGCAGATGGGATACACAGGTGCTAATGTCAGAGTTGCTGTTTTTGATACTGGGCTCAGTGAGAAGCATCCGCATTTTAAGAATGTGAAGGAGAG |
| **P3h1** | NM_019782.3 | TGCTGTTGTACGGAGTTATAGGAACCTGCGGCTGAGTAGTCATTGGTCAGTCGAACTATTTATTTAGTGAGCACCAGCTTGTTGCTGGGAATATGAGAGG |
| **P4hb** | NM_011032.2 | AAGTGACGGTCATCGGTTTCTTCAAGGACGTAGAGTCAGACTCTGCCAAGCAGTTCTTGCTGGCAGCAGAGGCTATTGATGACATACCTTTTGGAATCAC |
| **Pmm1** | NM_013872.4 | CCTGGAGGAGAGGATCGAGTTCTCGGAACTGGACAAGAAGGAGAAGATCCGGGAGAAGTTTGTGGAAGCCTTGAAGACAGAGTTTGCTGGCAAGGGGCTG |
| **Pomgnt1** | NM_026651.2 | CTCTCTGCTCTCAGCCCAAGGGGTATCTCCACAGATGATAACGGTCTTCATTGATGGCTACTATGAGGAGCCAATGGATGTGGTGGCGCTGTTTGGTCTG |
| **Pxdn** | NM_181395.2 | GGCTTATGTTGGCACTAAGGCCATTGTGGAGTGTACTTATATGATCCCTATGCTGATAGGATTACCTTCCTAGACATAGCTAGACGCAAAGCCACATGTG |
| **Rab23** | NM_008999.3 | TAGTAACAAGATTGGTGTCTTTAATGCATCGGTCAGAAGTCACTTGGGCCAGAATTCAAGTTCCCTTAATGGTGGCGATGTCATCAACCTTAGACCTAAC |
| **Rad23b** | NM_009011.4 | ACCTCACACTCACACCAGTGCATTACACTAACCTTGTTCACTGGATTGTCTGGGGTGACTTGGGCTCATATCCACAATATTTGGTATACGGTAGTAGGTT |
| **Rplp0** | NM_007475.5 | TCAGAACACTGGTCTAGGACCCGAGAAGACCTCCTTCTTCCAGGCTTTGGGCATCACCACGAAAATCTCCAGAGGCACCATTGAAATTCTGAGTGATGTG |
| **Sec13** | NM_024206.4 | TGACCGGAAAGTCATCATCTGGAAGGAGGAAAACGGCACTTGGGAGAAGACCCATGAGCACTCGGGACACGACTCCTCAGTGAACTCTGTTTGCTGGGCC |
| **Sec23a** | NM_009147.2 | TCAGGAGCTATTGGACCCTGTGTGTCTCTTAATTCAAAAGGACCTTGCGTGTCTGAAAATGAGATTGGAACAGGAGGCACTTGTCAGTGGAAAATCTGTG |

| | | |
|---|---|---|
| **Sec24d** | NM_027135.2 | TCTTATGGTTTGGAGTGGGCAGCCCACCAGAGCTGATTCAGGGAATATTTAATGTGCCATCGTTTGCACATATCAACACAGATATGACATCGCTGCCTGA |
| **Sec31a** | NM_026969.1 | ATTTACCTAGCCACAGGAACATCCGCTCAGCAGTTGGATGCGACGTTTAGTACCAATGCTTCCCTCGAGATATTCGAGCTGGACCTTTCGGACCCATCCT |
| **Skp2** | NM_013787.2 | CTTCAGCTCTTTCCGGGTACAGCACATGGACCTGTCGAACTCAGTGATAAATGTGTCGAACCTCCATAAGATTCTGTCCGAGTGCTCCAAGCTGCAGAAT |
| **Spsb1** | NM_029035.2 | CCCAGGTTGGACTTCCCTTGGGCCAACGAGTGCCAGCTTTAATGTCAGCTGCCGGTGCTCTGTGGCCTGTATTTATTCTTTAAACAGTAGCAAAGGCCAT |
| **Stt3a** | NM_008408.4 | ATGGTCTCTTCATGGGGAGGCTATGTGTTCCTGATCAACTTGATTCCTCTACATGTCCTGGTGCTAATGCTGACAGGCCGTTTTTCTCACCGGATCTACG |
| **Tfg** | NM_001252443.1 | TGATACTGTGGATGGCAGGGAAGAGAAGCCTGCCGCTTCTGACTCTTCTGGGAAACAGTCAACTCAGGTTATGGCAGCAAGTATGTCAGCTTTTGATCCT |
| **Tll1** | NM_009390.2 | CCTGTACTCCCATGCTCAGTTTGGTGATAATAACTACCCAGGACAACTGGACTGTGAATGGTTGTTGGTGTCAGAACGAGGATCTCGACTTGAATTGTCC |
| **Tuba1a** | NM_011653.2 | GGGAGGAAGAAGGAGAGGAATACTAAATTAAATGTCACAAGGTGCTGCTTCCACAGGGATGTTTATTGTGTTCCAACACAGAAAGTTGTGGTCTGATCAG |
| **Tubb6** | NM_026473.2 | CAGGACGCCACGGTCAATGATGGGGAAGAGGCATTTGAAGACGAGGATGAAGAAGAGATCAACGAATAGGGAGCCATAAGATGCTACAGTGAACGTCTGC |
| **Uba1** | NM_001136085.1 | ACTGGTGACTTCGTCTCATTCTCAGAAGTACAGGGCATGATCCAACTCAATGGATGTCAGCCCATGGAGATCAAAGTGCTGGGTCCTTATACCTTTAGTA |
| **UBAC1** | NM_001362681.1 | AGAGCCTCAAAGGCCCTTCGGCTGAACCACATGTCAGTGCCTCAGGCCATGGAGTGGCTAATTGAACACTCAGAAGATCCAGCTATTGACACACCTCTTC |

#### Supplementary Methods

##### Western blot

Protein lysates were generated from mouse lung fibroblasts using RIPA buffer with protease and phosphatase inhibitor tablets. Lysates were applied to SDS-acrylamide gels (7.5 or 10%) for electrophoresis. Electroblotting was performed on PVDF membranes and blocked with 1% TWEEN20 10 mM Tris-buffered saline (TTBS) containing 2.5% skim milk, and 2.5% bovine serum albumin (BSA). Probing was performed for EphB4 (R&D, catalog #AF446-SP) at 1:500, phospho-EphB4 (ThermoFisher, catalog #PA5-64792) at 1:500, elastin (Abcam, ab213720) at 1:300, phospho-SMAD 2/3 (Abcam, catalog #ab272332), SMAD 2/3 (Abcam, catalog #ab217553), and GAPDH (Abcam, catalog #ab9485) at 1:1000 for 2 h. Membranes were washed with TTBS 3 times for 5 min each and incubated in HRP-conjugated anti-rabbit (1:5,000; Sigma-Aldrich, catalog #SAB3700843) or anti-mouse (1:10,000; Sigma-Aldrich, catalog #A2179) secondary antibodies for 1 h at room temperature. Membranes were washed multiple times in TTBS before visualizing with chemiluminescence substrate (Clarify Western ECL Substrate, BIO-RAD, or SuperSignal West Pico, Thermo Science) and scanned using a BIO-RAD Chemidoc apparatus. Membrane signal intensity was quantified using ImageJ (National Institutes of Health, USA).

##### RNA isolation and qRT-PCR

RNA isolation was performed using the TRIzol-chloroform method. Nanodrop was used to evaluate the quality of the RNA, and cDNA was generated using the QuantiTect™ Reverse Transcription kit (catalog 205311; Qiagen) according to the manufacturer's instructions. qRT-PCR was performed using SYBR™ Green PCR Master Mix (catalog 4309155; ThermoFisher) according to the manufacturer's instructions. qRT-PCR primer sequences can be found in **Supplementary Table 13**. CT values generated from qRT-PCR were analyzed according to the ΔCT method and expressed as relative expression ($2^{-\Delta CT}$).
